# TERT accelerates BRAF mutant-induced thyroid cancer dedifferentiation and progression by regulating ribosome biogenesis

**DOI:** 10.1101/2023.01.29.526126

**Authors:** Pengcheng Yu, Ning Qu, Rui Zhu, Jiaqian Hu, Peizhen Han, Licheng Tan, Hualei Gan, Cong He, Chuantao Fang, Yubin Lei, Jian Li, Chenxi He, Fei Lan, Xiao Shi, Wenjun Wei, Yu Wang, Qinghai Ji, Fa-Xing Yu, Yu-Long Wang

**Author notes:** **Correspondence to:** (Yu-Long Wang) and (Fa-Xing Yu). These authors contributed equally: Pengcheng Yu, Ning Qu, Rui Zhu and Jiaqian Hu.

## Abstract

TERT reactivation occurs frequently in human malignancies. While BRAF activating mutation widely existed in cancers at various stages, TERT reactivation mainly occurs in advanced tumors. However, *in vivo* evidence for TERT role in cancer progression and the underlying mechanism is currently lacking. In this study, we induced TERT and/or BRAF V600E expression in mouse thyroid epithelium. TERT overexpression alone had no evident effect on tumor initiation. BRAF^VE^ expression itself induced mediocre papillary thyroid cancer (PTC). Notably, the co-expression of BRAF^VE^ and TERT resulted in aggressive poorly differentiated thyroid carcinoma (PDTC). Spatial transcriptome revealed that tumors from co-mutant mice were highly heterogeneous and dedifferentiation process significantly correlated with ribosomal pathways. Mechanistically, TERT boosted ribosomal RNA expression and protein synthesis. CX-5461, a rRNA transcription inhibitor, effectively blocked proliferation and induced redifferentiation. Thus, TERT promotes thyroid cancer progression by inducing dedifferentiation, and ribosome biogenesis inhibition represents a potential treatment strategy for TERT-reactivated cancers.

**Highlights:**

➢ TERT accelerated thyroid cancer dedifferentiation and metastasis *in vivo*
➢ TERT regulated rRNA metabolism and MTORC1/ S6K/RPS6 activities
➢ CX-5461 inhibited the progression of TERT-reactivated melanoma and thyroid cancer
➢ Inhibition of rRNA induced redifferentiation of advanced thyroid cancer with TERT activation

## Introduction

Telomerase activity is detected in more than 90% of cancers (1). TERT (telomerase reverse transcriptase) and TERC (telomeric RNA) are two main subunits of telomerase, which respectively provide reverse transcriptase activity and RNA template for telomere elongation (2). TERC is expressed in virtually all somatic cells, while TERT is silenced in most terminally differentiated cells and can only be observed in those with stemness properties (3). TERT expression can be reactivated through gene amplification or, more frequently, promoter-activating mutation in cancer (4,5). Reactivation of TERT is associated with poor prognosis in multiple malignancies, including thyroid cancer, melanoma, glioma, and pleural mesothelioma (6). Therapies targeting TERT activity are considered suitable for most cancers since TERT reactivation is a common event (7,8). However, severe adverse effects such as bone marrow failure and cardiovascular damage can occur during TERT inhibition, as stem and progenitor cells are also markedly inhibited by telomerase inhibitors (9–11). Therefore, it is important to develop novel drugs that target TERT reactivation for cancer treatment.

Besides its classical role in telomere length regulation, TERT possesses non-classical functions independent of telomerase activity. Recent studies have shown that TERT promotes tumorigenesis via crosstalk with NF-κB, Wnt pathway, FOXO3a, or MYC in multiple cancers (12–14). Moreover, TERT stimulates RNA polymerase I and III (Pol I & III) activity and induces tRNA synthesis (15,16), suggesting functions in ribosomal biogenesis. These factors associated with TERT may be potentially targeted to treat cancer and are likely to produce fewer side effects compared with direct TERT inhibition.

Thus far, most of the TERT studies have used cancer cell lines that were either immortalized by TERT activation or by Alternative Length of Telomere (ALT) (17). Several studies using *Tert*^*–/–*^ mice concluded that TERT depletion inhibits cancer progression and tissue renewal (18,19). Unfortunately, a genetically modified mouse model mimicking *TERT* promoter mutation is not available, as the promoter sequence of the *Tert* gene is not conserved in mice. Ectopic expression of TERT in tumors to some extent may mimic gene reactivation. A TERT transgenic mouse line under the control of the keratin promoter has been successfully established and revealed a role of high TERT expression in the initiation of chemical carcinogen-induced skin cancer (20). Nevertheless, it remains a challenge to investigate TERT reactivation in tumor initiation and progression *in vivo* in different tissues.

In this study, we established a conditional TERT transgenic mouse model and demonstrated that TERT-reactivation accelerated BRAF V600E-induced thyroid cancer dedifferentiation and progression. Spatial transcriptomic analysis and molecular characterization indicated that TERT participated in rRNA metabolism and promoted translation efficiency. Moreover, targeted inhibition of rRNA transcription effectively repressed tumor progression in cancers with TERT reactivation.

## Results

### TERT reactivation occurred frequently in multiple cancers and predicted a worse prognosis

TERT activation was underestimated in cancer genomics studies due to whole-exon sequencing missing *TERT* promoter region. Recently, with the identification of *TERT* promoter mutation, the *TERT* promoter status was examined in several large-scale pan-cancer sequencing studies. Three large-scale cohorts, MSK-IMPACT, MSK-MET and China Origimed2020 cohorts, which include the most abundant cancer types and samples, analyzed the *TERT* promoter region. Given that TERT reactivation mostly occurs via promoter mutation or gene amplification (19,20), we scanned both events in the above three cohorts (21), as well as TERT amplification in the TCGA cohort (22,23). We found frequent occurrence of *TERT* promoter mutation in thyroid cancer, melanoma, bladder cancer, hepatocellular carcinoma, glioma, as well as head and neck cancer, whereas TERT amplification appeared commonly in lung, ovarian, and bladder cancers (Figure 1A, supplementary figure 1A). Among different cancer types, melanoma and thyroid cancer exhibited high concordance within the cohorts. Furthermore, in MSKCC datasets, *TERT* promoter mutation predicted a worse prognosis in thyroid cancer(21), glioma(22), and bladder cancer(23) (Figure 1B, supplementary figure 1B).

**Figure 1.**
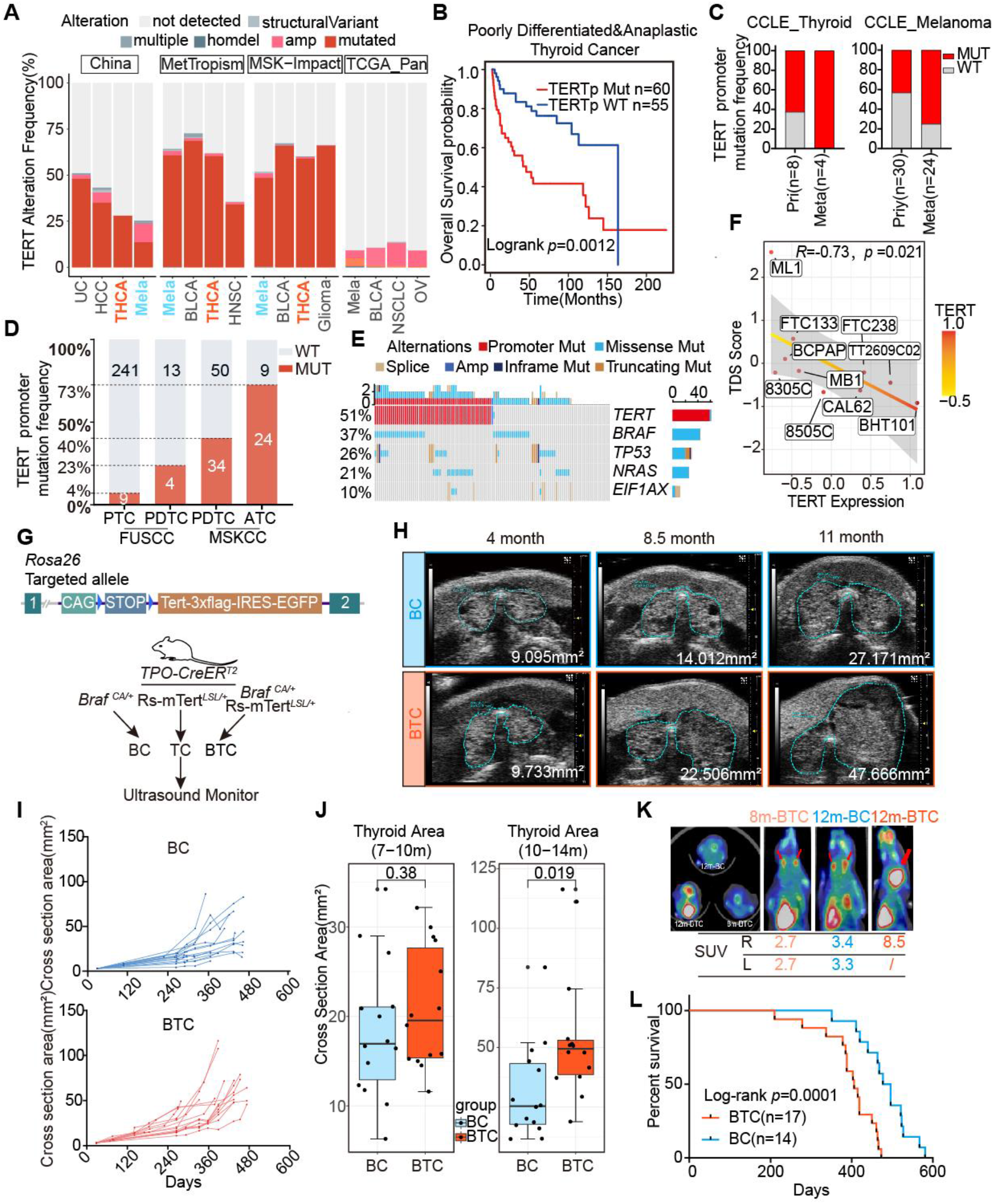
TERT reactivation predicted a worse prognosis. A. Histogram showing TERT alteration frequency in the China Origimed2020, MetTropism, MSK-Impact and TCGA datasets. UC=Urothelial Carcinoma, HCC=Hepatocellular Carcinoma, BLCA=bladder cancer, THCA=Thyroid Carcinoma, NSCLC=Non-small Cell Lung Cancer, OV=Ovarian Epithelial Tumor. B. Kaplan-Meier curve of PDTC /ATC patients with or without TERT promoter status in MSKCC cohort. C. TERT promoter mutation frequency of PTC, PDTC and ATC in FUSCC and MSKCC cohorts. D. TERT promoter mutation frequency of thyroid cancer and melanoma cell lines derived from primary and metastatic foci in CCLE dataset. E. Oncoprint showing the co-occurrence of *BRAF, TP53, NRAS, EIF1A* with *TERT* promoter mutation in PDTC and ATC from MSKCC. F. Correlation between TERT expression and TDS score of thyroid follicular cell lines in CCLE datasets (n=10). Spearman correlation *R*=-0.73. G. Top, schematic diagram of the constructing strategy of Rs-mTERT ^*LSL/+*^ mice. Bottom, experimental flow chart depicting protocol to generate BC, TC and BTC transgenic mice. Blue triangle, loxP site. H. Representative serial ultrasound images of BC and BTC thyroid tumors at 4-month, 8.5-month and 9.5-month age. I. Growth patterns of BC (n=14) and BTC (n=17) thyroid tumors. J. Cross section areas (mm^2^) comparison of BC and BTC thyroid at 7-10m old or 10-14m old. Student’s t-test. Boxplot showed the median with IQR and 1.5 IQR whiskers. K. ^18^FDG PET-CT analysis of representative 8m-BTC, 12m-BC and 12m-BTC mice. L. Overall survival of BC (n=14) and BTC (n=17) mice.

We also examined 503 cell lines with *TERT* promoter status defined by whole genome sequencing (WGS) and/or targeted sequencing in Cancer Cell Line Encyclopedia (CCLE) datasets (24). Promoter mutation was the dominant alteration for TERT activation in thyroid cancer and melanoma. In both cancer types, cell lines derived from metastasis foci showed a higher frequency of *TERT* promoter mutation (Figure 1C), suggesting a potential function of TERT activation in cancer progression and metastasis.

The frequency of *TERT* promoter mutation has been shown to be associated with the pathological stage and differentiation status of thyroid cancer. In differentiated thyroid cancer (DTC) which is known for favorable survival, *TERT* promoter mutation occurs at a frequency of 4-12% (21,25). Meanwhile, approximately 50% of poorly differentiated thyroid cancer (PDTC) and anaplastic thyroid cancer (ATC), which predict poor prognosis, harbored *TERT* promoter mutation (Figure 1D). Notably, most tumors with *TERT* promoter mutation also exhibited a mutation in *BRAF* or *NRAS* (Figure 1E) (21). TCGA program has established a thyroid differentiation score (TDS) system to evaluate thyroid cancer differentiation levels using mRNA expression levels of 16 genes (*TG, TPO, PAX8, DIO1, DIO2, DUOX1, DUOX2, FOXE1, GLIS3, NKX2-1, SLC26A4, SLC5A5, SLC5A8, THRA, THRB, TSHR*)(26). CCLE_THCA thyroid cancer cell lines expression analysis revealed that *TERT* expression level was negatively correlated with TDS score (R=-0.73, Spearman *p*=0.021) (Figure 1F). Together, these results show that *TERT* reactivation, especially promoter mutation, occurs frequently in a variety of cancers, and correlates with tumor progression and poor prognosis.

### TERT accelerated *Braf*^CA^-induced thyroid cancer progression

As mentioned above, *TERT* promoter mutation commonly co-occurs with *BRAF* or *NRAS* activating mutations (Figure 1E). We speculated that instead of initiating carcinogenesis, TERT overexpression may promote cancer progression with existing oncogenic signals. Previous studies showed that TERT induced proliferation, invasion, and many other tumor-like behaviors *in vitro*. In SV-40 immortalized thyroid cell line NTHY-ori 3-1, which harbors no *BRAF* or *TERT* promoter mutation, overexpression of BRAF-VE (V600E) and TERT collaboratively increased cell migration (supplementary figure 2A, 2B, 2C, 2D). Unexpectedly, *in vitro* upregulation of BRAF VE and TERT did not guarantee an increased proliferation rate (supplementary figure 2E). On the contrary, the knockdown of TERT in *TERT* promoter mutant cells inhibited proliferation and quickly induces senescence (supplementary figure 2F, 2G, 2H).

To elucidate the underlying mechanism between TERT reactivation and tumor progression *in vivo*, we generated a conditional *TERT* transgenic mouse model by inserting the *Loxp-stop-loxp-Tert-3xFlag-IRES-EGFP* sequence into *the Rosa26* allele (Rs-mTERT^*LSL/+*^ mice, Figure 1G). Mice were then crossed with TPO-CreER^T2^ mice to generate a thyroid-specific *Tert* transgenic mouse model (TC mice), which mimicked high TERT expression in tumors. TERT overexpression alone could not induce tumor formation, and neither could it promote epithelial proliferation (20) (Supplementary Figure 3A, 3B, 3C). We then adopted a previously used thyroid cancer mice model, which was induced by the expression of constitutively active *BRAF* (V600E mutant) in thyroid follicular cells (BC mice), and crossed with Rs-mTERT^*LSL/+*^ mice to generate BTC mice (Figure 1G).

Follow-up ultrasound indicated no significant difference in the growth patterns of BC and BTC thyroid tumors during the first 7 to 10 months after generation (Figure 1H,1I,1J). After 10-14 months, thyroid tumors from BTC mice grew much faster than those of BC mice (Figure 1H,1I,1J). Tumor progression in BTC and BC mice was also analyzed by ^18^F-FDG PET-CT. The standard uptake value (SUV) of a 12-month BTC thyroid tumor was much higher than those of 12-month BC and 8-month BTC mice (8.5 vs 3.35, 2.7). 12-month BC and 8-month BTC mice have similar FDG SUVs (Figure 1K). In line with the above results, the BTC group showed a worse prognosis with a median survival of 406 days vs 486.5 days of the BC group (Figure 1L). Together, these results indicated a more rapid tumor progression when both BRAF and TERT were activated.

### TERT promoted dedifferentiation of BRAF mutant-induced thyroid cancer

Compared with BC mice, BTC mice displayed accelerated tumor growth and worse survival (Figure 1J, 1L). We then characterized the morphological and molecular changes of tumors at different stages. The ultrasound examination, along with the gross appearance following dissection and H&E pathology, showed that most of the BTC tumors progressed into PDTC with larger sizes in 10-14 months (15/17, 88.2%). In stark contrast, the majority of BC thyroid tumors remained PTC, despite the progression of tumors from some senior BC mice (16-18 months, 5/14, 35.7%) into PDTC (Figure 2A). Immunohistochemistry revealed that tumors from BC mice contained elevated levels of cancer identification markers, including thyroglobulin (TG), TTF-1, PAX8, and CK19. All of these markers have been widely used in clinical thyroid cancer identification (27,28). However, the expression of TG and TTF-1 in BC tumors was lower than in neighboring normal follicular cells (Figure 2A). On the other hand, tumors from BTC mice expressed low levels of TG, TTF-1, and PAX8, and high levels of Ki67, indicative of hyperproliferation and dedifferentiation (Figure 2A). Tumors from BTC mice originated from thyroid follicular cells, as noted by the expression of ectopic Flag-TERT (Figure 2A).

**Figure 2.**
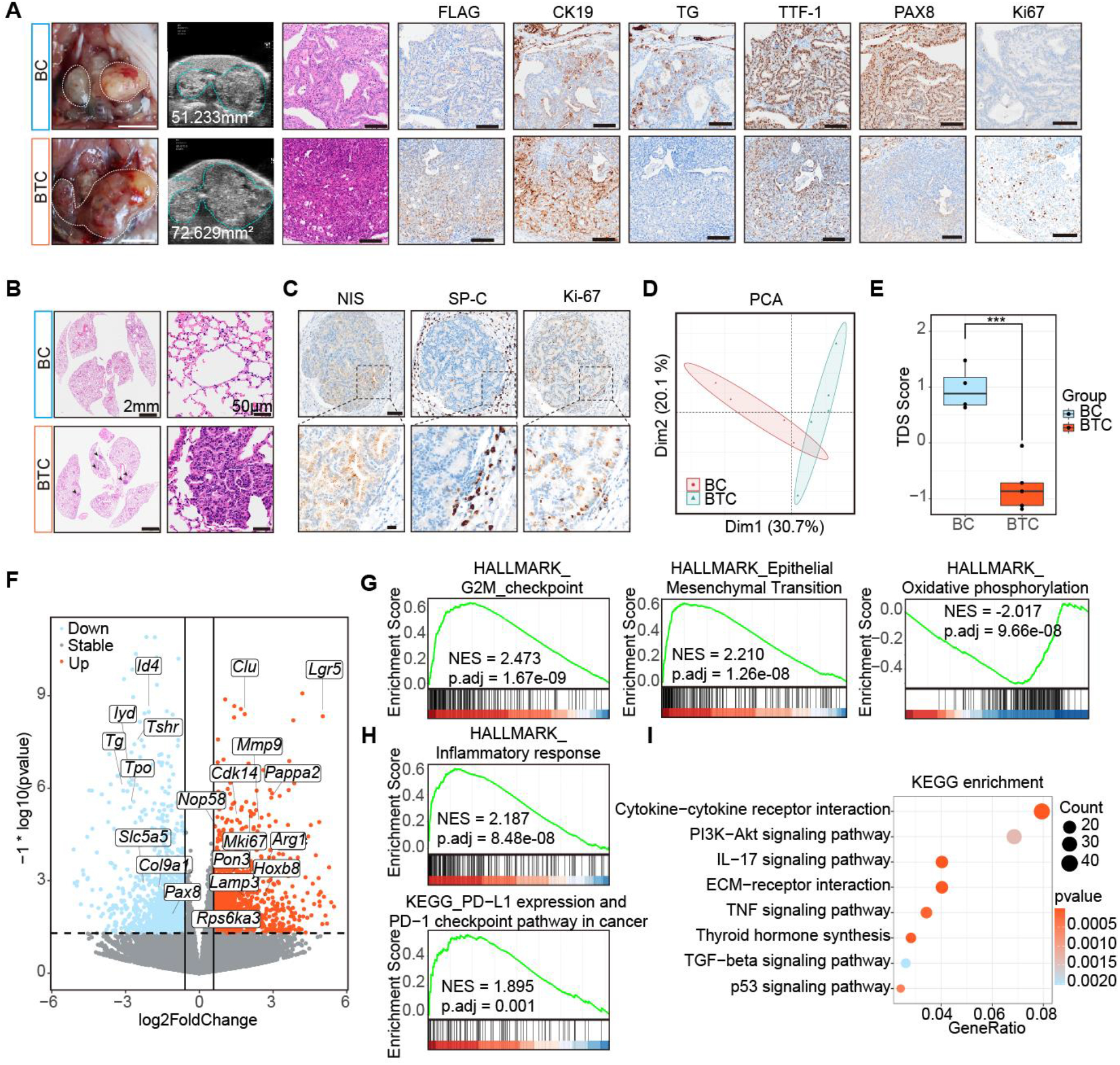
TERT promoted dedifferentiation of BRAF mutant-induced thyroid cancer. A. Photographs of the whole thyroid (white dashed line, thyroid gross appearance. scale bar, 5 mm) and ultrasound imaging (scale bar, 2 mm). H&E and Flag, CK19, TG, TTF1, PAX8, Ki-67 IHC results (scale bar, 100 μm) of BC and BTC mice thyroid. B. HE staining of BC and BTC lung sections (scale bar, left 2 mm; right 50 μm). C. NIS, SP-C and Ki-67 IHC results of BTC mice lung metastasis foci (scale bar, up 100 μm; down 20 μm). D. PCA result of BC (n=4) and BTC (n=5) transcriptome data. E. Boxplot showed TDS scores of BC and BTC. Median with IQR and 1.5 IQR whiskers. Student’s t-test, ***, *p* < 0.001. F. Volcano plot indicated upregulated and downregulated genes in BTC and BC mice. G. and H. GSEA showed BTC mice positively enriched in G2M checkpoint, epithelial-mesenchymal transition, the inflammatory response (HALLMARK) and PD-L1 and PD-1 checkpoint pathway in cancer (KEGG), and negatively enriched in oxidative phosphorylation of hallmark. NES, normalized enrichment score. I. Dot plot showed KEGG pathway enrichment of differentially expressed genes between BTC and BC mice.

Advanced thyroid cancer frequently metastasizes to distant organs, including the lung(29). Although the survival status of BTC mice became worse earlier than BC, widespread metastasis foci were found inside the BTC lung (Figure 2B). Cells in metastasis foci were negative for SP-C (a marker of AT2 lung cells) and positive for sodium iodide symporter (NIS, a marker of thyroid epithelial cells), indicating a thyroid origin of these metastasized cells (Figure 2C).

The transcriptomes of BC and BTC tumors were analyzed to identify potential mechanisms underlying cell dedifferentiation and tumor metastasis. Tumors from BC and BTC mice (n=4 and 5, respectively) were collected and subjected to RNA sequencing (RNA-seq). Principal component analysis (PCA) indicated that BTC and BC thyroid tumors showed significantly different gene expression profiles (Figure 2D). Global assessment displayed a much lower TDS score (Figure 2E, supplementary figure 4A). Thyroid-specific genes such as *Tg, Pax8, Tshr*, and *Tpo* were downregulated in those tumors. Multiple tumor-promoting genes, such as *Lgr5, Clu, Mmp9, Pappa2, Mki67*, and *Arg1*, were upregulated in tumors from BTC mice (Figure 2F). While cells from BTC thyroid could be immortalized *in vitro*, cells from BC thyroid tumors developed senescence rapidly after several passages (supplementary figure 4B). These results were in agreement with IHC analysis, indicating a hyperproliferative and dedifferentiation phenotype associated with tumors from BTC mice.

To identify key cancer pathways associated with the succession from PTC to PDTC in BTC mice, gene set enrichment analysis (GSEA) and gene set variation analysis (GSVA) for hallmark gene sets were performed(30). Interestingly, several cancer hallmarks, such as E2F targets, G2M checkpoints, epithelial-mesenchymal transition (EMT), inflammatory response, PD-L1 expression, and PD-1 checkpoint pathway were significantly activated, whereas oxidative phosphorylation and fatty acid metabolism were downregulated (Figure 2G, 2H, supplementary figure 4C). In addition, KEGG enrichment analysis of differentially expressed genes indicated that cytokine-cytokine receptor interaction, PI3K-Akt signaling pathway, IL-17 signaling pathway, ECM-receptor interaction, and TNF signaling pathway were significantly enriched in BTC mice tumors (Figure 2I).

### Spatial transcriptomic analysis of BTC mouse thyroid revealed rRNA metabolism-associated dedifferentiation

To further explore the function of TERT in dedifferentiation of *Braf* ^CA^-induced thyroid cancer, a tumor from BTC mice with profound heterogeneity was subjected to 10X spatial transcriptome sequencing. The tumor had regions with features of normal follicular cells, PTC, and PDTC, which potentially represented the whole differentiation trajectory of thyroid cancer progression (Figure 3A, supplementary figure 5A, 5B). After filtering, a total of 2,552 spots covering the thyroid, esophagus, and trachea regions were subjected to further analysis (supplementary figure 5C). Using the t-Distributed Stochastic Neighbor Embedding (T-SNE) method, all these spots are divided into 17 clusters. Combined with the spatial pathological features, the 17 clusters were further categorized into the following 8 cell types: thyroid epithelial cells (*Epcam, Tg, Tpo*), salivary gland cells (*Pigr, Msln, Agr2*), adipocyte cells (*Adipoq, Cidec, Cfd*), immune cells (*Ptprcap, Cd79b, Ccl5, Cxcl9*), muscle cells (*Actn2, Trdn, Mypz1*), erythroid cells (*Hbb-bt, Hba-a1, Hba-a2*) and esophagus epithelial cells (*Mt4, Lgals7, Krt78, Fam25c*) (supplementary figure 5E, 5F, 5G, 5H).

**Figure 3.**
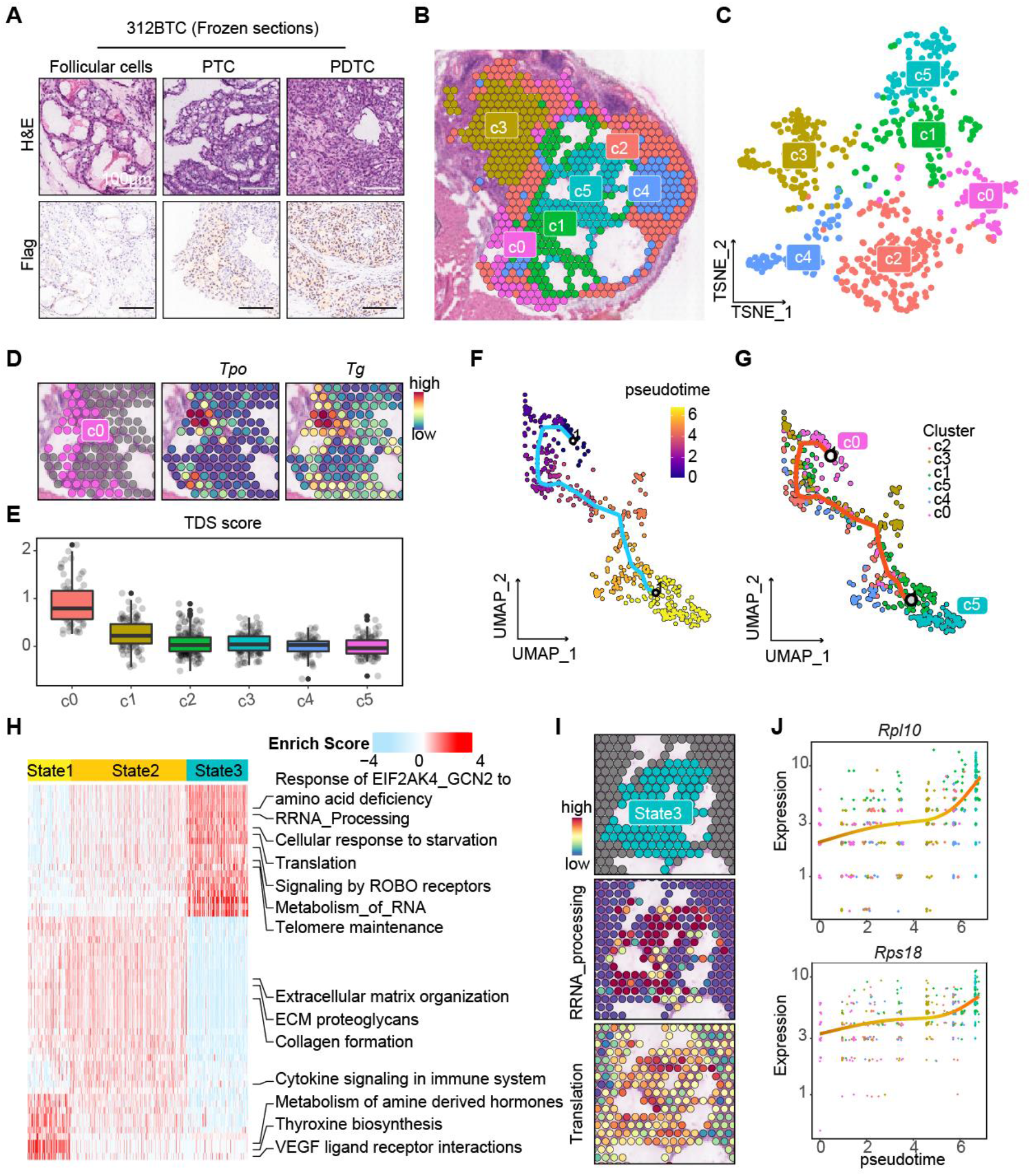
Spatial transcriptomic analysis revealed heterogeneity of BTC mice thyroid. A. H&E and anti-Flag staining results of 312BTC thyroid presented normal follicular cells, PTC, and PDTC contents. Scale bar, 100 μm. B. and C. Spatial and T-SNE projection showed different clusters of 312BTC thyroid spots. D. Spotplots showed high expression of *Tg* and *Tpo* in c0. E. Boxplot and scatter plots showed the TDS score of each thyrocyte cluster. F. and G. Monocle3 trajectory visualization of thyrocytes, colored by pseudotime and clusters. H. Heatmap presented Reactome pathways enrichment scores of state1/2/3 spots calculated by AUCell. I. Spatial feature plots showed high-enrichment of Ribosomal RNA processing and Translation in state3. J. Scatter plot showed *Rpl10* and *Rps18* gene expression level change over pseudotime, colored by clusters.

To explore the transcriptional heterogeneity of thyrocytes, we re-clustered the 637 thyroid epithelial spots (Figure 3B, 3C). We found that cells in cluster 0 (c0) express high levels of *Tg* and *Tpo*, marking well-differentiated follicular cells (Figure 3D). As several genes involved in TDS score were too low to detect, we re-defined the TDS score with *Tg, Tpo, Dio1, Sorbs2*, and *Fhl1* genes. TDS score showed that c0 was the best-differentiated cluster, followed by c1 and c2/c3/c4/c5 with poor differentiation status (Figure 3E, supplementary figure 6A).

To trace the evolutionary dynamics of the thyroid clusters, we adopted Monocle3 and indicated c0, the well-differentiated follicular cells, as the trajectory starting root (Figure 3F). In the uniform manifold approximation and projection (UMAP) projection, thyrocytes exhibited an evolution pattern of “c0-c1/c2/c3/c4-c5”, and c0 and c5 separately collected at the start and end point of the trajectory route, with the other four clusters discretion in the middle region (Figure 3G). Therefore, we could postulate that thyroid cells evolved from c0 (represented as state1), through the middle clusters c2/c3/c4 (state2) and finally dedifferentiated into c5 (state3). TDS score presented a decreasing trend from state1 to state3 (supplementary figure 6B). GSVA enrichment of hallmark genesets indicated that state 1 was enriched in KRAS signaling down pathway, and state 3 in MYC targets V1, MTORC1 signaling, and unfolded protein response pathways (supplementary figure 6C). To further identify molecular pathway alterations, Reactome enrichment scores of the three states were calculated using AUCell (Figure 3H). Consistent with the c0 normal thyroid manifestations, state1 enriched in the Thyroxine biosynthesis pathway. State2 presented an intermediate condition between state1 and state3, mainly enriched in extracellular matrix (ECM) and immune-associated pathways. Intriguingly, on the one hand, state3 was more active than state1/2 in telomere maintenance; on the other hand, state3 was significantly enriched in rRNA processing and translation pathways, which reflected the ribosome biogenesis and activities (Figure 3H, 3I). Pseudo-temporal gene expression dynamics during dedifferentiation represented multiple ribosomal proteins, such as *Rpl10* and *Rps18*, gradually increasing with pseudo-time, whereas *Actb* remained stable (Figure 3J, supplementary figure 6D). The observation confirmed the important role of the ribosome in c05/State3 cells.

Over pseudo-time, as reported in our previous study, *Tmsb4x* gradually increased(31) (supplementary figure 6E). Cell cycle regulator *Rgcc* and ferroptosis regulator *Fth1* ascended (supplementary figure 6F, 6G), while surfactant proteins *Sftpa1, Sftpb, and Sftpd* were first found to be increased (supplementary figure 6H). *Scg5*, which plays a role in regulating pituitary hormone secretion, represented a decreasing pattern same as *Tpo* and *Dio1*, indicating its involvement in thyroid differentiation (supplementary figure 6I).

### TERT induced rRNA synthesis and ribosomal activity

ERK activation was considered the upstream signaling of ribosomal RNA (rRNA) transcription, likely via phosphorylation of Upstream Binding Transcription Factor (UBF) (32). Moreover, TERT has been shown to induce rRNA and transfer RNA (tRNA) transcription (15,16). We evaluated the pre-45S rRNA levels in NTHY-CTL, NTHY-B, and NTHY-BT cells. We found that pre-45S rRNA transcription was upregulated by BRAF V600E, and further induced upon TERT expression (supplementary figure 7A). Hence, TERT and BRAF V600E may work together to enhance ribosomal biogenesis.

In cancer cells, ribosomes are frequently deregulated, resulting in uncontrolled proliferation and metastasis (33). To explore whether TERT reactivation is associated with ribosomal activities in cancer, we analyzed genes co-expressed with TERT in the datasets of CCLE. The genes which positively correlated with TERT expression (Spearman *R* > 0.25, *p* < 0.01) were involved in cell cycle progression and ribosomal RNA metabolism (Figure 4A). The ribosome consists of rRNA and ribosomal proteins, and the ribosomal biogenesis needs the participation of all of POLI/II/III, as well as snoRNA and multiple other enzymes involved in the rRNA process and maturation(34,35). We found that TERT positively correlated with the rRNA transcription regulators UBTF, POLR3E, and FBL, and the rRNA exonuclease EXOSC2 in the CCLE database (Figure 4B). The GSEA analysis also indicated that TERT high-expression group had significantly higher rRNA metabolism and ribosome biogenesis (supplementary figure 7B). This CCLE pan-cancer analysis also supported that a role of TERT in rRNA metabolism. However, we found that there’s nearly no enrichment difference in ribosome-associated pathways between BC and BTC mice. The rRNA metabolism heterogeneity was diluted and masked by various cell types in bulk tissues. Thus, spatial transcriptomic sequencing was critical to detect gene expression differences in individual cell types.

**Figure 4.**
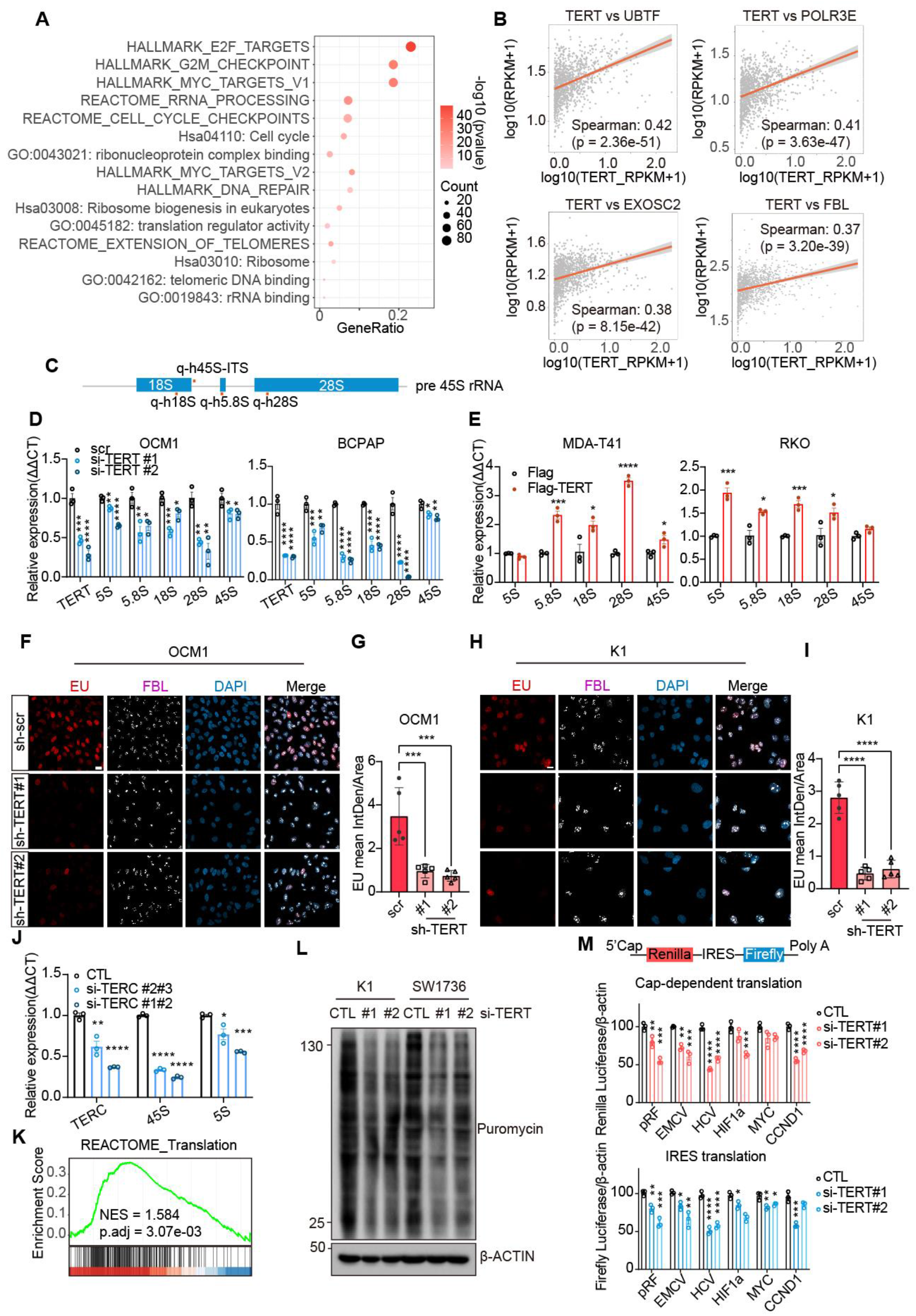
TERT induced rRNA synthesis and ribosomal activity. **A**. Enrichment results of TERT positive correlation genes (*R* > 0.25) in CCLE database (n=1156). **B**. Correlation between TERT and UBTF, POLR3E, EXOSC2, and FBL (n=1156). **C**. Schematic diagram of pre-45S rRNA and the positions of qPCR primers. **D**. and **E**. QPCR results of ribosomal RNA 5S, 5.8S, 18S, 28S, and 45S change after TERT knockdown (D, in OCM1 and BCPAP cells) and TERT over-expression (E, in MDA-T41 and RKO cells). Data represent the mean ± SEM (n=3). *, *p* < 0.05, **, *p* < 0.01, ***, *p* < 0.001, ****, *p* < 0.0001. One-way ANOVA and Dunnett multiple comparisons test in **D**. and student *t*-test in **E.** **F** and **H** EU (red) staining and FBL (white) immunofluorescence results after TERT knockdown. **G** and **I** Quantifications results of F and H EU staining results using ImageJ software (n=5). Data represent the mean ± SD.***, *p* < 0.001, ****, *p* < 0.0001. One-way ANOVA and Dunnett multiple comparisons test. **J**. QPCR results of TERC,45S, and 5S after TERC knockdown in BCPAP cells. Data represent the mean ± SEM (n=3). **K**. GSEA result of between TERT high- (n=538) and low- (n=538) expression cells in CCLE datasets. **L**. SUnSET assay results after TERT siRNA treatment of K1 and SW1736. **M**. pRF, EMCV, HCV, HIF-1a, MYC, CCND1 Bicistronic reporter assay results of HEK 293T after TERT siRNA treatment. The ratios represented the relative luciferase reads normalized by separative β-Actin protein levels. Data represent the mean ± SEM (n=3). *, *p* < 0.05, **, *p* < 0.01, ***, *p* < 0.001, ****, *p* < 0.0001. One-way ANOVA and Dunnett multiple comparisons test.

A dramatic decrease in rRNA levels was observed after TERT knockdown in K1, OCM1, and BCPAP cells (all have TERT promoter mutation) (Figure 4C, 4D, 4E, supplementary figure 7C, 7D). RRNA downregulation occurred not only in the POL I transcribed 18S, 5.8S, and 28S but also in the POL III transcribed 5S rRNA (Figure 4D, supplementary figure 7D). Nevertheless, rRNA expression increased significantly when TERT was overexpressed in thyroid cell line MDA-T41 and colon cancer cell lines RKO and HT-29 (all with wild-type TERT promoter) (Figure 4E, supplementary figure 7E). As more than 80% of cellular RNA was ribosomal RNA, an ethynyl uridine (EU) assay was performed to label nascent RNA, especially rRNA. In K1 and OCM1 cells, TERT knockdown significantly repressed nascent rRNA synthesis in the nucleus, indicating the importance of TERT in ensuring efficient rRNA synthesis. On the other hand, the protein level of Fibrillarin (FBL) was not changed (Figure 4F-4I, supplementary figure 7F). Taken together, the results indicated that TERT was required for rRNA synthesis.

As TERT is one of the main subunits of telomerase, we wondered whether telomerase activity was involved in rRNA synthesis. We found that knockdown of TERC, another main component of telomerase, in B-CPAP cells led to a reduction of rRNA expression (Figure 4J), suggesting that the regulation of rRNA synthesis by TERT was at least partially dependent on telomerase activity.

The major function of ribosomes is translation, in which protein is synthesized using mRNA as a template. As TERT could enhance rRNA transcription, we wondered whether TERT could regulate the translation process. In our spatial transcriptome data and CCLE datasets, high TERT expression was associated with positive activity in the translation pathway (Figure 3H, 4K). To validate the role of TERT in translation, we adopted two assays to examine translational efficiency. Intriguingly, the surface sensing of translation (SUnSet) assay demonstrated that TERT knockdown resulted in decreased nascent protein production in BRAF and TERT co-mutant K1 and SW1736 cells (Figure 4L). In bicistronic reporter assay, knockdown of TERT inhibited both the 5’ cap- and various internal ribosomal entry site (IRES)-dependent translation efficiencies (Figure 4M, supplementary figure 7G). Hence, the protein synthesis appeared addicted to TERT in TERT-reactivated cancer cells.

### TERT interacted with ribosomal scaffolds and enhanced MTORC1 activity

The functional telomerase complex includes multiple components besides TERT and TERC, such as Dyskerin Pseudouridine Synthase 1 (DKC1) and Pescadillo Ribosomal Biogenesis Factor 1 (PES1). DKC1 and PES1 have been linked to ribosome function (36–39). Accordingly, we asked whether TERT participates in the ribosome pathway by interacting with rRNA metabolism-associated proteins. We performed immunoprecipitation and mass spectrometry to identify potential binding partners of TERT. Several proteins involved in RNA catabolic process, ribosome biogenesis, translational control, rRNA processing, and telomere maintenance have thus been identified (Figure 5A, supplementary figure 8A, 8B). We further confirmed in co-immunoprecipitation assays that TERT interacted with EXOSC2/7/8, EIF2S1/2, and POLR1C (Figure 5B,5C). In addition, the interaction between TERT and Bystin Like (BYSL), which participates in pre-rRNA processing in yeast and human, was identified (Figure 5C) (40). Collectively, these results suggested that TERT may regulate ribosome biogenesis by interacting with known ribosome regulators.

**Figure 5.**
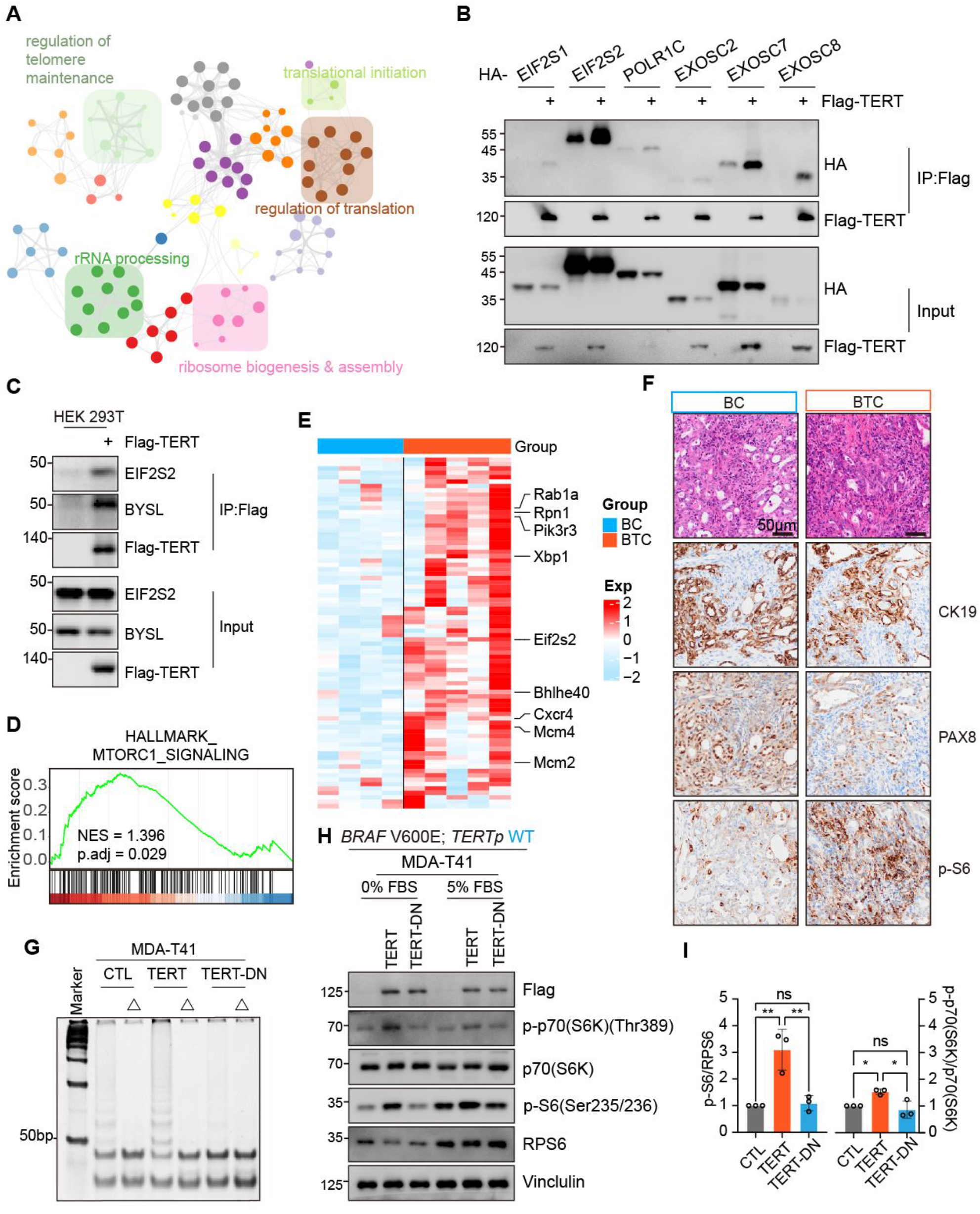
TERT interacted with ribosomal scaffolds and enhanced MTORC1 activity. **A**. TERT binding protein networks identified by IP/MS and visualized in Metascape (http://metascape.org). **B**. and **C**. Co-IP analysis of protein extracted from HEK 293T cells. **D**. GSEA showed positive enrichment of MTORC1 signaling with BTC mice thyroid (n=5), compared to BC mice thyroid (n=4). **E**. Heatmap showed that multiple genes involved in the MTORC1 pathway were highly expressed in BTC thyroid (n=5), compared to BC thyroid (n=4). **F**. H&E morphology and anti-CK19, PAX8, and p-S6 IHC staining in BC and BTC thyroid. **G**. TRAP results of MDA-T41 (CTL/ TERT/ TERT-DN) protein lysate. Δ, heat shock at 85°C, 5 min for telomerase inactivation. **H**. Western blot showed in 0% and 5% FBS media, P-P70 (S6K), P70 (S6K), P-S6, and RPS6 change in MDA-T41 (CTL/ TERT/ TERT-DN) cells. **I**. Quantification results of H. Data represent the mean ± SD (n=3). ns, not significant; *, *p* < 0.05; **, *p* < 0.01. One-way ANOVA and Dunnett multiple comparisons test.

PI3K-AKT and MTORC1 signaling pathways play a major role in regulating protein synthesis at the ribosome(41,42). In our bulk and spatial transcriptomic analysis, PI3K-AKT and MTORC1 signaling pathways were enriched in BTC tumors (Figure 5D, supplementary figure 4C, 6C). Consistently, multiple genes involved in the MTORC1 pathway, such as *Rpn1, Xbp1, Eif2s2*, were upregulated in the BTC group (Figure 5E). Moreover, as a readout of MTORC1 kinase activity, p-S6 level was significantly enhanced in tumors from BTC mice (Figure 5F). To further investigate the role of TERT in MTORC1 signaling in thyroid cancer, we expressed WT-TERT or DN-TERT (mutant without telomerase activity) in MDA-T41 cells and analyzed the phosphorylation of known downstream effectors of MTORC1 (Figure 5G, 5H). In agreement with *in vivo* results, expression of WT-TERT effectively induced phosphorylation of S6 and S6K, especially at serum-starved conditions (Figure 5H,5I). Hence, MTORC1 might participate in the activation of ribosomes by TERT.

### Targeted inhibition of rRNA transcription blocked tumor progression

Targeting downstream TERT mechanisms, such as ribosome biogenesis and MTORC1 signaling, may serve as alternative anti-tumor strategies with reduced side effects compared with traditional TERT-targeted drugs. We analyzed Genomics of Drug Sensitivity in Cancer (GDSC) datasets(43) and observed that cells with higher TERT expression levels were more sensitive to CX-5461, an inhibitor of POL I-mediated rRNA transcription, which was tolerated in normal somatic cells. CX-5461 presented a pattern like previous telomerase targeted drugs, such as BIBR 1532 and telomerase inhibitor IX (Figure 6A) (44–48). *BRAF* V600E and *TERT* promoter co-mutant BCPAP (thyroid cancer cell line) and OCM1 (melanoma cell line) cells, and a primary cell line 312BTC derived from BTC thyroid tumor, were used to test the effect of CX-5461. CX-5461 inhibited cancer cell proliferation in a dose-dependent manner (Figure 6B-6D), comparable to the effects of TERT knockdown (Supplementary Figure 2G). Thyroid BCPAP and melanoma OCM1 subcutaneous xenograft models were established and CX-5461 was orally administrated to test the anti-tumor effect of CX-5461 *in vivo*. We found that CX-5461 repressed tumor growth effectively with no significant weight loss (Figure 6E-6J). Moreover, Ki67 expression in tumors was also reduced upon CX-5461 treatment (Figure 6K-6N). Together, these results indicated that CX-5461 was effective in inhibiting the proliferation of cells with *BRAF* mutation and TERT reactivation.

**Figure 6.**
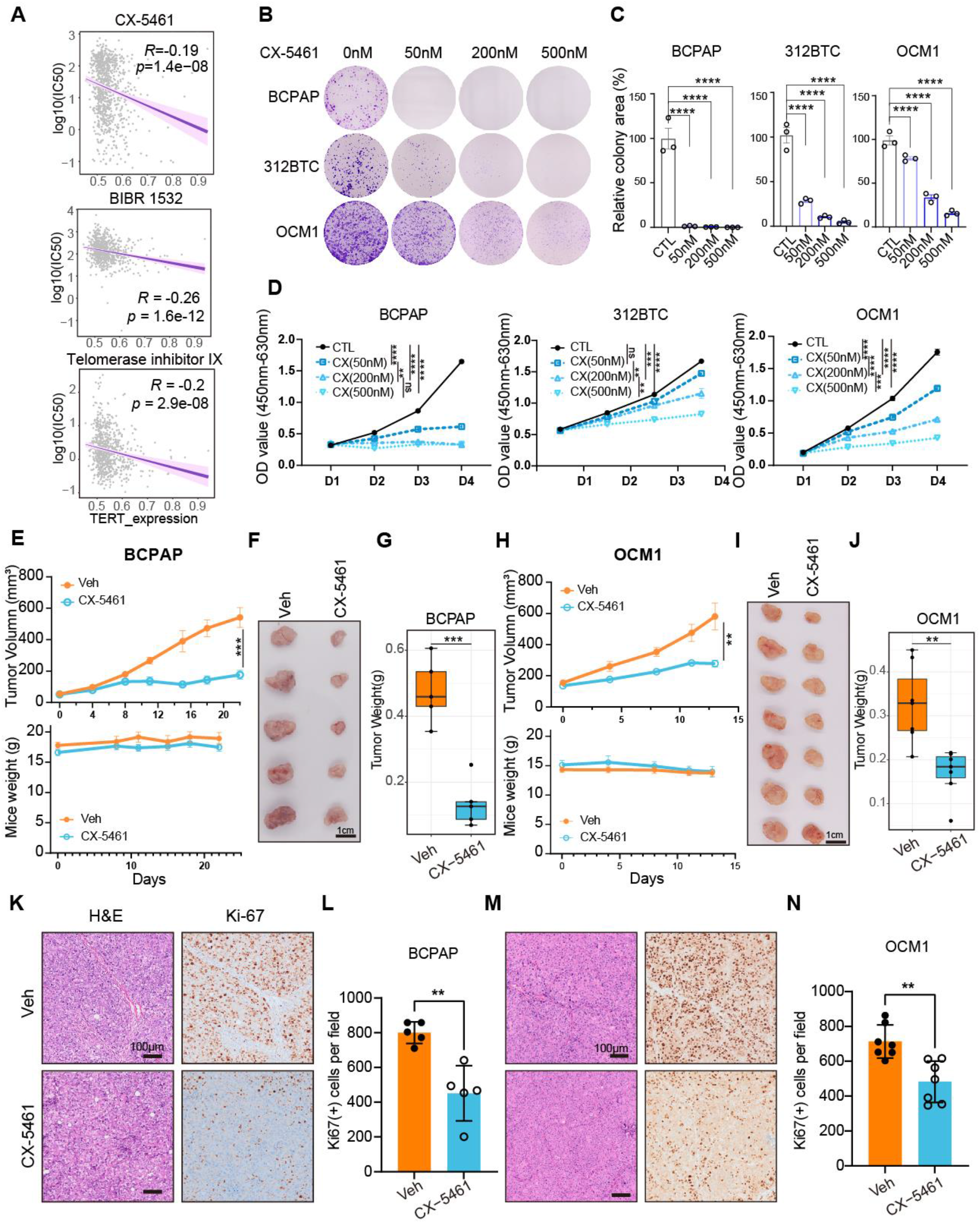
CX-5461 inhibited multiple tumor progression. **A**. Correlation between TERT expression and log10 (IC50) of CX-5461, BIBR 1532, telomerase inhibitor IX in GDSC datasets. Spearman correlation test. **B**. Effects of concentration gradient CX-5461 on colony formation of BCPAP, 312BTC, and OCM1 cells. **C**. Quantifications results of B. **D**. CCK8 assays concentration gradient CX-5461 effect on BCPAP, 312BTC, and OCM1 cells. Data represent the mean ± SEM (n=3). ns, not significant, **, *p* < 0.01, ***, *p* < 0.001, ****, *p* < 0.0001. One-way ANOVA and Dunnett multiple comparisons test. **E. F. G. H**. Growth curve, mice weight, tumor gross appearance and tumor weight of B-CPAP xenografts treated by vehicle or CX-5461. n=5 for each group. P. O. for 5 times. **I. J. K. L**. Growth curve, mice weight, tumor gross appearance, and tumor weight of OCM1 xenografts treated by vehicle or CX-5461. n=7 for each group. P. O. for 4 times. **M, O** H&E, Ki-67 staining of BCPAP and OCM1 xenografts with vehicle or CX-5461. scale bar=100 μm. **N, P** Quantifications results of Ki-67 of **M, O** per field (10X objective). Data assessed by Student’s t-test. **, *p*<0.01. **E, H** Data represent the mean ± SEM. **G, J** Data represent the median with IQR and 1.5 IQR whiskers. **L, N** Data represent the mean ± SD.

### CX-5461 induced thyroid cancer re-differentiation

Currently, radioactive iodine (RAI) therapy is regarded as the first-line treatment for metastatic thyroid cancers (49). However, nearly all PDTC and ATC, as well as some of the well-differentiated thyroid cancer, do not uptake iodine because of the deficiency of functional thyroid follicular markers especially sodium iodide symporter (NIS). These patients are classified as RAI refractory, and RAI refraction is one of the leading causes of thyroid cancer-related deaths (50). Given that the major role of TERT in tumor progression is to induce tumor dedifferentiation, blocking TERT or its downstream effectors may lead to cancer differentiation.

Thus, the effect of CX-5461 on differentiation markers in thyroid cancer cells was evaluated. Strikingly, the expression levels of *PAX8, DIO1, THRA, THRB, FOXE1*, and *TSHR* in K1, BAPAP, and SW1736 cells were significantly upregulated after 24 hours or 48 hours of 8 CX-5461 treatment (Figure 7A, 7B, 7C). NIS and PAX8 protein expression levels were also elevated in K1 and SW1736 cells after treatment with CX-5461 (Figure 7D, 7E, 7F). Consistently, high NIS expression was observed in the BCPAP tumors with CX-5461 treatment (Figure 7G), suggesting that rRNA transcription blockade was an effective way to induce thyroid cancer re-differentiation.

**Figure 7.**
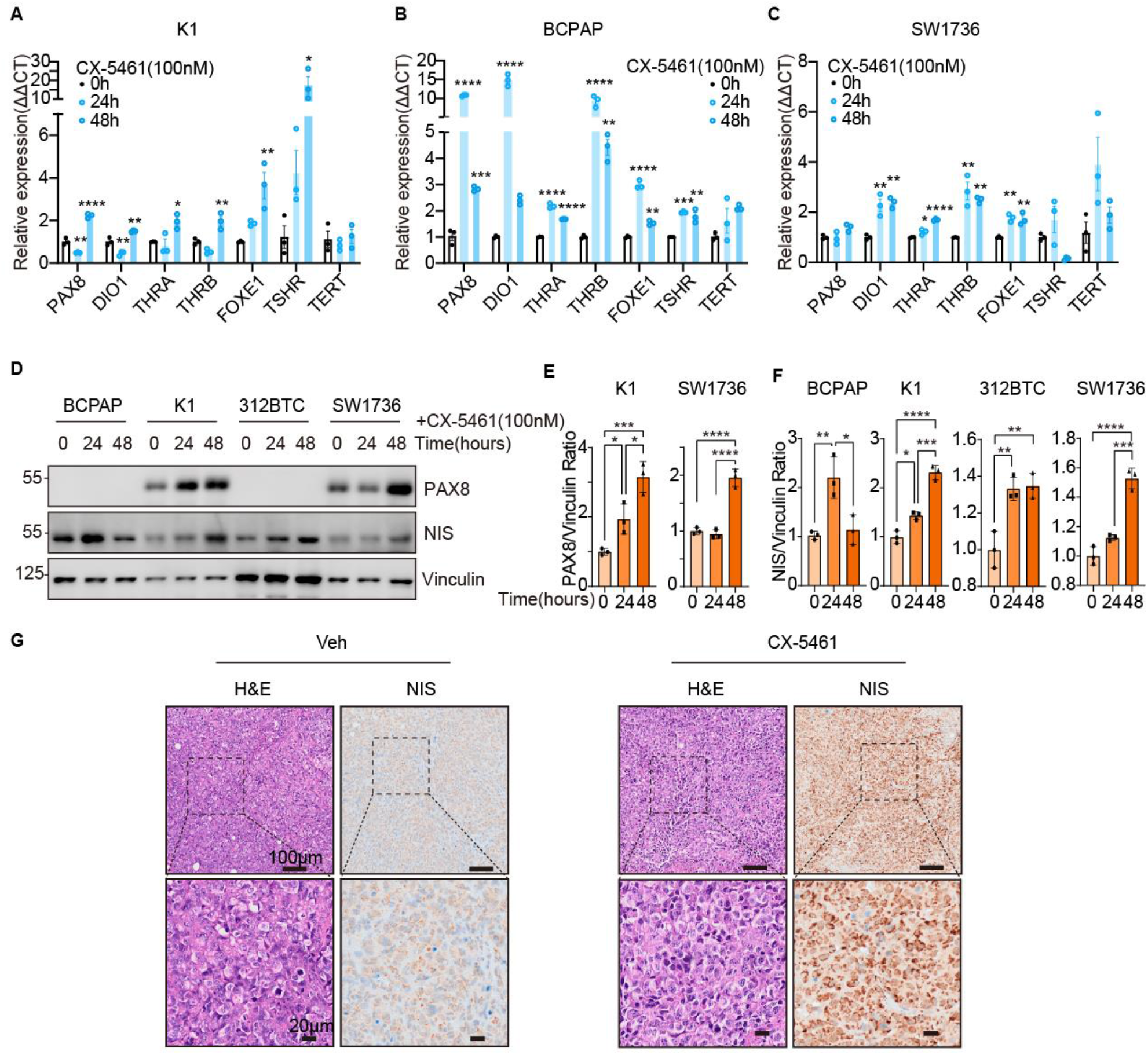
CX-5461 induced thyroid cancer re-differentiation. **A, B, C** Representative thyroid differentiation-associated genes PAX8, DIO1, THRA, THRB, FOXE1, and TSHR qPCR results of K1, BCPAP, and SW1736 after CX-5461 (100nM) treatment for 0h, 24h, and 48h. Data represent the mean ± SEM (n=3). **D**. Western blot showed PAX8 and NIS change of K1, BCPAP, 312BTC, and SW1736 treated by CX-5461 (100nM) for 0h, 24h, and 48h. **E. and F**. Quantification results of PAX8 and NIS change in D. Data represent the mean ± SD (n=3). **G**. H&E morphology and anti-NIS staining after 5 times CX-5461 treatment in BCPAP xenografts. *, *p*<0.05; **, *p*<0.01; ***, *p*<0.001; ****, *p*<0.0001 versus 0h by one-way ANOVA and Dunnett multiple comparisons test.

**Figure 8.**
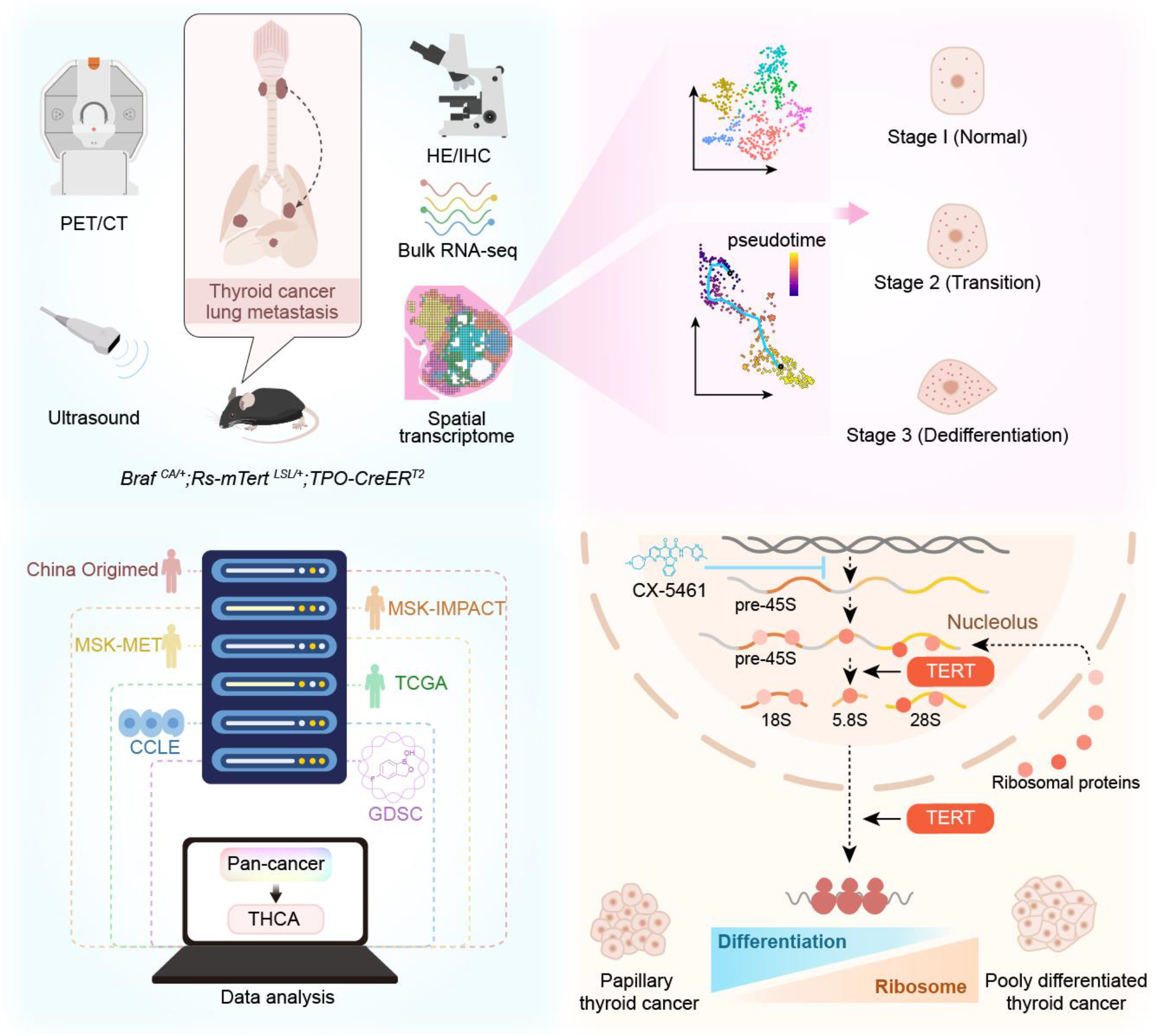
Schematic Diagram of the present study. Comprehensive integration of animal models, spatial transcriptome sequencing, databases analysis and experiments revealed the role of TERT in ribosomal pathways.

## Discussion

We demonstrated in this study that conditional transgenic expression of TERT promoted dedifferentiation and cancer metastasis driven by *BRAF* V600E mutation in mouse thyroid. The expression of TERT in both mouse and human *BRAF* V600E tumor cells induced rRNA transcription and ribosomal biogenesis. The rRNA transcription inhibitor CX-5461 induced tumor differentiation and retarded progression, reversing TERT-induced oncogenic features effectively.

A large proportion of human PTC maintain stability or progress rather slowly during active surveillance, while poorly-differentiated and anaplastic thyroid cancers usually exhibit a sudden increase in tumor size and abrupt deterioration (51). Previous studies have modeled thyroid cancer in mice, mainly by transgenic expression or conditional mutation of BRAF. The *Tg-Braf* mice exhibited a PDTC pattern at 3 months old. Transgenic expression of *Braf* ^*CA*^ under *Tg* promoter at the embryoid stage resulted in elevated TSH levels and destroyed thyroid function after birth. A high dose of *Braf* ^*CA*^ may contribute to early tumor formation and dedifferentiation (52,53). In the *Braf* ^*CA*^*;TPO-CreER*^*T2*^ (BC) model, *BRAF*^*CA*^ expression was induced by one dose of tamoxifen at 2 months of age to mimic *BRAF* mutation in human thyroid cancer. No lung metastasis or profound tumor dedifferentiation was observed (54). In our study, tumor progression in BC and BTC mouse models was monitored by ultrasound imaging. Tumors in BTC mice exhibited a sudden increase in tumor size and metastasis (Figure 1, 2). Spatial transcriptomic analysis of a tumor from a BTC mouse revealed a continuous trajectory of dedifferentiation of thyroid cancer cells (Figure 3). The *BRAF*-activating mutation presents in both PTC and PDTC, while TERT reactivation occurs more frequently in PDTC(55). Both molecular and phenotypic transitions from PTC to PDTC in human thyroid cancers were mimicked in our genetic mouse models. The biological behavior of tumors from BC and BTC mice was of high identity to that of PTC and PDTC, respectively. Studies of BC and BTC mouse models indicated an essential role of TERT in the progression of thyroid cancer into the advanced stage.

TERT reactivation is observed frequently in various cancer types. Studies have found a crosstalk between TERT and important signaling pathways, such as NF-κB, Wnt pathway, FOXO3a, and MYCs (12–14). In addition, TERT stimulated Pol I activity and tRNA transcription (15,16). Our study highlighted important roles of rRNA transcription and ribosomal biogenesis in cancer progression upon TERT reactivation (Figure 4, Figure 5). With regards to how TERT regulates rRNA transcription and ribosomal activity, our analysis indicated that TERT activated MTORC1 signaling, which induces ribosomal biogenesis at both rRNA transcription and translation efficiency levels (Figure 5)(42). Several ribosome scaffold proteins appeared to interact with TERT as potential mediators in regulating ribosome activity (Figure 5B, 5C). Further investigations are required to understand the precise mechanism linking TERT and ribosomal activity.

Ribosomal biogenesis is crucial for various cancer cell activities including survival, proliferation, differentiation, and metastasis. Ribosomes are therefore attractive molecular targets for cancer drugs (34,56,57). CX-5461 is a small molecule that targets rRNA transcription, which has been reported to repress the proliferation of melanoma, breast cancer, and ovarian cancer (44–47). There is an ongoing clinical trial (*NCT04890613*) exploring the role of CX-5461 in *BRCA1/2, PALB2*, or homologous recombination deficiency (HRD) mutant solid tumors. We have shown that CX-5461 effectively inhibited the proliferation of different cancer cells with *BRAF* V600E and *TERT* promoter mutations (Figure 6). Moreover, CX-5461 successfully increased the expression of thyroid differentiation markers including PAX8, THRA, THRB, and NIS of human and murine thyroid tumors, suggesting re-differentiation. These results provide strong evidence supporting the effectiveness of CX-5461 in treating advanced thyroid tumors, especially those dedifferentiated thyroid cancers insensitive to radioiodine therapies.

## Methods

### Mouse models and administration strategy

In the C57BL6/J background, C57BL/6-*Gt(ROSA)26Sor*^*em1(CAG-LSL-Tert-3xFlag-IRES-EGFP-WPRE-polyA)Smoc*^ conditional TERT overexpression model was generated by *loxp-stop-loxp-Tert-3xflag-IRES-EGFP-polyA* sequence insertion in mouse *Rosa26* foci. Tg (TPO-cre/ ERT2)1139Tyj (Strain #:026512) and B6;129-Braf ^tm1Mmcm/+^ (Strain #:017837) mice were obtained from Jackson lab. Genotyping of Braf and TPO-CreER^T2^ mice was performed separately according to Jackson protocol. Primers for Rs26-mTert mice genotyping were described in the supplementary table S1. Expression of the transgene was induced using 20mg/ml Tamoxifen (MCE, HY-13757A) by intraperitoneal administration 4 times at 1 month of age. All animals were housed in the SPF environment. Animal care and experimental procedures were performed with approval by Fudan University Animal Ethics Community.

### Xenograft model

Human thyroid cancer cell line BCPAP and human uveal melanoma cell line OCM1 were used to construct a subcutaneous tumor model. 5*10^6^ cells (100 μL volume per injection) per mouse were subcutaneously injected into BALB/c nude mice (4-5 weeks old). Mice with tumors were randomized in the vehicle and CX-5461 treatment groups when the average tumor size reached about 60-150 mm^3^. 50mg/kg CX-5461 (MCE, HY-13323A) was administrated by oral gavage twice a week. Mice were sacrificed when the average tumor size of the vehicle group reached 1000mm^3^.

### Cell culture

HEK 293T, OCM1, and BCPAP cells were maintained in DMEM (BasalMedia, L110KJ), and NTHY-ori 3-1, MDA-T41, SW1736, and RKO were maintained in RPMI-1640 (BasalMedia, L210KJ). Media were supplemented with 10% fetal bovine serum (ExCell Bio, FSP500) and 50mg/ml Penicillin/ Streptomycin (Meilunbio, MA0110) unless otherwise stated in the text. All the cells were incubated at 37°C, with 5% CO_2_.

### Establishment of primary mouse thyroid cell line

Primary thyroid cells were separated and cultured as previously described(58). After dissection using scissors, thyroid tissues were digested in F-12 medium (Gibco, 11765-054) with the following regents:100U/ml Collagenase A (Gibco, 10103578001), 1mg/ml Dispase II (Solarbio, D6430), 10% FBS, 50mg/ml P/S at 37°C for 1 hour and filtered with 70 μm cell strainer.

The first three passages were cultured in F-12 medium with 5ug/ml transferrin, 10ug/ml Bovine insulin (Biosharp, BS001), 3.5ng/ml hydrocortisone (Solarbio, G8450), 2ng/ml Glycyl-L-histidyl-L-lysine (MCE, HY-P0046), and 10ng/ml somatostatin (Meilunbio, MB1225). After three passages, cells were cultured in a DMEM medium with 10% FBS.

### Plasmids and lentivirus package

For overexpression, the target mRNA CDS sequence was cloned using PrimeSTAR® Max (TAKARA, R045) or Q5® High-Fidelity DNA Polymerase (NEB, M0491). After separation by agarose gel electrophoresis and retraction, the fragments were subcloned into the pLVX-puro vector (Takara, #632164) with Ha- or Flag-tag. ShRNAs targeting TERT were cloned into pLKO.1. Transfection was performed using PolyJet™ DNA In Vitro Transfection Reagent (Signagen, #SL100688) or PEI (Polysciences Inc, 23966-2) following manufacturer’s instructions. For lentivirus production, pLVX-based plasmids were co-transfected into HEK293T cells together with psPAX2 and pMD.2g.

### Transfection of plasmids and siRNA

For siRNA transfection, control siRNA, TERT siRNA, and TERC siRNA were separately transfected into cultured cells using riboFECT CP Transfection Kit (RIBOBIO, C10511-05). The sequences of siRNAs were listed in the supplementary table S1.

### Animal ultrasound imaging and PET-CT

Fujifilm Vevo 2100 was adopted to monthly monitor the thyroid morphology of mice. Briefly, after anesthesia by isoflurane and dehairing, an MS400 detector was used to scan the neck region. All images were collected using the same preset.

Micro-PET/CT (Inveon, Siemens) scanning was performed at the Department of Nuclear Medicine, Fudan University Shanghai Cancer Center. After fasting overnight, ^18^F-FDG was intraperitoneally injected for 45 min before scanning. The PET and CT images were fused using Inveon Research Workplace software (Siemens Medical Solutions). Data were expressed in the form of standardized uptake value (SUV).

### IHC staining

Formalin-fixed, paraffin-embedded sections were baked at 65°C for 1h. After deparaffinization and hydration, sodium citrate buffer (Beyotime, P0083) was adopted for heat-treated antigen and boiled for 15 minutes at 100°C. Subsequently, endogenous peroxidase activities were blocked using 3% H_2_O_2_ for 20 min. After blocking with 3% BSA/TBST for 1h at room temperature, the primary antibodies were incubated at the optimized concentration at 4°C overnight. DAKO (K4003) and GenTech (GK500705) secondary HRP-conjugated antibodies were separately incubated for 1h and reacted with DAB. Hematoxylin was counterstained for nuclear staining. Sections were scanned using Olympus VS200.

### RNA extraction, cDNA synthesis, and RT-qPCR

Total RNA of adherent cells was isolated with the EZB Kit (EZB, B0004DP), and the total RNA of mouse thyroid tissues was isolated with the Tiangen Kit (Tiangen, DP419), then reverse-transcribed to cDNA using the One-Step gDNA Removal and cDNA Synthesis SuperMix (TransGen, AU311). The amplification and fluorescence detection of quantitative RT-qPCR was performed in Q6 (Invitrogen), using the TAKARA TB Green Premix (RR420). The relative expression level of target genes was calculated with the ΔΔCT method, with β-actin as the internal control. The primer sequences were listed in the supplementary table S1.

### Bulk RNA Sequencing

The total RNA samples (1μg) were used for the following library preparation. Double-stranded cDNA was purified and treated to repair both ends and add adaptors to them. Each sample was then amplified by PCR using P5 and P7 primers and the PCR products were validated. Libraries with different indexes were multiplexed and loaded on Illumina Novaseq 6000 instrument for sequencing. Clean data of BTC and BC mice thyroid were aligned to reference genome via software Hisat2 (v2.0.1). GRCm39.104 was used as a reference genome in alignment. Gene expression levels were estimated using HTseq (v0.6.1) and differential expression analysis used the DEseq2(59) Bioconductor package. PCA analysis and visualization were performed with R packages FactoMineR(v2.4) and factoextra (v1.0.7). Single-sample GSVA (ssGSVA) and GSEA were performed to determine the Hallmarks of Cancer phenotypes of BTC and BC thyroid cancers. Visualizations were generated using ggplot2 (v3.3.6) and pheatmap (v1.0.12) R packages.

### Spatial transcriptome sample preparation, library construction and sequencing

Fresh thyroid tissues of BTC mice were fast-frozen in the pre-cooled isopentane and embedded with the Optimal Cutting Compound (OCT) (Sakura, 4583). Tissues were stored at -80°C. The optimal permeabilization time was found to be 12 min using 10x Genomics Visium Tissue Optimization Kit. Spatially tagged cDNA libraries were constructed with the 10x Genomics Visium Spatial Gene Expression 3’ Library Construction V1 Kit and then sequenced on Novaseq (Illumina)

### Spatial transcriptome data processing and visualization

Spaceranger-1.1.0 pipeline was used to process raw fastq files with mm10 reference genome, and the output files were integrated using Seurat (v4.1.0) R package. Seurat SCTransform function was used for normalization. FindNeighbors and FindClusters functions of Seurat were used for downstream graph-based clustering, and t-sne dimensionality reduction was then used for projection visualization(60). To identify the marker genes of the clusters, FindAllMarkers function was performed (logFC.threshold=0.25, test.use=“wilcox”). Seurat’s AddModuleScore function was applied to quantify the differentiation scores of 312BTC thyroid spots.

### Trajectory analysis

Trajectory inference and pseudotime analysis were performed using Monocle3 (v1.2.9). All spots of the 312BTC sample were integrated using new_cell_data_set function and thyrocytes were chosen using the choose_cells function. Cluster c0 was indicated as the root.

### Single-cell transcriptome enrichment analysis

We adopted AUCell (v1.16.0) to calculate the enrichment score of each spot with Reactome database. Visualization was performed with ComplexHeatmap (v2.10.0).

### EU assay

To label nascent RNA, Cell-Light EU RNA Imaging Kit (RIBOBIO, C10316) was adopted. For OCM1 and K1 cells, EU treatment lasted for 3 hours, then the EU-labeled RNA was detected following the protocol. To visualize the nucleolar region, the anti-FBL immunofluorescence was co-stained.

### Immunoblotting

Whole-cell lysates were separated by sodium dodecyl sulfate-polyacrylamide gel electrophoresis (SDS-PAGE), and proteins were transferred onto nitrocellulose membranes. Following blocking in 5% non-fat milk, membranes were incubated with primary antibodies in 3% bovine serum albumin (BSA) overnight at 4°C, and then with secondary antibodies in 5% milk for 1h at room temperature. High-sig ECL Western Blotting Substrate (Tanon, #180-501) was adopted for chemiluminescence detection with Tanon 5200S imaging system or BioRad XRS+ imaging system.

### Immunoprecipitation

Cells were lysed with mild lysis buffer (50 mM HEPES at pH 7.5, 150 mM NaCl, 1 mM EDTA, 1% NP-40, 10 mM pyrophosphate, 10 mM glycerophosphate, 50 mM NaF, and 1.5 mM Na3VO4) containing EDTA-Free Protease Inhibitor Cocktail (Bimake, B14012). After sonication and centrifugation, the supernatants were incubated with primary antibodies for 1h at 4°C. Protein A beads were then added and incubated for additional 2 hours, followed by four times centrifugation at 2000 rpm and washing with mild lysis buffer at 4°C. Proteins precipitated were dissolved in 1x SDS-PAGE sample buffer and followed by immunoblotting.

### IP-LC/MS

For the identification of the FLAG-TERT interaction proteomics, about 10^7^ HEK 293T cells were transfected with 5ug pLVXyu-FLAG-TERT plasmid. The immunoprecipitated proteins were separated by SDS-PAGE using the 4-15% Tris/gly gel. After in-gel digestion, the peptides were analyzed by online nanoflow liquid chromatography-tandem mass spectrometry performed on an EASY-nanoLC 1200 system (Thermo Fisher Scientific, MA, USA) connected to a Q Exactive™ Plusmass spectrometer (Thermo Fisher Scientific, MA, USA). Tandem mass spectra were processed by PEAKS Studio version X+ (Bioinformatics Solutions Inc., Waterloo, Canada). Interacting proteins identified are shown in the supplementary table S3.

### Telomere Repeat Amplification Protocol (TRAP) assay

The telomerase activity was measured using the TRAPeze® Telomerase Detection Kit (Millipore, S7700). Briefly, 10^6^ cells were lysed in 200ul 1xCHAPS buffer with RNase inhibitor. After centrifugation and BCA protein quantification, the telomerase activity to length telomere template was detected using the PCR method with the procedure: 30°C, 30min, 1 cycle; 95°C, 2min, 1 cycle; 94°C, 15sec, 59°C, 30sec, 72°C, 1min, 32 cycles; 4°C keep. The PCR products were separated using the 10% Acis/Bis Gel, followed by staining with GelRed (Tsingke, TSJ003), using 50bp DNA ladder (Generay, DL0501) as the reference.

### SUnSET

The SUnSET assay was performed just as previously reported(61). Briefly, cells without puromycin resistance were labeled with 10ug/ml Puromycin (Invivogen, ant-pr) for 10min, followed by a 1h chase. Then, the anti-Puromycin primary antibody (Sigma-Aldrich, MABE343) was used to detect the puromycin incorporation with the immunoblot assay.

### Bicistronic Dual Luciferase Reporter

The plasmids used in the bicistronic reporter assay were gifts from Mark T. Bedford Lab. After 24h transfection of reporter plasmids, 10^5^ HEK293T cells were seeded in a 24-well plate. Then, 48h after si-RNA transfection, cell lysates were treated with the dual luciferase measurement kit (Transgene, FR201), and the firefly and renilla luciferase activities were measured using the BioTek Synergy LX, with β-ACTIN protein level (by western blot) as the internal control.

### Public datasets analysis

MSK-IMPACT, MSK-MET, China Origimed2020 cohorts, TCGA, and CCLE datasets were accessed through cbioportal (https://www.cbioportal.org/datasets). CCLE counts data were downloaded from Xena. Visualization of TERT alteration frequency and cancer mutation landscape was performed with ggplot2 (v3.3.6) R package.

### Colony formation

1000 to 3000 cells were seeded in 12-well plates. After culture for 1-2 weeks, the cells were fixed by 4% PFA for 10 min, followed by PBS washing and 0.1% crystal violet (Servicebio, G1014) staining for 15 min. Colony areas were measured using Image J.

### CCK8

About 5,000 to 10,000 cells were seeded in each well of 96-well plates. For CCK8 assay, the cells were added with CCK8 solution (meilunbio, MA0218) and incubated for 1h at 37°C. The absorbance was measured at 450nm and at a reference wavelength of 630nm with BioTek Synergy LX.

### Statistics

Comparisons between two groups were performed using Student’s two-sided t-test. One-way ANOVA and Dunnett’s multiple comparisons test were adopted for multiple comparisons. All statistical analyses were performed using GraphPad Prism v9.0.0 or R package ggsignif (v0.6.3). P value less than 0.05 was considered as significant. The number of replicates was noted in the figures and legends.

## Author contributions

Conceptualization, Y.-L.W., F.Y. and N.Q.; Experiments: P.Y., R.Z. J.H., L.T., C.F., Y.L., J.L. and C.-X.H; Bioinformatics, P.Y., P.H., C.F., and X.S.; Pathological analysis, H.G., Y.-L.W., W.W. and X.S.; Validation, J.H., P.Y., C.H., L.T., and J.H.; Visualization, P.Y., P.H.; Writing-Original Draft, P.Y., F.Y., J.H., N.Q. and C.H.; Writing-Review & Editing, Y.-L.W., F.Y., Y.W., and W.W.; Supervision, Y.-L.W., F.Y., F.L. and Q.J.

## Notes

**Funding Statement:** This work was supported by grants from the Ministry of Science and Technology of China (National Key R&D program, 2018YFA0800304 and 2020YFA0803202), the Science and Technology Commission of Shanghai Municipality (21S11905000), and the Shanghai Municipal Health Commission (2022XD049) to Fa-Xing Yu, and grants from the National Natural Science Foundation of China (81972501) and Shanghai Shenkang Hospital Development Center (SHDC22021201) to Yu-Long Wang.

**Disclosure:** The authors have no conflicts of interest to declare.

### Competing Interest Statement

The authors have declared no competing interest.

## References

1. Kim NW, Piatyszek MA, Prowse KR, Harley CB, West MD, Ho PL, et al. Specific association of human telomerase activity with immortal cells and cancer. Science. 1994;266:2011–5.

2. He Y, Wang Y, Liu B, Helmling C, Sušac L, Cheng R, et al. Structures of telomerase at several steps of telomere repeat synthesis. Nature. 2021;593:454–9.

3. Hohaus S, Voso MT, Orta-La Barbera E, Cavallo S, Bellacosa A, Rutella S, et al. Telomerase activity in human hematopoietic progenitor cells. Haematologica. 1997;82:262–8.

4. Li X, Qian X, Wang B, Xia Y, Zheng Y, Du L, et al. Programmable base editing of mutated TERT promoter inhibits brain tumour growth. Nat Cell Biol. Nature Publishing Group; 2020;22:282–8.

5. Bell RJA, Rube HT, Kreig A, Mancini A, Fouse SD, Nagarajan RP, et al. Cancer. The transcription factor GABP selectively binds and activates the mutant TERT promoter in cancer. Science. 2015;348:1036–9.

6. Pirker C, Bilecz A, Grusch M, Mohr T, Heidenreich B, Laszlo V, et al. Telomerase Reverse Transcriptase Promoter Mutations Identify a Genomically Defined and Highly Aggressive Human Pleural Mesothelioma Subgroup. Clin Cancer Res. 2020;26:3819–30.

7. Amen AM, Fellmann C, Soczek KM, Ren SM, Lew RJ, Knott GJ, et al. Cancer-specific loss of TERT activation sensitizes glioblastoma to DNA damage. Proc Natl Acad Sci U S A. 2021;118:e2008772118.

8. Guterres AN, Villanueva J. Targeting telomerase for cancer therapy. Oncogene. 2020;39:5811–24.

9. Ruden M, Puri N. Novel anticancer therapeutics targeting telomerase. Cancer Treat Rev. Elsevier; 2013;39:444–56.

10. Ait-Aissa K, Ebben JD, Kadlec AO, Beyer AM. Friend or Foe? Telomerase as a Pharmacological target in Cancer and Cardiovascular Disease. Pharmacol Res. 2016;111:422–33.

11. Rha SY, Izbicka E, Lawrence R, Davidson K, Sun D, Moyer MP, et al. Effect of Telomere and Telomerase Interactive Agents on Human Tumor and Normal Cell Lines1. Clin Cancer Res. 2000;6:987–93.

12. Koh CM, Khattar E, Leow SC, Liu CY, Muller J, Ang WX, et al. Telomerase regulates MYC-driven oncogenesis independent of its reverse transcriptase activity. J Clin Invest. 2015;125:2109–22.

13. Ghosh A, Saginc G, Leow SC, Khattar E, Shin EM, Yan TD, et al. Telomerase directly regulates NF-κB-dependent transcription. Nat Cell Biol. 2012;14:1270–81.

14. Hu C, Ni Z, Li B, Yong X, Yang X, Zhang J, et al. hTERT promotes the invasion of gastric cancer cells by enhancing FOXO3a ubiquitination and subsequent ITGB1 upregulation. Gut. BMJ Publishing Group; 2017;66:31–42.

15. Khattar E, Kumar P, Liu CY, Akıncılar SC, Raju A, Lakshmanan M, et al. Telomerase reverse transcriptase promotes cancer cell proliferation by augmenting tRNA expression. J Clin Invest. 2016;126:4045–60.

16. Gonzalez OG, Assfalg R, Koch S, Schelling A, Meena JK, Kraus J, et al. Telomerase stimulates ribosomal DNA transcription under hyperproliferative conditions. Nat Commun. 2014;5:4599.

17. Henson JD, Neumann AA, Yeager TR, Reddel RR. Alternative lengthening of telomeres in mammalian cells. Oncogene. Nature Publishing Group; 2002;21:598–610.

18. Ding Z, Wu C-J, Jaskelioff M, Ivanova E, Kost-Alimova M, Protopopov A, et al. Telomerase Reactivation following Telomere Dysfunction Yields Murine Prostate Tumors with Bone Metastases. Cell. Elsevier; 2012;148:896–907.

19. Strong MA, Vidal-Cardenas SL, Karim B, Yu H, Guo N, Greider CW. Phenotypes in mTERT+/− and mTERT−/− Mice Are Due to Short Telomeres, Not Telomere-Independent Functions of Telomerase Reverse Transcriptase ▿. Mol Cell Biol. 2011;31:2369–79.

20. González-Suárez E, Samper E, Ramírez A, Flores JM, Martín-Caballero J, Jorcano JL, et al. Increased epidermal tumors and increased skin wound healing in transgenic mice overexpressing the catalytic subunit of telomerase, mTERT, in basal keratinocytes. EMBO J. 2001;20:2619–30.

21. Landa I, Ibrahimpasic T, Boucai L, Sinha R, Knauf JA, Shah RH, et al. Genomic and transcriptomic hallmarks of poorly differentiated and anaplastic thyroid cancers. J Clin Invest. 2016;126:1052–66.

22. Jonsson P, Lin AL, Young RJ, DiStefano NM, Hyman DM, Li BT, et al. Genomic Correlates of Disease Progression and Treatment Response in Prospectively Characterized Gliomas. Clin Cancer Res Off J Am Assoc Cancer Res. 2019;25:5537–47.

23. Zehir A, Benayed R, Shah RH, Syed A, Middha S, Kim HR, et al. Mutational landscape of metastatic cancer revealed from prospective clinical sequencing of 10,000 patients. Nat Med. 2017;23:703–13.

24. Ghandi M, Huang FW, Jané-Valbuena J, Kryukov GV, Lo CC, McDonald ER, et al. Next-generation characterization of the Cancer Cell Line Encyclopedia. Nature. 2019;569:503–8.

25. Yu P-C, Tan L-C, Zhu X-L, Shi X, Chernikov R, Semenov A, et al. Arms-qPCR Improves Detection Sensitivity of Earlier Diagnosis of Papillary Thyroid Cancers With Worse Prognosis Determined by Coexisting BRAF V600E and Tert Promoter Mutations. Endocr Pract Off J Am Coll Endocrinol Am Assoc Clin Endocrinol. 2021;27:698–705.

26. Cancer Genome Atlas Research Network. Integrated genomic characterization of papillary thyroid carcinoma. Cell. 2014;159:676–90.

27. Cheung CC, Ezzat S, Freeman JL, Rosen IB, Asa SL. Immunohistochemical Diagnosis of Papillary Thyroid Carcinoma. Mod Pathol. Nature Publishing Group; 2001;14:338–42.

28. Chernock RD. Immunohistochemistry of thyroid gland carcinomas: clinical utility and diagnostic pitfalls. Diagn Histopathol. 2016;22:184–90.

29. Besic N, Gazic B. Sites of Metastases of Anaplastic Thyroid Carcinoma: Autopsy Findings in 45 Cases from a Single Institution. Thyroid. Mary Ann Liebert, Inc., publishers; 2013;23:709–13.

30. Liberzon A, Birger C, Thorvaldsdóttir H, Ghandi M, Mesirov JP, Tamayo P. The Molecular Signatures Database (MSigDB) hallmark gene set collection. Cell Syst. 2015;1:417–25.

31. Pu W, Shi X, Yu P, Zhang M, Liu Z, Tan L, et al. Single-cell transcriptomic analysis of the tumor ecosystems underlying initiation and progression of papillary thyroid carcinoma. Nat Commun. 2021;12:6058.

32. Stefanovsky VY, Pelletier G, Hannan R, Gagnon-Kugler T, Rothblum LI, Moss T. An Immediate Response of Ribosomal Transcription to Growth Factor Stimulation in Mammals Is Mediated by ERK Phosphorylation of UBF. Mol Cell. Elsevier; 2001;8:1063–73.

33. Ebright RY, Lee S, Wittner BS, Niederhoffer KL, Nicholson BT, Bardia A, et al. Deregulation of ribosomal protein expression and translation promotes breast cancer metastasis. Science. 2020;367:1468–73.

34. Pelletier J, Thomas G, Volarević S. Ribosome biogenesis in cancer: new players and therapeutic avenues. Nat Rev Cancer. 2018;18:51–63.

35. Martin R, Hackert P, Ruprecht M, Simm S, Brüning L, Mirus O, et al. A pre-ribosomal RNA interaction network involving snoRNAs and the Rok1 helicase. RNA. 2014;20:1173–82.

36. Guerrieri AN, Zacchini F, Onofrillo C, Di Viggiano S, Penzo M, Ansuini A, et al. DKC1 Overexpression Induces a More Aggressive Cellular Behavior and Increases Intrinsic Ribosomal Activity in Immortalized Mammary Gland Cells. Cancers [Internet]. 2020 [cited 2021 May 26];12. Available from: https://www.ncbi.nlm.nih.gov/pmc/articles/PMC7760958/

37. Cheng L, Yuan B, Ying S, Niu C, Mai H, Guan X, et al. PES1 is a critical component of telomerase assembly and regulates cellular senescence. Sci Adv [Internet]. 2019 [cited 2021 May 2];5. Available from: https://www.ncbi.nlm.nih.gov/pmc/articles/PMC6520020/

38. Lapik YR, Fernandes CJ, Lau LF, Pestov DG. Physical and Functional Interaction between Pes1 and Bop1 in Mammalian Ribosome Biogenesis. Mol Cell. 2004;15:17–29.

39. Ruggero D, Grisendi S, Piazza F, Rego E, Mari F, Rao PH, et al. Dyskeratosis congenita and cancer in mice deficient in ribosomal RNA modification. Science. 2003;299:259–62.

40. Chen W, Bucaria J, Band DA, Sutton A, Sternglanz R. Enp1, a yeast protein associated with U3 and U14 snoRNAs, is required for pre-rRNA processing and 40S subunit synthesis. Nucleic Acids Res. 2003;31:690–9.

41. Nguyen LXT, Mitchell BS. Akt activation enhances ribosomal RNA synthesis through casein kinase II and TIF-IA. Proc Natl Acad Sci U S A. 2013;110:20681–6.

42. Iadevaia V, Liu R, Proud CG. mTORC1 signaling controls multiple steps in ribosome biogenesis. Semin Cell Dev Biol. 2014;36:113–20.

43. Yang W, Soares J, Greninger P, Edelman EJ, Lightfoot H, Forbes S, et al. Genomics of Drug Sensitivity in Cancer (GDSC): a resource for therapeutic biomarker discovery in cancer cells. Nucleic Acids Res. 2013;41:D955–61.

44. Sanij E, Hannan KM, Xuan J, Yan S, Ahern JE, Trigos AS, et al. CX-5461 activates the DNA damage response and demonstrates therapeutic efficacy in high-grade serous ovarian cancer. Nat Commun. Nature Publishing Group; 2020;11:2641.

45. Xu H, Di Antonio M, McKinney S, Mathew V, Ho B, O’Neil NJ, et al. CX-5461 is a DNA G-quadruplex stabilizer with selective lethality in BRCA1/2 deficient tumours. Nat Commun. 2017;8:14432.

46. Cornelison R, Biswas K, Llaneza DC, Harris AR, Sosale NG, Lazzara MJ, et al. CX-5461 Treatment Leads to Cytosolic DNA-Mediated STING Activation in Ovarian Cancer. Cancers. Multidisciplinary Digital Publishing Institute; 2021;13:5056.

47. Drygin D, Lin A, Bliesath J, Ho CB, O’Brien SE, Proffitt C, et al. Targeting RNA Polymerase I with an Oral Small Molecule CX-5461 Inhibits Ribosomal RNA Synthesis and Solid Tumor Growth. Cancer Res. American Association for Cancer Research; 2011;71:1418–30.

48. Haddach M, Schwaebe MK, Michaux J, Nagasawa J, O’Brien SE, Whitten JP, et al. Discovery of CX-5461, the First Direct and Selective Inhibitor of RNA Polymerase I, for Cancer Therapeutics. ACS Med Chem Lett. American Chemical Society; 2012;3:602–6.

49. Haugen BR, Alexander EK, Bible KC, Doherty GM, Mandel SJ, Nikiforov YE, et al. 2015 American Thyroid Association Management Guidelines for Adult Patients with Thyroid Nodules and Differentiated Thyroid Cancer: The American Thyroid Association Guidelines Task Force on Thyroid Nodules and Differentiated Thyroid Cancer. Thyroid Off J Am Thyroid Assoc. 2016;26:1–133.

50. Fugazzola L, Elisei R, Fuhrer D, Jarzab B, Leboulleux S, Newbold K, et al. 2019 European Thyroid Association Guidelines for the Treatment and Follow-Up of Advanced Radioiodine-Refractory Thyroid Cancer. Eur Thyroid J. Karger Publishers; 2019;8:227–45.

51. Sugitani I, Ito Y, Takeuchi D, Nakayama H, Masaki C, Shindo H, et al. Indications and Strategy for Active Surveillance of Adult Low-Risk Papillary Thyroid Microcarcinoma: Consensus Statements from the Japan Association of Endocrine Surgery Task Force on Management for Papillary Thyroid Microcarcinoma. Thyroid. 2021;31:183–92.

52. Fernández LP, López-Márquez A, Santisteban P. Thyroid transcription factors in development, differentiation and disease. Nat Rev Endocrinol. Nature Publishing Group; 2015;11:29–42.

53. Knauf JA, Sartor MA, Medvedovic M, Lundsmith E, Ryder M, Salzano M, et al. Progression of BRAF-induced thyroid cancer is associated with epithelial-mesenchymal transition requiring concomitant MAP kinase and TGFβ signaling. Oncogene. 2011;30:3153–62.

54. McFadden DG, Vernon A, Santiago PM, Martinez-McFaline R, Bhutkar A, Crowley DM, et al. p53 constrains progression to anaplastic thyroid carcinoma in a Braf-mutant mouse model of papillary thyroid cancer. Proc Natl Acad Sci U S A. 2014;111:E1600–1609.

55. Landa I, Ganly I, Chan TA, Mitsutake N, Matsuse M, Ibrahimpasic T, et al. Frequent somatic TERT promoter mutations in thyroid cancer: higher prevalence in advanced forms of the disease. J Clin Endocrinol Metab. 2013;98:E1562–1566.

56. Zhang Q, Shalaby NA, Buszczak M. Changes in rRNA transcription influence proliferation and cell fate within a stem cell lineage. Science. 2014;343:298–301.

57. Panda A, Yadav A, Yeerna H, Singh A, Biehl M, Lux M, et al. Tissue- and development-stage-specific mRNA and heterogeneous CNV signatures of human ribosomal proteins in normal and cancer samples. Nucleic Acids Res. 2020;48:7079–98.

58. Jeker LT, Hejazi M, Burek CL, Rose NR, Caturegli P. Mouse thyroid primary culture. Biochem Biophys Res Commun. 1999;257:511–5.

59. Love MI, Huber W, Anders S. Moderated estimation of fold change and dispersion for RNA-seq data with DESeq2. Genome Biol. 2014;15:550.

60. Maaten L van der, Hinton G. Visualizing Data using t-SNE. J Mach Learn Res. 2008;9:2579–605.

61. Schmidt EK, Clavarino G, Ceppi M, Pierre P. SUnSET, a nonradioactive method to monitor protein synthesis. Nat Methods. 2009;6:275–7.

