## Supplementary Figures for "TERT accelerates BRAF mutant-induced thyroid cancer dedifferentiation and progression by regulating ribosome biogenesis"

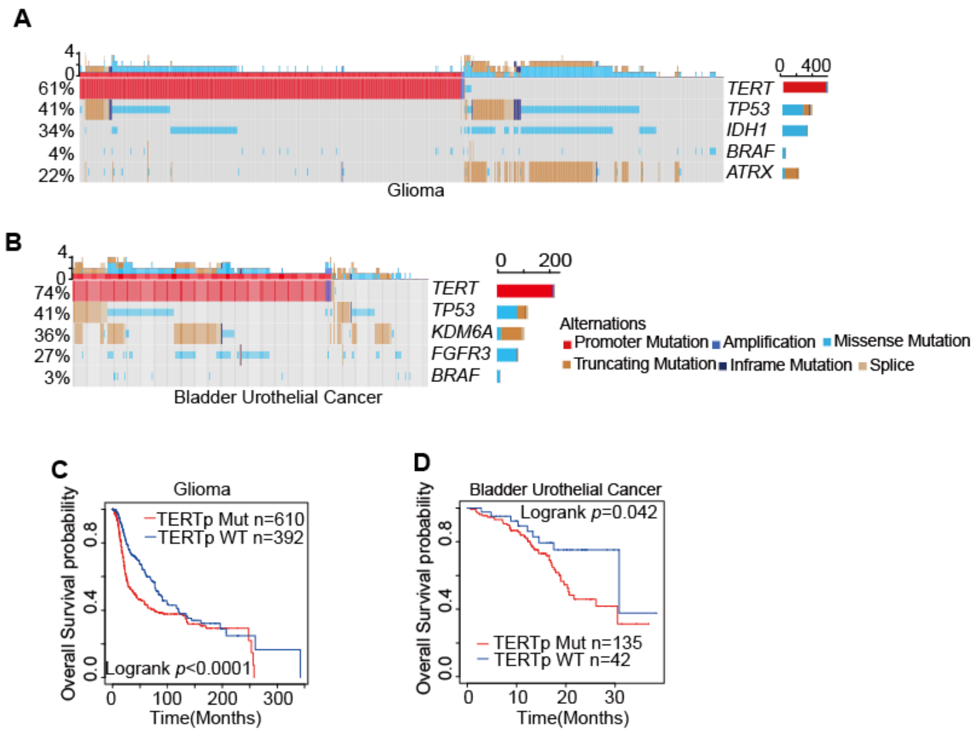


**Supplementary Figure 1. *TERT* promoter mutation occurs frequently in cancers and predicts a worse prognosis.**

A. and B. Mutation landscape of glioma and bladder urothelial cancer.

C. and D. Comparison of survival curves based on *TERT* promoter status in glioma and bladder urothelial cancer.

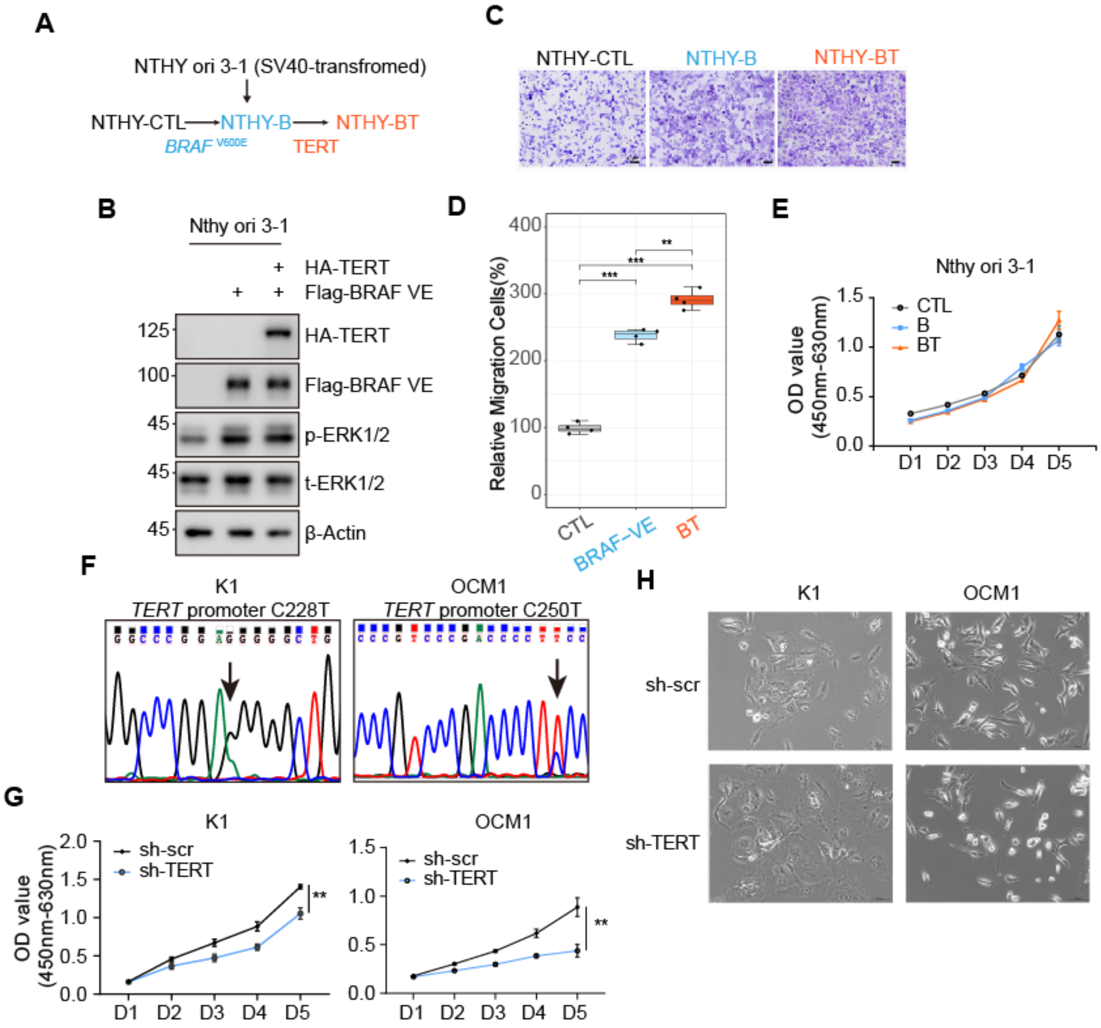


**Supplementary Figure 2. TERT partly regulated tumor cell migration and proliferation *in vitro*.**

A. Schematic diagram of stable transfected NTHY ori 3-1 cell lines construction.

B. Western blot validation of NTHY-CTL, NTHY-B, and NTHY-BT cells.

C. and D. Photomicrographs and statistic results of transwell assays of the NTHY-CTL, NTHY-B, and NTHY-BT cells. Boxplot showed the median with IQR and 1.5 IQR whiskers. **, $p < 0.01$, ***, $p < 0.001$.

E. CCK8 assay showed that NTHY-CTL, NTHY-B, and NTHY-BT cells proliferated at a similar rate.

F. Sanger sequencing showed K1 and OCM1 separately have TERT promoter C228T or C250T mutation. Arrow, mutation site.

G. CCK8 assays showed that TERT knockdown inhibited K1 and OCM1 proliferation. **, $p < 0.01$.

H. Bright field images of K1 and OCM1 cells stable infection with sh-scr or sh-TERT virus.

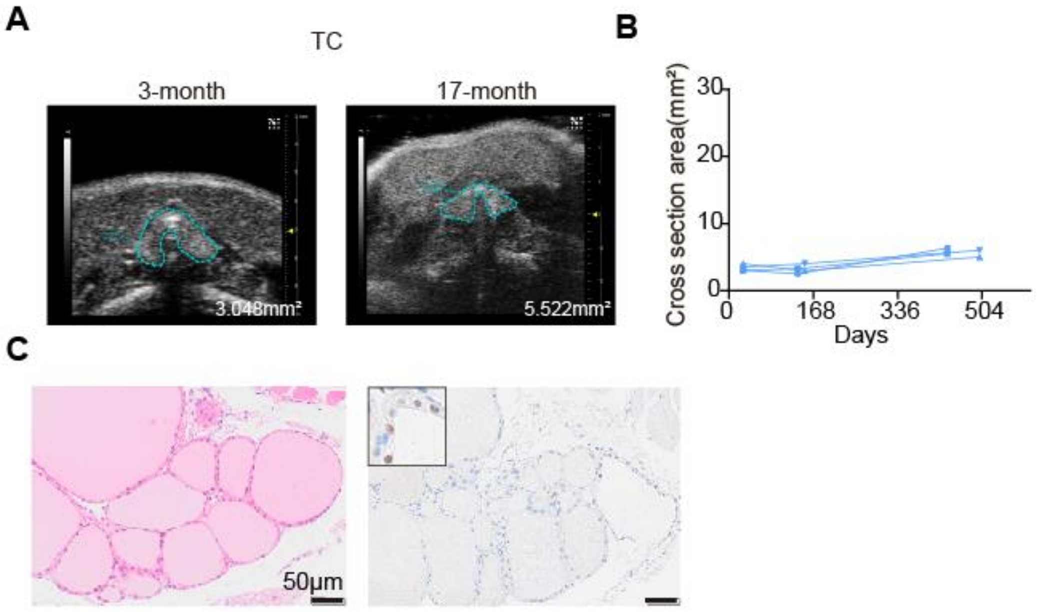


**Supplementary Figure 3. TERT alone failed to initiate tumor formation.**

A. Representative ultrasound imaging of TC mice thyroid at 3-month and 17-month old.

B. Growth patterns of TC mice thyroid (n=4).

C. H&E and anti-Flag IHC result of 17-month TC mice thyroid (scale bar, 50 μm).

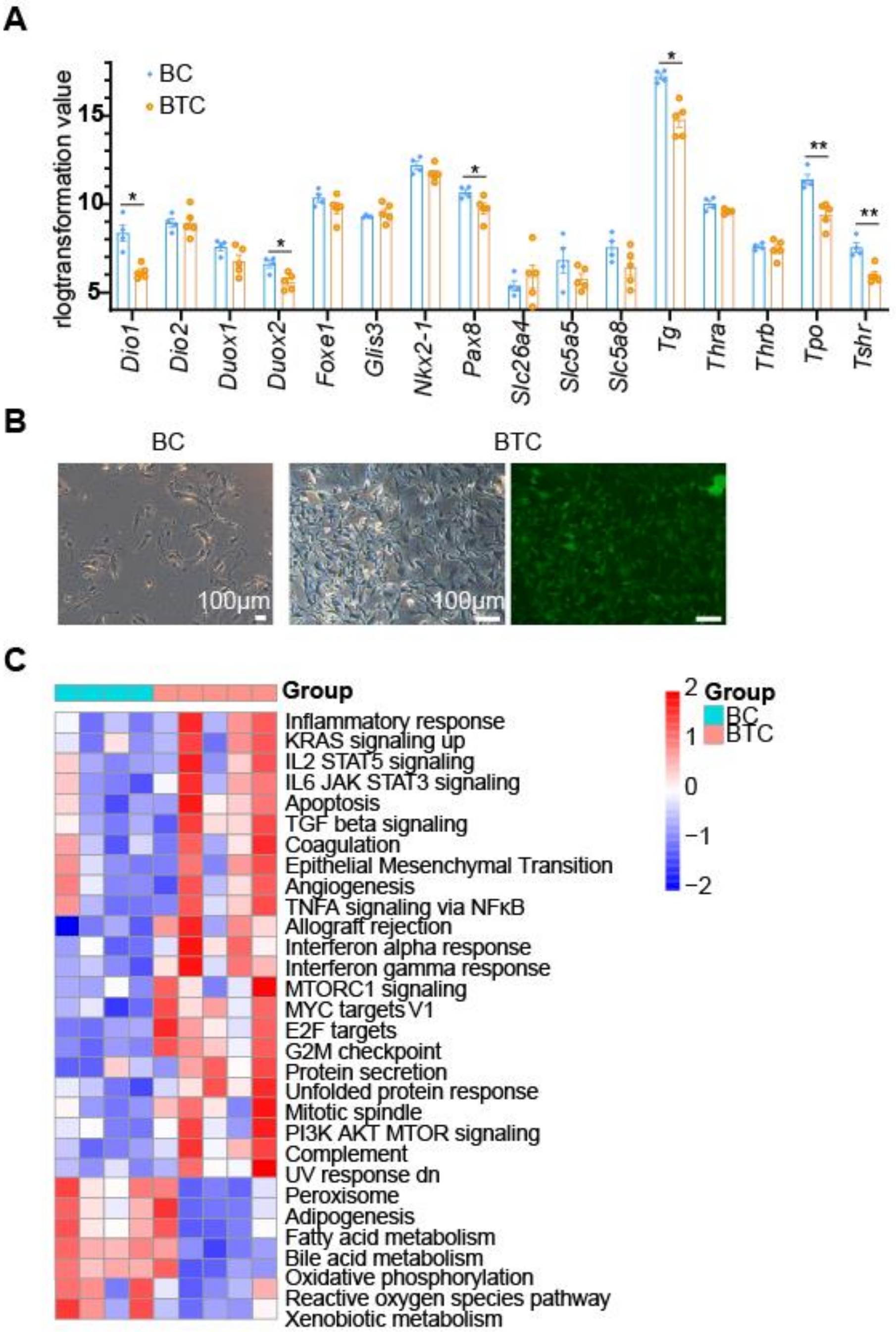


**Supplementary Figure 4. TERT promoted dedifferentiation of BRAF-induced thyroid cancer**

**A.** TDS score genes rlogtransformation value of BC and BTC thyroid RNA-seq data. Data presentation as mean±SEM.

**B.** Morphology of primary thyroid cells derived from BC and BTC thyroid in bright and fluorescence field. Scale bar, 100 μm.

**C.** Heatmap showed Hallmarks GSVA results of BC and BTC mice thyroid.

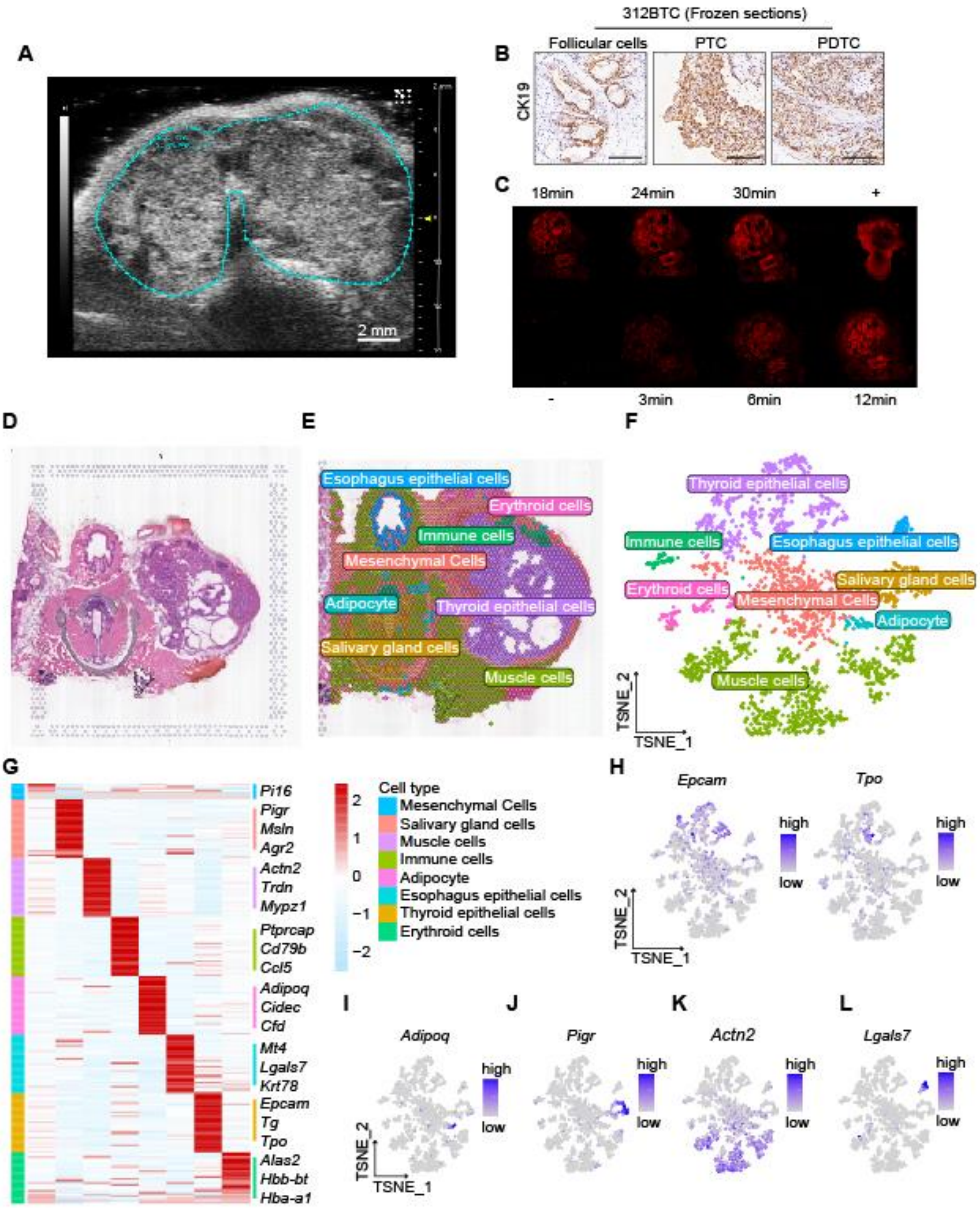


**Supplementary Figure 5. Identification of various components in 312BTC spatial transcriptome sample.**

A. Ultrasound imaging of 312BTC thyroid. Blue dashed line region, thyroid tumor.

B. Anti-CK19 IHC staining of 312BTC thyroid. Scale bar, 100 μm.

C. Optimal permeabilization time for 312BTC thyroid.

D. H&E staining of 312BTC for 10x sequencing.

E. and F. Spatial and T-SNE projections showed different cell types of all 312BTC spots.

G. Heatmap showed the relative expression levels of representative markers of different cell types.

H. I. J. K. and L. TSNE feature plot showed the representative markers of the thyroid (*Epcam, Tpo*), adipocyte (*Adipoq*), salivary gland cells (*Pigr*), muscle cells (*Actn2*) and esophagus epithelial cells (*Lgals7*).

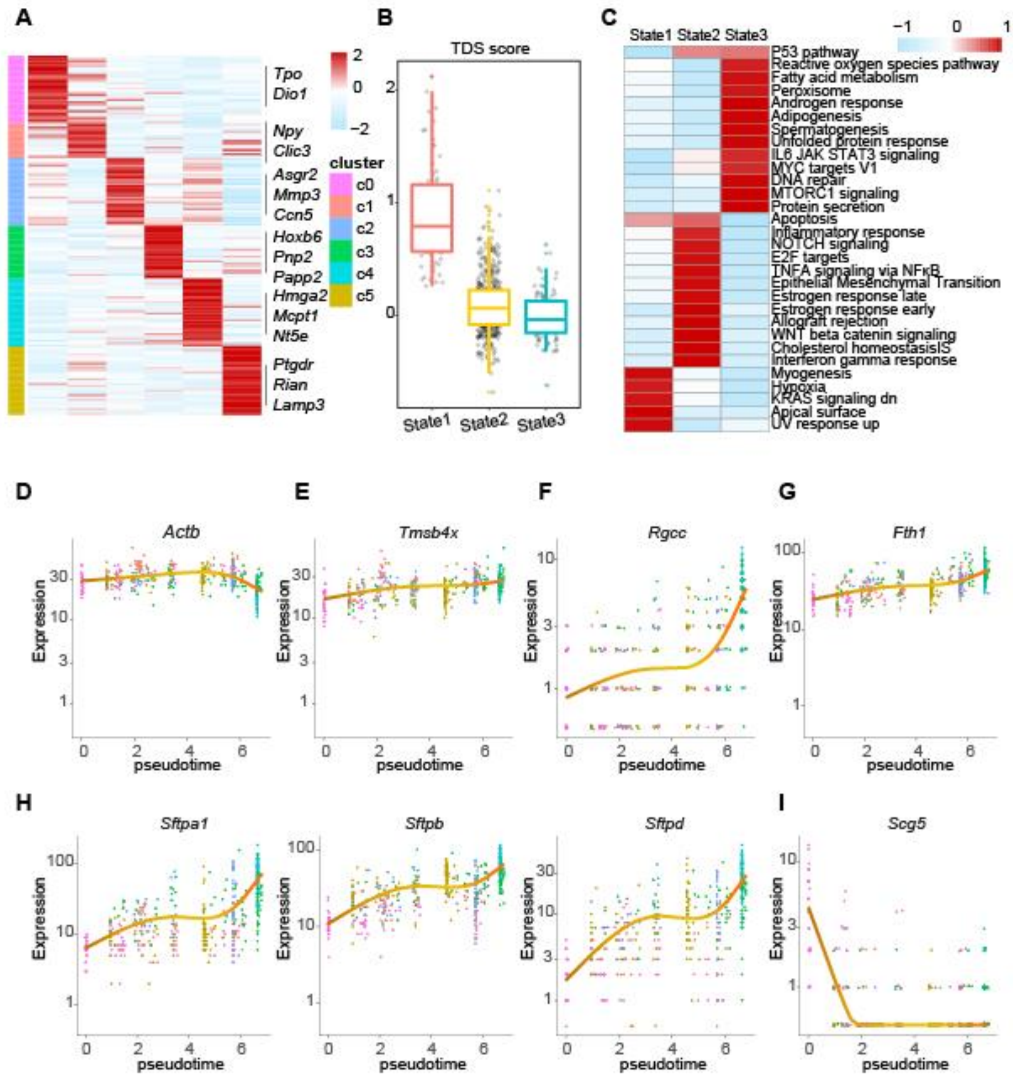


**Supplementary Figure 6. Characterization of different states of thyroid components of the spatial transcriptome.**

A. Heatmap showed representative markers of six thyroid clusters.

B. Boxplot showed the decreasing trend of TDS score from state1 to state3.

C. Heatmap presented the GSVA enrichment scores of hallmark gene sets.

D. E. F. G. H. I. Plots showed expression levels of indicated genes changed over pseudotime.

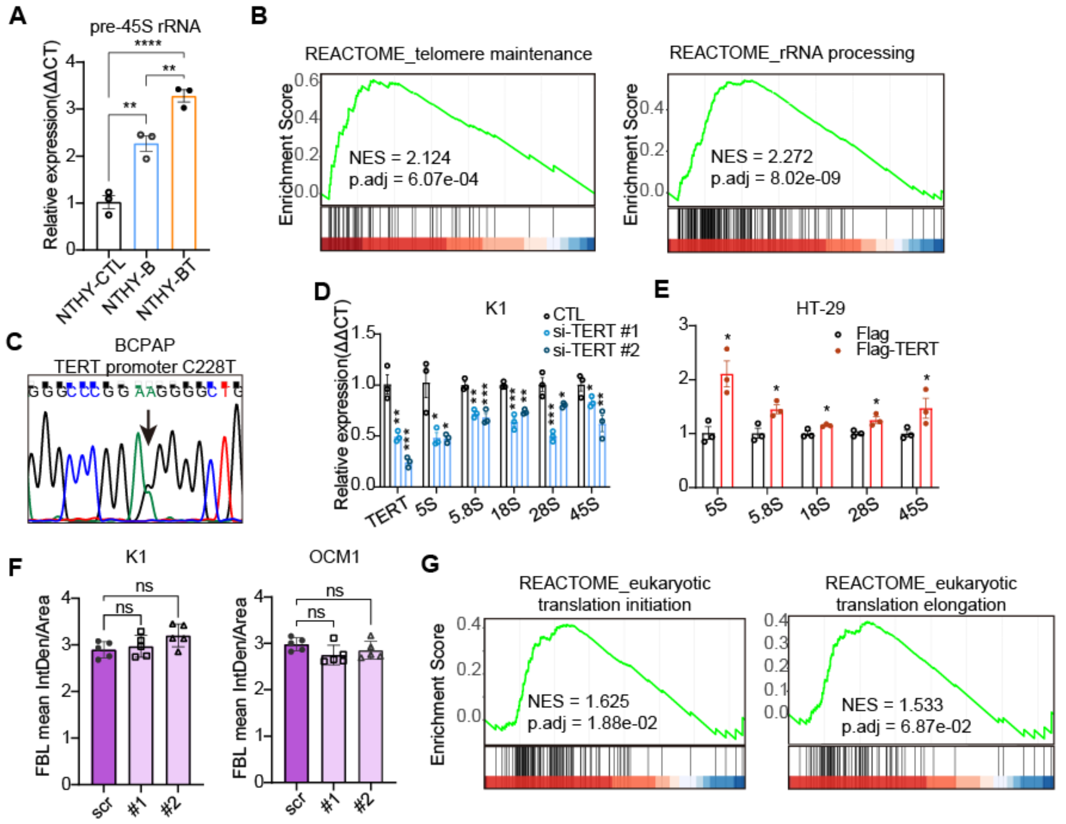


**Supplementary Figure 7. TERT regulated ribosome-related functions.**

**A.** QPCR results of pre-45S rRNA in NTHY-CTL, NTHY-B, and NTHY-BT cells. Data presented as mean±SEM (n=3). **, $p$<0.01; ****, $p$<0.0001.

**B.** GSEA results showed TERT high-expression cells were positively enriched in telomere maintenance and rRNA metabolic process in the CCLE dataset.

**C.** Sanger sequencing result showed *TERT* promoter C228T mutation site in BCPAP.

**D.** and **E.** QPCR results of ribosomal RNA 5S, 5.8S, 18S, 28S and 45S change after TERT knockdown (D, in K1 cells) and TERT over-expression (E, in HT-29 cells). Data presented as mean±SD (n=5). *, $p$<0.05; **, $p$<0.01; ***, $p$<0.001.

**F.** Quantification results of fig.4F and fig.4H FBL results using ImageJ software (n=5).

**G.** GSEA results showed TERT high-expression cells were positively enriched in translation initiation and translation elongation.

A

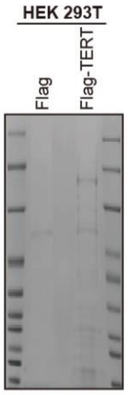


B

| | Gene ID | CTL #PSM | Flag-TERT #PSM |
|---|---|---|---|
| rRNA processing | BYSL | 0 | 7 |
| | EXOSC10 | 0 | 13 |
| | EXOSC4 | 0 | 6 |
| | EXOSC2 | 0 | 3 |
| translation | EIF4A3 | 0 | 11 |
| | EIF2S1 | 0 | 7 |
| | EIF2S2 | 0 | 5 |
| telomerase | PES1 | 0 | 4 |
| | DKC1 | 0 | 5 |

**Supplementary Figure 8. Identification of TERT-interacting proteins**

A. SDS-PAGE of lysate immunoprecipitated with anti-Flag antibody showed the staining result of coomassie brilliant blue and rendered in grayscale.

B. Representative proteins identified by IP/LC-MS. PSM, peptide spectra match.
