## Supplementary Tables for "TERT accelerates BRAF mutant-induced thyroid cancer dedifferentiation and progression by regulating ribosome biogenesis"

**Supplementary table S1:**

| **Genotyping primers (5' to 3')** | |
|---|---|
| Rs-mTert-F | TCAGATTCTTTTATAGGGGACACA |
| Rs-mTert-R1 | GGTGTTGTCGGGGAAATCATCGTC |
| Rs-mTert-R2 | AGGAGCCTGCCAAGTAAC |

| **Cloning primers (5' to 3')** | |
|---|---|
| hTERT-F | ctgttccaggggcccaccggtCCGCGCGCTCCCCGCTGC |
| hTERT-R | ctccagcgaattggcgaattcTCAGTCCAGGATGGTCTTGAAGT |
| hBRAF-F | ccaggggcccaccggtATGGCGGCGCTGAGCGGTG |
| hBRAF-R | gcgaattggcgaattcCTACTTGAAGGCTGCAAATT |
| EIF2S1-F | ctgttccaggggcccaccggtATGCCGGGTCTAAGTTGTAGATTT |
| EIF2S1-R | ctccagcgaattggcgaattcTTAATCTTCAGCTTTGGCTTCCA |
| EIF2S2-F | ctgttccaggggcccaccggtATGTCTGGGGACGAGATGATTT |
| EIF2S2-R | ctccagcgaattggcgaattcTTAGTTAGCTTTGGCACGGAGC |
| POLR1C-F | ctgttccaggggcccaccggtATGGCGGCTTCTCAGGCG |
| POLR1C-R | ctccagcgaattggcgaattcTCAGTCCATCTGAACCGCATC |
| EXOSC2-F | ctgttccaggggcccaccggtATGGCGATGGAGATGAGGC |
| EXOSC2-R | ctccagcgaattggcgaattcTTATCCCTCCTGTTCCAAAAGC |
| EXOSC7-F | ctgttccaggggcccaccggtATGGCGTCCGTGACGCTG |
| EXOSC7-R | ctccagcgaattggcgaattcTCATCCCAGGAATCCAACTTTC |
| EXOSC8-F | ctgttccaggggcccaccggtATGGCGGCTGGGTTCAAA |
| EXOSC8-R | ctccagcgaattggcgaattcTTATTTGGGTTTCATACTCTTAATTACTTC |

| **siRNA/shRNA sequence (5' to 3')** | |
|---|---|
| Negative control | UUCUCCGAACGUGUCACGUTT |
| | ACGUGACACGUUCGGAGAATT |
| TERT（human） | CGAAGAACGUGCUGGCCUUTT |
| siRNA#1 | AAGGCCAGCACGUUCUUCGTT |
| TERT（human） | AGAACGUUCCGCAGAGAAATT |
| siRNA#3 | UUUCUCUGCGGAACGUUCUTT |
| TERC（human） | GUCUAACCCUAACUGAGAATT |
| siRNA#1 | UUCUCAGUUAGGGUUAGACTT |
| TERC（human） | CGUUCAUUCUAGAGCAAACTT |
| siRNA#2 | GUUUGCUCUAGAAUGAACGTT |
| TERC（human） | AGGGCGAGGUUCAGGCCUUTT |
| siRNA#3 | AAGGCCUGAACCUCGCCCUTT |
| shTERT_1_F | CCGGGACATGGAGAACAAGCTGTTTCTCGAGAAACAGCT |
| | TGTTCTCCATGTCTTTTTTG |
| shTERT_1_R | AATTCAAAAAAGACATGGAGAACAAGCTGTTTCTCGAGA |
| | AACAGCTTGTTCTCCATGTC |
| shTERT_2_F | CCGGGAAGAGTGTCTGGAGCAAGTTCTCGAGAACTTGCT |
| | CCAGACACTCTTCTTTTTTG |

| | |
|---|---|
| shTERT_2_R | AATTCAAAAAAGAAGAGTGTCTGGAGCAAGTTCTCGAGA ACTTGCTCCAGACACTCTTC |

**Supplementary table S2:**

| **Antibodies** | **Cat** | **Dilution** |
|---|---|---|
| Mouse monoclonal anti-Flag Tag (HRP conjugated) | Sigma-Aldrich, Cat#A8592; RRID: AB_439702 | 1:150 (IHC), 1:10000 (WB) |
| Rabbit monoclonal anti-Ha Tag (C29F4) (HRP Conjugate) | Cell Signaling Technology,Cat#14031S; RRID: AB_2798368 | 1:10000 (WB) |
| Rabbit monoclonal anti-Phospho-p44/42 MAPK (Erk1/2) (Thr202/Tyr204) (D13.14.4E) | Cell Signaling Technology, Cat#4370 | 1: 3000 (WB) |
| Rabbit monoclonal anti-p44/42 MAPK (Erk1/2) (137F5) | Cell Signaling Technology, Cat#4695 | 1:2000 (WB) |
| Mouse Monoclonal anti-Beta Actin | Proteintech, Cat#66009-1-Ig | 1:10000 (WB) |
| Rabbit monoclonal anti-cytokeratin 19 (EP1580Y) | Abcam, Cat#ab52625;RRID: AB_2281020 | 1:600 (IHC) |
| Rabbit monoclonal anti-Thyroglobulin (EPR9730) | Abcam, Cat#ab156008 | 1:500 (IHC) |
| Rabbit monoclonal anti-TTF-1(EP1584Y) | Abcam, Cat#ab76013 | 1:500 (IHC) |
| Rabbit polyclonal anti-PAX8 | Proteintech, Cat#10336-1-AP | 1:2000 (IHC) |
| Rabbit polyclonal anti-Ki67 | Abcam,Cat#ab15580; RRID: AB_443209 | 1:800 (IHC) |
| Rabbit polyclonal anti-Sodium iodide symporter | Proteintech, Cat #24324-1-AP | 1:400 (IHC) 1:1000 (WB) |
| Rabbit monoclonal anti-Prosurfactant (EPR19839) | Abcam, Cat#ab211326 | 1:500 (IHC) |
| Rabbit polyclonal anti-Fibrillarin | Abcam, Cat#ab5821 | 1:1000 (IF) |
| Mouse monoclonal anti-Puromycin (12D10) | Sigma, Cat#MABE343 | 1:1500 (WB) |
| Rabbit polyclonal anti-EIF2S2 | Proteintech, Cat #10227-1-AP | 1:2000 (WB) |
| Rabbit polyclonal anti-BYSL | Proteintech, Cat #28319-1-AP | 1:5000 (WB) |
| Rabbit monoclonal anti-Phospho-S6 Ribosomal Protein (Ser235/236)(D57.2.2E) | Cell Signaling Technology, Cat#4858 | 1:600 (IHC), 1:2000 (WB) |
| Rabbit polyclonal anti-Phospho-p70 S6 Kinase (Thr389) | Cell Signaling Technology, Cat#9205 | 1:2000 (WB) |
| Rabbit polyclonal anti-RPS6 | ServiceBio, Cat#GB112610 | 1:1000 (WB) |
| Rabbit polyclonal anti-p70 S6 Kinase | Cell Signaling Technology, Cat#9202 | 1:1000 (WB) |
| Rabbit monoclonal anti-Vinculin (E1E9V) | Cell Signaling Technology, Cat#13901S; RRID: AB_2728768 | 1:3000 (WB) |
| Peroxidase AffiniPure Goat AntiMouse IgG (H+L) | Jackson ImmunoResearch Laboratories, Cat#115-035-003; RRID: AB_10015289 | |
| Peroxidase AffiniPure Goat AntiRabbit IgG (H+L) | Jackson ImmunoResearch Laboratories, Cat#111-035-003; RRID: AB_2313567 | |

**Supplementary table S3:**

| Accession | #Spec Elu-CTL-2 | #Spec Elu-F-T-2 | Description |
|---|---|---|---|
| O14746 | 0 | 568 | Telomerase reverse transcriptase OS=Homo sapiens OX=9606 GN=TERT PE=1 SV=1 |
| Q00839 | 0 | 109 | Heterogeneous nuclear ribonucleoprotein U OS=Homo sapiens OX=9606 GN=HNRNPU PE=1 SV=6 |
| P55795 | 0 | 68 | Heterogeneous nuclear ribonucleoprotein H2 OS=Homo sapiens OX=9606 GN=HNRNPH2 PE=1 SV=1 |
| P10412 | 0 | 62 | Histone H1.4 OS=Homo sapiens OX=9606 GN=HIST1H1E PE=1 SV=2 |
| P16402 | 0 | 61 | Histone H1.3 OS=Homo sapiens OX=9606 GN=HIST1H1D PE=1 SV=2 |
| Q12906 | 0 | 55 | Interleukin enhancer-binding factor 3 OS=Homo sapiens OX=9606 GN=ILF3 PE=1 SV=3 |
| Q13247 | 0 | 54 | Serine/arginine-rich splicing factor 6 OS=Homo sapiens OX=9606 GN=SRSF6 PE=1 SV=2 |
| Q07955 | 0 | 53 | Serine/arginine-rich splicing factor 1 OS=Homo sapiens OX=9606 GN=SRSF1 PE=1 SV=2 |
| O43390 | 0 | 40 | Heterogeneous nuclear ribonucleoprotein R OS=Homo sapiens OX=9606 GN=HNRNPR PE=1 SV=1 |
| P14866 | 0 | 39 | Heterogeneous nuclear ribonucleoprotein L OS=Homo sapiens OX=9606 GN=HNRNPL PE=1 SV=2 |
| Q99729 | 0 | 37 | Heterogeneous nuclear ribonucleoprotein A/B OS=Homo sapiens OX=9606 GN=HNRNPAB PE=1 SV=2 |
| Q14103 | 0 | 35 | Heterogeneous nuclear ribonucleoprotein D0 OS=Homo sapiens OX=9606 GN=HNRNPD PE=1 SV=1 |
| Q8IWS0 | 0 | 35 | PHD finger protein 6 OS=Homo sapiens OX=9606 GN=PHF6 PE=1 SV=1 |
| Q08170 | 0 | 35 | Serine/arginine-rich splicing factor 4 OS=Homo sapiens OX=9606 GN=SRSF4 PE=1 SV=2 |
| Q9H6T3 | 0 | 34 | RNA polymerase II-associated protein 3 OS=Homo sapiens OX=9606 GN=RPAP3 PE=1 SV=2 |
| O60506 | 0 | 34 | Heterogeneous nuclear ribonucleoprotein Q OS=Homo sapiens OX=9606 GN=SYNCRIP PE=1 SV=2 |
| Q9Y3U8 | 0 | 32 | 60S ribosomal protein L36 OS=Homo sapiens OX=9606 GN=RPL36 PE=1 SV=3 |
| Q9Y3F4 | 0 | 31 | Serine-threonine kinase receptor-associated protein OS=Homo sapiens OX=9606 GN=STRAP PE=1 SV=1 |
| P43243 | 0 | 30 | Matrin-3 OS=Homo sapiens OX=9606 GN=MATR3 PE=1 SV=2 |
| Q08211 | 0 | 29 | ATP-dependent RNA helicase A OS=Homo sapiens OX=9606 GN=DHX9 PE=1 SV=4 |
| P46087 | 0 | 27 | Probable 28S rRNA (cytosine(4447)-C(5))-methyltransferase OS=Homo sapiens OX=9606 GN=NOP2 PE=1 SV=2 |
| P62910 | 0 | 27 | 60S ribosomal protein L32 OS=Homo sapiens OX=9606 GN=RPL32 PE=1 SV=2 |
| P51114 | 0 | 23 | Fragile X mental retardation syndrome-related protein 1 OS=Homo sapiens OX=9606 GN=FXR1 PE=1 SV=3 |
| Q7L2E3 | 0 | 21 | ATP-dependent RNA helicase DHX30 OS=Homo sapiens OX=9606 GN=DHX30 PE=1 SV=1 |
| Q7L2J0 | 0 | 20 | 7SK snRNA methylphosphate capping enzyme OS=Homo sapiens OX=9606 GN=MEPCE PE=1 SV=1 |
| Q01130 | 0 | 20 | Serine/arginine-rich splicing factor 2 OS=Homo sapiens OX=9606 GN=SRSF2 PE=1 SV=4 |

| | | | |
|---|---|---|---|
| Q9BVP2 | 0 | 19 | Guanine nucleotide-binding protein-like 3 OS=Homo sapiens OX=9606 GN=GNL3 PE=1 SV=2 |
| Q13243 | 0 | 19 | Serine/arginine-rich splicing factor 5 OS=Homo sapiens OX=9606 GN=SRSF5 PE=1 SV=1 |
| Q9HCE1 | 0 | 18 | Helicase MOV-10 OS=Homo sapiens OX=9606 GN=MOV10 PE=1 SV=2 |
| Q14258 | 0 | 18 | E3 ubiquitin/ISG15 ligase TRIM25 OS=Homo sapiens OX=9606 GN=TRIM25 PE=1 SV=2 |
| Q13435 | 0 | 17 | Splicing factor 3B subunit 2 OS=Homo sapiens OX=9606 GN=SF3B2 PE=1 SV=2 |
| O15355 | 0 | 16 | Protein phosphatase 1G OS=Homo sapiens OX=9606 GN=PPM1G PE=1 SV=1 |
| Q71RC2 | 0 | 15 | La-related protein 4 OS=Homo sapiens OX=9606 GN=LARP4 PE=1 SV=3 |
| P31942 | 0 | 15 | Heterogeneous nuclear ribonucleoprotein H3 OS=Homo sapiens OX=9606 GN=HNRNPH3 PE=1 SV=2 |
| P60866 | 0 | 15 | 40S ribosomal protein S20 OS=Homo sapiens OX=9606 GN=RPS20 PE=1 SV=1 |
| Q86V81 | 0 | 14 | THO complex subunit 4 OS=Homo sapiens OX=9606 GN=ALYREF PE=1 SV=3 |
| P55265 | 0 | 14 | Double-stranded RNA-specific adenosine deaminase OS=Homo sapiens OX=9606 GN=ADAR PE=1 SV=4 |
| P31948 | 0 | 14 | Stress-induced-phosphoprotein 1 OS=Homo sapiens OX=9606 GN=STIP1 PE=1 SV=1 |
| Q9Y383 | 0 | 14 | Putative RNA-binding protein Luc7-like 2 OS=Homo sapiens OX=9606 GN=LUC7L2 PE=1 SV=2 |
| Q99733 | 0 | 14 | Nucleosome assembly protein 1-like 4 OS=Homo sapiens OX=9606 GN=NAP1L4 PE=1 SV=1 |
| Q9BQG0 | 0 | 14 | Myb-binding protein 1A OS=Homo sapiens OX=9606 GN=MYBBP1A PE=1 SV=2 |
| Q9Y4P3 | 0 | 14 | Transducin beta-like protein 2 OS=Homo sapiens OX=9606 GN=TBL2 PE=1 SV=1 |
| P62266 | 0 | 13 | 40S ribosomal protein S23 OS=Homo sapiens OX=9606 GN=RPS23 PE=1 SV=3 |
| Q01780 | 0 | 13 | Exosome component 10 OS=Homo sapiens OX=9606 GN=EXOSC10 PE=1 SV=2 |
| Q07666 | 0 | 13 | KH domain-containing RNA-binding signal transduction-associated protein 1 OS=Homo sapiens OX=9606 GN=KHDRBS1 PE=1 SV=1 |
| Q92499 | 0 | 13 | ATP-dependent RNA helicase DDX1 OS=Homo sapiens OX=9606 GN=DDX1 PE=1 SV=2 |
| P62854 | 0 | 13 | 40S ribosomal protein S26 OS=Homo sapiens OX=9606 GN=RPS26 PE=1 SV=3 |
| O15226 | 0 | 12 | NF-kappa-B-repressing factor OS=Homo sapiens OX=9606 GN=NKRF PE=1 SV=2 |
| P32969 | 0 | 12 | 60S ribosomal protein L9 OS=Homo sapiens OX=9606 GN=RPL9 PE=1 SV=1 |
| Q9UN86 | 0 | 12 | Ras GTPase-activating protein-binding protein 2 OS=Homo sapiens OX=9606 GN=G3BP2 PE=1 SV=2 |
| Q14444 | 0 | 12 | Caprin-1 OS=Homo sapiens OX=9606 GN=CAPRIN1 PE=1 SV=2 |
| P62244 | 0 | 12 | 40S ribosomal protein S15a OS=Homo sapiens OX=9606 GN=RPS15A PE=1 SV=2 |
| Q12905 | 0 | 11 | Interleukin enhancer-binding factor 2 OS=Homo sapiens OX=9606 GN=ILF2 PE=1 SV=2 |
| Q9GZR7 | 0 | 11 | ATP-dependent RNA helicase DDX24 OS=Homo sapiens OX=9606 GN=DDX24 PE=1 SV=1 |
| Q7Z417 | 0 | 11 | Nuclear fragile X mental retardation-interacting protein 2 OS=Homo sapiens OX=9606 GN=NUFIP2 PE=1 SV=1 |
| P38919 | 0 | 11 | Eukaryotic initiation factor 4A-III OS=Homo sapiens OX=9606 GN=EIF4A3 PE=1 SV=4 |

| | | | |
|---|---|---|---|
| Q99459 | 0 | 11 | Cell division cycle 5-like protein OS=Homo sapiens OX=9606 GN=CDC5L PE=1 SV=2 |
| Q06787 | 0 | 11 | Synaptic functional regulator FMR1 OS=Homo sapiens OX=9606 GN=FMR1 PE=1 SV=1 |
| Q9UJV9 | 0 | 11 | Probable ATP-dependent RNA helicase DDX41 OS=Homo sapiens OX=9606 GN=DDX41 PE=1 SV=2 |
| P82650 | 0 | 11 | 28S ribosomal protein S22 mitochondrial OS=Homo sapiens OX=9606 GN=MRPS22 PE=1 SV=1 |
| Q96PK6 | 0 | 10 | RNA-binding protein 14 OS=Homo sapiens OX=9606 GN=RBM14 PE=1 SV=2 |
| Q9P258 | 0 | 10 | Protein RCC2 OS=Homo sapiens OX=9606 GN=RCC2 PE=1 SV=2 |
| P51116 | 0 | 10 | Fragile X mental retardation syndrome-related protein 2 OS=Homo sapiens OX=9606 GN=FXR2 PE=1 SV=2 |
| Q96KR1 | 0 | 9 | Zinc finger RNA-binding protein OS=Homo sapiens OX=9606 GN=ZFR PE=1 SV=2 |
| Q07065 | 0 | 9 | Cytoskeleton-associated protein 4 OS=Homo sapiens OX=9606 GN=CKAP4 PE=1 SV=2 |
| Q9H5H4 | 0 | 9 | Zinc finger protein 768 OS=Homo sapiens OX=9606 GN=ZNF768 PE=1 SV=2 |
| Q8NE71 | 0 | 9 | ATP-binding cassette sub-family F member 1 OS=Homo sapiens OX=9606 GN=ABCF1 PE=1 SV=2 |
| Q7KZF4 | 0 | 9 | Staphylococcal nuclease domain-containing protein 1 OS=Homo sapiens OX=9606 GN=SND1 PE=1 SV=1 |
| Q9Y2X3 | 0 | 9 | Nucleolar protein 58 OS=Homo sapiens OX=9606 GN=NOP58 PE=1 SV=1 |
| Q08J23 | 0 | 9 | tRNA (cytosine(34)-C(5))-methyltransferase OS=Homo sapiens OX=9606 GN=NSUN2 PE=1 SV=2 |
| Q9BXP5 | 0 | 9 | Serrate RNA effector molecule homolog OS=Homo sapiens OX=9606 GN=SRRT PE=1 SV=1 |
| P42285 | 0 | 9 | Exosome RNA helicase MTR4 OS=Homo sapiens OX=9606 GN=MTREX PE=1 SV=3 |
| Q8IY81 | 0 | 8 | pre-rRNA 2'-O-ribose RNA methyltransferase FTSJ3 OS=Homo sapiens OX=9606 GN=FTSJ3 PE=1 SV=2 |
| Q92522 | 0 | 8 | Histone H1x OS=Homo sapiens OX=9606 GN=H1FX PE=1 SV=1 |
| Q5BKZ1 | 0 | 8 | DBIRD complex subunit ZNF326 OS=Homo sapiens OX=9606 GN=ZNF326 PE=1 SV=2 |
| Q9NWK9 | 0 | 8 | Box C/D snoRNA protein 1 OS=Homo sapiens OX=9606 GN=ZNHIT6 PE=1 SV=1 |
| Q9NZB2 | 0 | 8 | Constitutive coactivator of PPAR-gamma-like protein 1 OS=Homo sapiens OX=9606 GN=FAM120A PE=1 SV=2 |
| Q01081 | 0 | 8 | Splicing factor U2AF 35 kDa subunit OS=Homo sapiens OX=9606 GN=U2AF1 PE=1 SV=3 |
| P0DN76 | 0 | 8 | Splicing factor U2AF 35 kDa subunit-like protein OS=Homo sapiens OX=9606 GN=U2AF1L5 PE=1 SV=1 |
| Q92665 | 0 | 8 | 28S ribosomal protein S31 mitochondrial OS=Homo sapiens OX=9606 GN=MRPS31 PE=1 SV=3 |
| Q9BYD6 | 0 | 8 | 39S ribosomal protein L1 mitochondrial OS=Homo sapiens OX=9606 GN=MRPL1 PE=1 SV=2 |
| Q8NB90 | 0 | 8 | ATPase family protein 2 homolog OS=Homo sapiens OX=9606 GN=SPATA5 PE=1 SV=3 |
| Q6P5R6 | 0 | 8 | 60S ribosomal protein L22-like 1 OS=Homo sapiens OX=9606 GN=RPL22L1 PE=1 SV=2 |
| Q92552 | 0 | 7 | 28S ribosomal protein S27 mitochondrial OS=Homo sapiens OX=9606 GN=MRPS27 PE=1 SV=3 |
| O95232 | 0 | 7 | Luc7-like protein 3 OS=Homo sapiens OX=9606 GN=LUC7L3 PE=1 SV=2 |
| Q6NZY4 | 0 | 7 | Zinc finger CCHC domain-containing protein 8 OS=Homo sapiens OX=9606 GN=ZCCHC8 PE=1 SV=2 |
| Q9H7E9 | 0 | 7 | UPF0488 protein C8orf33 OS=Homo sapiens OX=9606 GN=C8orf33 PE=1 SV=1 |

| | | | |
|---|---|---|---|
| Q9HBE1 | 0 | 7 | POZ- AT hook- and zinc finger-containing protein 1 OS=Homo sapiens OX=9606 GN=PATZ1 PE=1 SV=1 |
| P82933 | 0 | 7 | 28S ribosomal protein S9 mitochondrial OS=Homo sapiens OX=9606 GN=MRPS9 PE=1 SV=2 |
| Q8N5U6 | 0 | 7 | RING finger protein 10 OS=Homo sapiens OX=9606 GN=RNF10 PE=1 SV=2 |
| P49756 | 0 | 7 | RNA-binding protein 25 OS=Homo sapiens OX=9606 GN=RBM25 PE=1 SV=3 |
| Q13242 | 0 | 7 | Serine/arginine-rich splicing factor 9 OS=Homo sapiens OX=9606 GN=SRSF9 PE=1 SV=1 |
| Q7L0Y3 | 0 | 7 | tRNA methyltransferase 10 homolog C OS=Homo sapiens OX=9606 GN=TRMT10C PE=1 SV=2 |
| Q6PKG0 | 0 | 7 | La-related protein 1 OS=Homo sapiens OX=9606 GN=LARP1 PE=1 SV=2 |
| P82914 | 0 | 7 | 28S ribosomal protein S15 mitochondrial OS=Homo sapiens OX=9606 GN=MRPS15 PE=1 SV=1 |
| Q13895 | 0 | 7 | Bystin OS=Homo sapiens OX=9606 GN=BYSL PE=1 SV=3 |
| Q96SI9 | 0 | 7 | Spermatid perinuclear RNA-binding protein OS=Homo sapiens OX=9606 GN=STRBP PE=1 SV=1 |
| O95373 | 0 | 7 | Importin-7 OS=Homo sapiens OX=9606 GN=IPO7 PE=1 SV=1 |
| Q16629 | 0 | 7 | Serine/arginine-rich splicing factor 7 OS=Homo sapiens OX=9606 GN=SRSF7 PE=1 SV=1 |
| P78362 | 0 | 7 | SRSF protein kinase 2 OS=Homo sapiens OX=9606 GN=SRPK2 PE=1 SV=3 |
| Q1KMD3 | 0 | 7 | Heterogeneous nuclear ribonucleoprotein U-like protein 2 OS=Homo sapiens OX=9606 GN=HNRNPUL2 PE=1 SV=1 |
| P63151 | 0 | 7 | Serine/threonine-protein phosphatase 2A 55 kDa regulatory subunit B alpha isoform OS=Homo sapiens OX=9606 GN=PPP2R2A PE=1 SV=1 |
| Q96EY7 | 0 | 7 | Pentatricopeptide repeat domain-containing protein 3 mitochondrial OS=Homo sapiens OX=9606 GN=PTCD3 PE=1 SV=3 |
| P62888 | 0 | 7 | 60S ribosomal protein L30 OS=Homo sapiens OX=9606 GN=RPL30 PE=1 SV=2 |
| O75400 | 0 | 7 | Pre-mRNA-processing factor 40 homolog A OS=Homo sapiens OX=9606 GN=PRPF40A PE=1 SV=2 |
| Q9HC36 | 0 | 7 | rRNA methyltransferase 3 mitochondrial OS=Homo sapiens OX=9606 GN=MRM3 PE=1 SV=2 |
| Q13283 | 0 | 7 | Ras GTPase-activating protein-binding protein 1 OS=Homo sapiens OX=9606 GN=G3BP1 PE=1 SV=1 |
| Q92615 | 0 | 7 | La-related protein 4B OS=Homo sapiens OX=9606 GN=LARP4B PE=1 SV=3 |
| P05198 | 0 | 7 | Eukaryotic translation initiation factor 2 subunit 1 OS=Homo sapiens OX=9606 GN=EIF2S1 PE=1 SV=3 |
| P08708 | 0 | 7 | 40S ribosomal protein S17 OS=Homo sapiens OX=9606 GN=RPS17 PE=1 SV=2 |
| P52298 | 0 | 7 | Nuclear cap-binding protein subunit 2 OS=Homo sapiens OX=9606 GN=NCBP2 PE=1 SV=1 |
| Q9NYF8 | 0 | 6 | Bcl-2-associated transcription factor 1 OS=Homo sapiens OX=9606 GN=BCLAF1 PE=1 SV=2 |
| O43684 | 0 | 6 | Mitotic checkpoint protein BUB3 OS=Homo sapiens OX=9606 GN=BUB3 PE=1 SV=1 |
| O43823 | 0 | 6 | A-kinase anchor protein 8 OS=Homo sapiens OX=9606 GN=AKAP8 PE=1 SV=1 |
| Q6UXN9 | 0 | 6 | WD repeat-containing protein 82 OS=Homo sapiens OX=9606 GN=WDR82 PE=1 SV=1 |
| Q8WU90 | 0 | 6 | Zinc finger CCCH domain-containing protein 15 OS=Homo sapiens OX=9606 GN=ZC3H15 PE=1 SV=1 |
| Q9BRJ6 | 0 | 6 | Uncharacterized protein C7orf50 OS=Homo sapiens OX=9606 GN=C7orf50 PE=1 SV=1 |

| | | | |
|---|---|---|---|
| P28074 | 0 | 6 | Proteasome subunit beta type-5 OS=Homo sapiens OX=9606 GN=PSMB5 PE=1 SV=3 |
| P23246 | 0 | 6 | Splicing factor proline- and glutamine-rich OS=Homo sapiens OX=9606 GN=SFPQ PE=1 SV=2 |
| Q4G0J3 | 0 | 6 | La-related protein 7 OS=Homo sapiens OX=9606 GN=LARP7 PE=1 SV=1 |
| Q9NW13 | 0 | 6 | RNA-binding protein 28 OS=Homo sapiens OX=9606 GN=RBM28 PE=1 SV=3 |
| Q2NL82 | 0 | 6 | Pre-rRNA-processing protein TSR1 homolog OS=Homo sapiens OX=9606 GN=TSR1 PE=1 SV=1 |
| P42696 | 0 | 6 | RNA-binding protein 34 OS=Homo sapiens OX=9606 GN=RBM34 PE=1 SV=2 |
| Q9Y3I0 | 0 | 6 | tRNA-splicing ligase RtcB homolog OS=Homo sapiens OX=9606 GN=RTCB PE=1 SV=1 |
| P57678 | 0 | 6 | Gem-associated protein 4 OS=Homo sapiens OX=9606 GN=GEMIN4 PE=1 SV=2 |
| Q9BZE4 | 0 | 6 | Nucleolar GTP-binding protein 1 OS=Homo sapiens OX=9606 GN=GTPBP4 PE=1 SV=3 |
| Q15233 | 0 | 6 | Non-POU domain-containing octamer-binding protein OS=Homo sapiens OX=9606 GN=NONO PE=1 SV=4 |
| Q9NPD3 | 0 | 6 | Exosome complex component RRP41 OS=Homo sapiens OX=9606 GN=EXOSC4 PE=1 SV=3 |
| O95347 | 0 | 6 | Structural maintenance of chromosomes protein 2 OS=Homo sapiens OX=9606 GN=SMC2 PE=1 SV=2 |
| Q9Y2W1 | 0 | 6 | Thyroid hormone receptor-associated protein 3 OS=Homo sapiens OX=9606 GN=THRAP3 PE=1 SV=2 |
| Q9Y676 | 0 | 6 | 28S ribosomal protein S18b mitochondrial OS=Homo sapiens OX=9606 GN=MRPS18B PE=1 SV=1 |
| P09874 | 0 | 6 | Poly [ADP-ribose] polymerase 1 OS=Homo sapiens OX=9606 GN=PARP1 PE=1 SV=4 |
| Q96MX6 | 0 | 6 | WD repeat-containing protein 92 OS=Homo sapiens OX=9606 GN=WDR92 PE=1 SV=1 |
| P61927 | 0 | 6 | 60S ribosomal protein L37 OS=Homo sapiens OX=9606 GN=RPL37 PE=1 SV=2 |
| Q5RKV6 | 0 | 5 | Exosome complex component MTR3 OS=Homo sapiens OX=9606 GN=EXOSC6 PE=1 SV=1 |
| Q7Z2T5 | 0 | 5 | TRMT1-like protein OS=Homo sapiens OX=9606 GN=TRMT1L PE=1 SV=2 |
| P11387 | 0 | 5 | DNA topoisomerase 1 OS=Homo sapiens OX=9606 GN=TOP1 PE=1 SV=2 |
| Q8N9T8 | 0 | 5 | Protein KRI1 homolog OS=Homo sapiens OX=9606 GN=KRI1 PE=1 SV=3 |
| Q9NQ29 | 0 | 5 | Putative RNA-binding protein Luc7-like 1 OS=Homo sapiens OX=9606 GN=LUC7L PE=1 SV=1 |
| Q86VM9 | 0 | 5 | Zinc finger CCCH domain-containing protein 18 OS=Homo sapiens OX=9606 GN=ZC3H18 PE=1 SV=2 |
| Q92900 | 0 | 5 | Regulator of nonsense transcripts 1 OS=Homo sapiens OX=9606 GN=UPF1 PE=1 SV=2 |
| Q9BUQ8 | 0 | 5 | Probable ATP-dependent RNA helicase DDX23 OS=Homo sapiens OX=9606 GN=DDX23 PE=1 SV=3 |
| P37108 | 0 | 5 | Signal recognition particle 14 kDa protein OS=Homo sapiens OX=9606 GN=SRP14 PE=1 SV=2 |
| P20042 | 0 | 5 | Eukaryotic translation initiation factor 2 subunit 2 OS=Homo sapiens OX=9606 GN=EIF2S2 PE=1 SV=2 |
| P25398 | 0 | 5 | 40S ribosomal protein S12 OS=Homo sapiens OX=9606 GN=RPS12 PE=1 SV=3 |
| Q9Y399 | 0 | 5 | 28S ribosomal protein S2 mitochondrial OS=Homo sapiens OX=9606 GN=MRPS2 PE=1 SV=1 |
| Q13610 | 0 | 5 | Periodic tryptophan protein 1 homolog OS=Homo sapiens OX=9606 GN=PWP1 PE=1 SV=1 |
| O60832 | 0 | 5 | H/ACA ribonucleoprotein complex subunit DKC1 OS=Homo sapiens OX=9606 GN=DKC1 PE=1 SV=3 |

| | | | |
|---|---|---|---|
| Q9NWU5 | 0 | 5 | 39S ribosomal protein L22 mitochondrial OS=Homo sapiens OX=9606 GN=MRPL22 PE=1 SV=1 |
| Q12986 | 0 | 5 | Transcriptional repressor NF-X1 OS=Homo sapiens OX=9606 GN=NFX1 PE=1 SV=2 |
| Q9BYN8 | 0 | 5 | 28S ribosomal protein S26 mitochondrial OS=Homo sapiens OX=9606 GN=MRPS26 PE=1 SV=1 |
| P14678 | 0 | 5 | Small nuclear ribonucleoprotein-associated proteins B and B' OS=Homo sapiens OX=9606 GN=SNRPB PE=1 SV=2 |
| P63162 | 0 | 5 | Small nuclear ribonucleoprotein-associated protein N OS=Homo sapiens OX=9606 GN=SNRPN PE=1 SV=1 |
| Q9NVP1 | 0 | 5 | ATP-dependent RNA helicase DDX18 OS=Homo sapiens OX=9606 GN=DDX18 PE=1 SV=2 |
| Q9Y3Y2 | 0 | 5 | Chromatin target of PRMT1 protein OS=Homo sapiens OX=9606 GN=CHTOP PE=1 SV=2 |
| Q9H8Y5 | 0 | 5 | Ankyrin repeat and zinc finger domain-containing protein 1 OS=Homo sapiens OX=9606 GN=ANKZF1 PE=1 SV=1 |
| Q13769 | 0 | 5 | THO complex subunit 5 homolog OS=Homo sapiens OX=9606 GN=THOC5 PE=1 SV=2 |
| O00505 | 0 | 5 | Importin subunit alpha-4 OS=Homo sapiens OX=9606 GN=KPNA3 PE=1 SV=2 |
| Q96I24 | 0 | 5 | Far upstream element-binding protein 3 OS=Homo sapiens OX=9606 GN=FUBP3 PE=1 SV=2 |
| P60842 | 0 | 5 | Eukaryotic initiation factor 4A-I OS=Homo sapiens OX=9606 GN=EIF4A1 PE=1 SV=1 |
| Q8IUF8 | 0 | 5 | Ribosomal oxygenase 2 OS=Homo sapiens OX=9606 GN=RIOX2 PE=1 SV=1 |
| Q9BXS6 | 0 | 5 | Nucleolar and spindle-associated protein 1 OS=Homo sapiens OX=9606 GN=NUSAP1 PE=1 SV=1 |
| Q96DV4 | 0 | 5 | 39S ribosomal protein L38 mitochondrial OS=Homo sapiens OX=9606 GN=MRPL38 PE=1 SV=2 |
| O75152 | 0 | 4 | Zinc finger CCCH domain-containing protein 11A OS=Homo sapiens OX=9606 GN=ZC3H11A PE=1 SV=3 |
| Q9H0U9 | 0 | 4 | Testis-specific Y-encoded-like protein 1 OS=Homo sapiens OX=9606 GN=TSPYL1 PE=1 SV=3 |
| Q5T3I0 | 0 | 4 | G patch domain-containing protein 4 OS=Homo sapiens OX=9606 GN=GPATCH4 PE=1 SV=2 |
| P55884 | 0 | 4 | Eukaryotic translation initiation factor 3 subunit B OS=Homo sapiens OX=9606 GN=EIF3B PE=1 SV=3 |
| Q9NUL7 | 0 | 4 | Probable ATP-dependent RNA helicase DDX28 OS=Homo sapiens OX=9606 GN=DDX28 PE=1 SV=2 |
| P51398 | 0 | 4 | 28S ribosomal protein S29 mitochondrial OS=Homo sapiens OX=9606 GN=DAP3 PE=1 SV=1 |
| Q14684 | 0 | 4 | Ribosomal RNA processing protein 1 homolog B OS=Homo sapiens OX=9606 GN=RRP1B PE=1 SV=3 |
| Q99714 | 0 | 4 | 3-hydroxyacyl-CoA dehydrogenase type-2 OS=Homo sapiens OX=9606 GN=HSD17B10 PE=1 SV=3 |
| O75533 | 0 | 4 | Splicing factor 3B subunit 1 OS=Homo sapiens OX=9606 GN=SF3B1 PE=1 SV=3 |
| Q8NHQ9 | 0 | 4 | ATP-dependent RNA helicase DDX55 OS=Homo sapiens OX=9606 GN=DDX55 PE=1 SV=3 |
| Q8N5A5 | 0 | 4 | Zinc finger CCCH-type with G patch domain-containing protein OS=Homo sapiens OX=9606 GN=ZGPAT PE=1 SV=3 |
| Q5SSJ5 | 0 | 4 | Heterochromatin protein 1-binding protein 3 OS=Homo sapiens OX=9606 GN=HP1BP3 PE=1 SV=1 |
| P62995 | 0 | 4 | Transformer-2 protein homolog beta OS=Homo sapiens OX=9606 GN=TRA2B PE=1 SV=1 |
| Q9NX58 | 0 | 4 | Cell growth-regulating nucleolar protein OS=Homo sapiens OX=9606 GN=LYAR PE=1 SV=2 |
| Q9BWF3 | 0 | 4 | RNA-binding protein 4 OS=Homo sapiens OX=9606 GN=RBM4 PE=1 SV=1 |
| P04843 | 0 | 4 | Dolichyl-diphosphooligosaccharide--protein glycosyltransferase subunit 1 OS=Homo sapiens OX=9606 GN=RPN1 PE=1 SV=1 |

| | | | |
|---|---|---|---|
| Q9NQ55 | 0 | 4 | Suppressor of SWI4 1 homolog OS=Homo sapiens OX=9606 GN=PPAN PE=2 SV=1 |
| Q14244 | 0 | 4 | Ensconsin OS=Homo sapiens OX=9606 GN=MAP7 PE=1 SV=1 |
| P04637 | 0 | 4 | Cellular tumor antigen p53 OS=Homo sapiens OX=9606 GN=TP53 PE=1 SV=4 |
| Q9BW19 | 0 | 4 | Kinesin-like protein KIFC1 OS=Homo sapiens OX=9606 GN=KIFC1 PE=1 SV=2 |
| P82673 | 0 | 4 | 28S ribosomal protein S35 mitochondrial OS=Homo sapiens OX=9606 GN=MRPS35 PE=1 SV=1 |
| O95801 | 0 | 4 | Tetratricopeptide repeat protein 4 OS=Homo sapiens OX=9606 GN=TTC4 PE=1 SV=3 |
| O00139 | 0 | 4 | Kinesin-like protein KIF2A OS=Homo sapiens OX=9606 GN=KIF2A PE=1 SV=3 |
| Q15427 | 0 | 4 | Splicing factor 3B subunit 4 OS=Homo sapiens OX=9606 GN=SF3B4 PE=1 SV=1 |
| P26368 | 0 | 4 | Splicing factor U2AF 65 kDa subunit OS=Homo sapiens OX=9606 GN=U2AF2 PE=1 SV=4 |
| Q99848 | 0 | 4 | Probable rRNA-processing protein EBP2 OS=Homo sapiens OX=9606 GN=EBNA1BP2 PE=1 SV=2 |
| Q9BZI7 | 0 | 4 | Regulator of nonsense transcripts 3B OS=Homo sapiens OX=9606 GN=UPF3B PE=1 SV=1 |
| Q9H2U1 | 0 | 4 | ATP-dependent DNA/RNA helicase DHX36 OS=Homo sapiens OX=9606 GN=DHX36 PE=1 SV=2 |
| Q53F19 | 0 | 4 | Nuclear cap-binding protein subunit 3 OS=Homo sapiens OX=9606 GN=NCBP3 PE=1 SV=2 |
| Q96C57 | 0 | 4 | Protein CUSTOS OS=Homo sapiens OX=9606 GN=CUSTOS PE=1 SV=2 |
| O76021 | 0 | 4 | Ribosomal L1 domain-containing protein 1 OS=Homo sapiens OX=9606 GN=RSL1D1 PE=1 SV=3 |
| O15381 | 0 | 4 | Nuclear valosin-containing protein-like OS=Homo sapiens OX=9606 GN=NVL PE=1 SV=1 |
| Q8WWM7 | 0 | 4 | Ataxin-2-like protein OS=Homo sapiens OX=9606 GN=ATXN2L PE=1 SV=2 |
| O00541 | 0 | 4 | Pescadillo homolog OS=Homo sapiens OX=9606 GN=PES1 PE=1 SV=1 |
| Q9BV38 | 0 | 4 | WD repeat-containing protein 18 OS=Homo sapiens OX=9606 GN=WDR18 PE=1 SV=2 |
| Q15024 | 0 | 4 | Exosome complex component RRP42 OS=Homo sapiens OX=9606 GN=EXOSC7 PE=1 SV=3 |
| P56182 | 0 | 4 | Ribosomal RNA processing protein 1 homolog A OS=Homo sapiens OX=9606 GN=RRP1 PE=1 SV=1 |
| Q9UHB9 | 0 | 4 | Signal recognition particle subunit SRP68 OS=Homo sapiens OX=9606 GN=SRP68 PE=1 SV=2 |
| Q96D09 | 0 | 4 | G-protein coupled receptor-associated sorting protein 2 OS=Homo sapiens OX=9606 GN=GPRASP2 PE=1 SV=1 |
| O94763 | 0 | 4 | Unconventional prefoldin RPB5 interactor 1 OS=Homo sapiens OX=9606 GN=URI1 PE=1 SV=3 |
| Q9UEG4 | 0 | 4 | Zinc finger protein 629 OS=Homo sapiens OX=9606 GN=ZNF629 PE=1 SV=2 |
| Q13428 | 0 | 4 | Treacle protein OS=Homo sapiens OX=9606 GN=TCOF1 PE=1 SV=3 |
| Q9Y450 | 0 | 4 | HBS1-like protein OS=Homo sapiens OX=9606 GN=HBS1L PE=1 SV=1 |
| O95625 | 0 | 4 | Zinc finger and BTB domain-containing protein 11 OS=Homo sapiens OX=9606 GN=ZBTB11 PE=1 SV=2 |
| O75683 | 0 | 4 | Surfeit locus protein 6 OS=Homo sapiens OX=9606 GN=SURF6 PE=1 SV=3 |
| P12532 | 0 | 4 | Creatine kinase U-type mitochondrial OS=Homo sapiens OX=9606 GN=CKMT1A PE=1 SV=1 |
| Q12797 | 0 | 4 | Aspartyl/asparaginyl beta-hydroxylase OS=Homo sapiens OX=9606 GN=ASPH PE=1 SV=3 |

| | | | |
|---|---|---|---|
| P19388 | 0 | 4 | DNA-directed RNA polymerases I II and III subunit RPABC1 OS=Homo sapiens OX=9606 GN=POLR2E PE=1 SV=4 |
| Q00577 | 0 | 4 | Transcriptional activator protein Pur-alpha OS=Homo sapiens OX=9606 GN=PURA PE=1 SV=2 |
| Q9BUJ2 | 0 | 4 | Heterogeneous nuclear ribonucleoprotein U-like protein 1 OS=Homo sapiens OX=9606 GN=HNRNPUL1 PE=1 SV=2 |
| O94906 | 0 | 4 | Pre-mRNA-processing factor 6 OS=Homo sapiens OX=9606 GN=PRPF6 PE=1 SV=1 |
| Q9BX40 | 0 | 4 | Protein LSM14 homolog B OS=Homo sapiens OX=9606 GN=LSM14B PE=1 SV=1 |
| P35250 | 0 | 4 | Replication factor C subunit 2 OS=Homo sapiens OX=9606 GN=RFC2 PE=1 SV=3 |
| Q96B26 | 0 | 3 | Exosome complex component RRP43 OS=Homo sapiens OX=9606 GN=EXOSC8 PE=1 SV=1 |
| Q9H0W5 | 0 | 3 | Coiled-coil domain-containing protein 8 OS=Homo sapiens OX=9606 GN=CCDC8 PE=1 SV=2 |
| Q9BY77 | 0 | 3 | Polymerase delta-interacting protein 3 OS=Homo sapiens OX=9606 GN=POLDIP3 PE=1 SV=2 |
| Q8N5F7 | 0 | 3 | NF-kappa-B-activating protein OS=Homo sapiens OX=9606 GN=NKAP PE=1 SV=1 |
| Q5C9Z4 | 0 | 3 | Nucleolar MIF4G domain-containing protein 1 OS=Homo sapiens OX=9606 GN=NOM1 PE=1 SV=1 |
| Q99613 | 0 | 3 | Eukaryotic translation initiation factor 3 subunit C OS=Homo sapiens OX=9606 GN=EIF3C PE=1 SV=1 |
| B5ME19 | 0 | 3 | Eukaryotic translation initiation factor 3 subunit C-like protein OS=Homo sapiens OX=9606 GN=EIF3CL PE=3 SV=1 |
| P84090 | 0 | 3 | Enhancer of rudimentary homolog OS=Homo sapiens OX=9606 GN=ERH PE=1 SV=1 |
| Q96T37 | 0 | 3 | RNA-binding protein 15 OS=Homo sapiens OX=9606 GN=RBM15 PE=1 SV=2 |
| Q9H0S4 | 0 | 3 | Probable ATP-dependent RNA helicase DDX47 OS=Homo sapiens OX=9606 GN=DDX47 PE=1 SV=1 |
| O00410 | 0 | 3 | Importin-5 OS=Homo sapiens OX=9606 GN=IPO5 PE=1 SV=4 |
| P78316 | 0 | 3 | Nucleolar protein 14 OS=Homo sapiens OX=9606 GN=NOP14 PE=1 SV=3 |
| P35637 | 0 | 3 | RNA-binding protein FUS OS=Homo sapiens OX=9606 GN=FUS PE=1 SV=1 |
| O43290 | 0 | 3 | U4/U6.U5 tri-snRNP-associated protein 1 OS=Homo sapiens OX=9606 GN=SART1 PE=1 SV=1 |
| Q9H0A0 | 0 | 3 | RNA cytidine acetyltransferase OS=Homo sapiens OX=9606 GN=NAT10 PE=1 SV=2 |
| P08579 | 0 | 3 | U2 small nuclear ribonucleoprotein B'' OS=Homo sapiens OX=9606 GN=SNRPB2 PE=1 SV=1 |
| Q96T21 | 0 | 3 | Selenocysteine insertion sequence-binding protein 2 OS=Homo sapiens OX=9606 GN=SECISBP2 PE=1 SV=2 |
| P60660 | 0 | 3 | Myosin light polypeptide 6 OS=Homo sapiens OX=9606 GN=MYL6 PE=1 SV=2 |
| Q13573 | 0 | 3 | SNW domain-containing protein 1 OS=Homo sapiens OX=9606 GN=SNW1 PE=1 SV=1 |
| O00178 | 0 | 3 | GTP-binding protein 1 OS=Homo sapiens OX=9606 GN=GTPBP1 PE=1 SV=3 |
| Q96HC4 | 0 | 3 | PDZ and LIM domain protein 5 OS=Homo sapiens OX=9606 GN=PDLIM5 PE=1 SV=5 |
| Q09161 | 0 | 3 | Nuclear cap-binding protein subunit 1 OS=Homo sapiens OX=9606 GN=NCBP1 PE=1 SV=1 |
| Q96QR8 | 0 | 3 | Transcriptional activator protein Pur-beta OS=Homo sapiens OX=9606 GN=PURB PE=1 SV=3 |
| O00159 | 0 | 3 | Unconventional myosin-Ic OS=Homo sapiens OX=9606 GN=MYO1C PE=1 SV=4 |
| Q8IX01 | 0 | 3 | SURP and G-patch domain-containing protein 2 OS=Homo sapiens OX=9606 GN=SUGP2 PE=1 SV=2 |

| | | | |
|---|---|---|---|
| O43395 | 0 | 3 | U4/U6 small nuclear ribonucleoprotein Prp3 OS=Homo sapiens OX=9606 GN=PRPF3 PE=1 SV=2 |
| Q9UN81 | 0 | 3 | LINE-1 retrotransposable element ORF1 protein OS=Homo sapiens OX=9606 GN=L1RE1 PE=1 SV=1 |
| O75821 | 0 | 3 | Eukaryotic translation initiation factor 3 subunit G OS=Homo sapiens OX=9606 GN=EIF3G PE=1 SV=2 |
| Q8IXB1 | 0 | 3 | DnaJ homolog subfamily C member 10 OS=Homo sapiens OX=9606 GN=DNAJC10 PE=1 SV=2 |
| O00566 | 0 | 3 | U3 small nucleolar ribonucleoprotein protein MPP10 OS=Homo sapiens OX=9606 GN=MPHOSPH10 PE=1 SV=2 |
| Q9UNM6 | 0 | 3 | 26S proteasome non-ATPase regulatory subunit 13 OS=Homo sapiens OX=9606 GN=PSMD13 PE=1 SV=2 |
| Q8TDD1 | 0 | 3 | ATP-dependent RNA helicase DDX54 OS=Homo sapiens OX=9606 GN=DDX54 PE=1 SV=2 |
| Q14257 | 0 | 3 | Reticulocalbin-2 OS=Homo sapiens OX=9606 GN=RCN2 PE=1 SV=1 |
| Q9Y3X0 | 0 | 3 | Coiled-coil domain-containing protein 9 OS=Homo sapiens OX=9606 GN=CCDC9 PE=1 SV=1 |
| P25205 | 0 | 3 | DNA replication licensing factor MCM3 OS=Homo sapiens OX=9606 GN=MCM3 PE=1 SV=3 |
| P63208 | 0 | 3 | S-phase kinase-associated protein 1 OS=Homo sapiens OX=9606 GN=SKP1 PE=1 SV=2 |
| Q9NSD9 | 0 | 3 | Phenylalanine--tRNA ligase beta subunit OS=Homo sapiens OX=9606 GN=FARSB PE=1 SV=3 |
| Q9UKV3 | 0 | 3 | Apoptotic chromatin condensation inducer in the nucleus OS=Homo sapiens OX=9606 GN=ACIN1 PE=1 SV=2 |
| P35659 | 0 | 3 | Protein DEK OS=Homo sapiens OX=9606 GN=DEK PE=1 SV=1 |
| Q9BQ67 | 0 | 3 | Glutamate-rich WD repeat-containing protein 1 OS=Homo sapiens OX=9606 GN=GRWD1 PE=1 SV=1 |
| P21127 | 0 | 3 | Cyclin-dependent kinase 11B OS=Homo sapiens OX=9606 GN=CDK11B PE=1 SV=4 |
| Q9UQ88 | 0 | 3 | Cyclin-dependent kinase 11A OS=Homo sapiens OX=9606 GN=CDK11A PE=1 SV=4 |
| Q9H0D6 | 0 | 3 | 5'-3' exoribonuclease 2 OS=Homo sapiens OX=9606 GN=XRN2 PE=1 SV=1 |
| Q9NXG2 | 0 | 3 | THUMP domain-containing protein 1 OS=Homo sapiens OX=9606 GN=THUMPD1 PE=1 SV=2 |
| Q9UH62 | 0 | 3 | Armadillo repeat-containing X-linked protein 3 OS=Homo sapiens OX=9606 GN=ARMCX3 PE=1 SV=1 |
| Q9NVN8 | 0 | 3 | Guanine nucleotide-binding protein-like 3-like protein OS=Homo sapiens OX=9606 GN=GNL3L PE=1 SV=1 |
| Q92667 | 0 | 3 | A-kinase anchor protein 1 mitochondrial OS=Homo sapiens OX=9606 GN=AKAP1 PE=1 SV=1 |
| Q96DI7 | 0 | 3 | U5 small nuclear ribonucleoprotein 40 kDa protein OS=Homo sapiens OX=9606 GN=SNRNP40 PE=1 SV=1 |
| Q9UHK0 | 0 | 3 | Nuclear fragile X mental retardation-interacting protein 1 OS=Homo sapiens OX=9606 GN=NUFIP1 PE=1 SV=2 |
| Q96GA3 | 0 | 3 | Protein LTV1 homolog OS=Homo sapiens OX=9606 GN=LTV1 PE=1 SV=1 |
| Q14974 | 0 | 3 | Importin subunit beta-1 OS=Homo sapiens OX=9606 GN=KPNB1 PE=1 SV=2 |
| O43852 | 0 | 3 | Calumenin OS=Homo sapiens OX=9606 GN=CALU PE=1 SV=2 |
| O60524 | 0 | 3 | Nuclear export mediator factor NEMF OS=Homo sapiens OX=9606 GN=NEMF PE=1 SV=4 |
| P62857 | 0 | 3 | 40S ribosomal protein S28 OS=Homo sapiens OX=9606 GN=RPS28 PE=1 SV=1 |
| P12956 | 0 | 3 | X-ray repair cross-complementing protein 6 OS=Homo sapiens OX=9606 GN=XRCC6 PE=1 SV=2 |
| Q15287 | 0 | 3 | RNA-binding protein with serine-rich domain 1 OS=Homo sapiens OX=9606 GN=RNPS1 PE=1 SV=1 |

| | | | |
|---|---|---|---|
| Q04837 | 0 | 3 | Single-stranded DNA-binding protein mitochondrial OS=Homo sapiens OX=9606 GN=SSBP1 PE=1 SV=1 |
| Q9H501 | 0 | 3 | ESF1 homolog OS=Homo sapiens OX=9606 GN=ESF1 PE=1 SV=1 |
| Q9H307 | 0 | 3 | Pinin OS=Homo sapiens OX=9606 GN=PNN PE=1 SV=5 |
| P19525 | 0 | 3 | Interferon-induced double-stranded RNA-activated protein kinase OS=Homo sapiens OX=9606 GN=EIF2AK2 PE=1 SV=2 |
| Q96S55 | 0 | 3 | ATPase WRNIP1 OS=Homo sapiens OX=9606 GN=WRNIP1 PE=1 SV=2 |
| Q9UII4 | 0 | 3 | E3 ISG15--protein ligase HERC5 OS=Homo sapiens OX=9606 GN=HERC5 PE=1 SV=2 |
| Q9UHI6 | 0 | 3 | Probable ATP-dependent RNA helicase DDX20 OS=Homo sapiens OX=9606 GN=DDX20 PE=1 SV=2 |
| Q16531 | 0 | 3 | DNA damage-binding protein 1 OS=Homo sapiens OX=9606 GN=DDB1 PE=1 SV=1 |
| O94832 | 0 | 3 | Unconventional myosin-Id OS=Homo sapiens OX=9606 GN=MYO1D PE=1 SV=2 |
| P40939 | 0 | 3 | Trifunctional enzyme subunit alpha mitochondrial OS=Homo sapiens OX=9606 GN=HADHA PE=1 SV=2 |
| Q96P70 | 0 | 3 | Importin-9 OS=Homo sapiens OX=9606 GN=IPO9 PE=1 SV=3 |
| P82675 | 0 | 3 | 28S ribosomal protein S5 mitochondrial OS=Homo sapiens OX=9606 GN=MRPS5 PE=1 SV=2 |
| Q8IV48 | 0 | 3 | 3'-5' exoribonuclease 1 OS=Homo sapiens OX=9606 GN=ERI1 PE=1 SV=3 |
| P62316 | 0 | 3 | Small nuclear ribonucleoprotein Sm D2 OS=Homo sapiens OX=9606 GN=SNRPD2 PE=1 SV=1 |
| Q9Y2Q9 | 0 | 3 | 28S ribosomal protein S28 mitochondrial OS=Homo sapiens OX=9606 GN=MRPS28 PE=1 SV=1 |
| Q13868 | 0 | 3 | Exosome complex component RRP4 OS=Homo sapiens OX=9606 GN=EXOSC2 PE=1 SV=2 |
| Q5F1R6 | 0 | 2 | DnaJ homolog subfamily C member 21 OS=Homo sapiens OX=9606 GN=DNAJC21 PE=1 SV=2 |
| Q86XN8 | 0 | 2 | RNA-binding protein MEX3D OS=Homo sapiens OX=9606 GN=MEX3D PE=1 SV=3 |
| Q9Y295 | 0 | 2 | Developmentally-regulated GTP-binding protein 1 OS=Homo sapiens OX=9606 GN=DRG1 PE=1 SV=1 |
| Q9H4L4 | 0 | 2 | Sentrin-specific protease 3 OS=Homo sapiens OX=9606 GN=SENP3 PE=1 SV=2 |
| Q5JTW2 | 0 | 2 | Centrosomal protein of 78 kDa OS=Homo sapiens OX=9606 GN=CEP78 PE=1 SV=1 |
| Q13405 | 0 | 2 | 39S ribosomal protein L49 mitochondrial OS=Homo sapiens OX=9606 GN=MRPL49 PE=1 SV=1 |
| Q16643 | 0 | 2 | Drebrin OS=Homo sapiens OX=9606 GN=DBN1 PE=1 SV=4 |
| Q8IVS2 | 0 | 2 | Malonyl-CoA-acyl carrier protein transacylase mitochondrial OS=Homo sapiens OX=9606 GN=MCAT PE=1 SV=2 |
| P61289 | 0 | 2 | Proteasome activator complex subunit 3 OS=Homo sapiens OX=9606 GN=PSME3 PE=1 SV=1 |
| Q15393 | 0 | 2 | Splicing factor 3B subunit 3 OS=Homo sapiens OX=9606 GN=SF3B3 PE=1 SV=4 |
| P00750 | 0 | 2 | Tissue-type plasminogen activator OS=Homo sapiens OX=9606 GN=PLAT PE=1 SV=1 |
| Q9NUD5 | 0 | 2 | Zinc finger CCHC domain-containing protein 3 OS=Homo sapiens OX=9606 GN=ZCCHC3 PE=1 SV=2 |
| Q9NY93 | 0 | 2 | Probable ATP-dependent RNA helicase DDX56 OS=Homo sapiens OX=9606 GN=DDX56 PE=1 SV=1 |
| Q9Y2R9 | 0 | 2 | 28S ribosomal protein S7 mitochondrial OS=Homo sapiens OX=9606 GN=MRPS7 PE=1 SV=2 |
| Q1ED39 | 0 | 2 | Lysine-rich nucleolar protein 1 OS=Homo sapiens OX=9606 GN=KNOP1 PE=1 SV=1 |

| | | | |
|---|---|---|---|
| Q659C4 | 0 | 2 | La-related protein 1B OS=Homo sapiens OX=9606 GN=LARP1B PE=1 SV=2 |
| Q92804 | 0 | 2 | TATA-binding protein-associated factor 2N OS=Homo sapiens OX=9606 GN=TAF15 PE=1 SV=1 |
| Q86XZ4 | 0 | 2 | Spermatogenesis-associated serine-rich protein 2 OS=Homo sapiens OX=9606 GN=SPATS2 PE=1 SV=1 |
| Q96EY1 | 0 | 2 | DnaJ homolog subfamily A member 3 mitochondrial OS=Homo sapiens OX=9606 GN=DNAJA3 PE=1 SV=2 |
| P08754 | 0 | 2 | Guanine nucleotide-binding protein G(i) subunit alpha OS=Homo sapiens OX=9606 GN=GNAI3 PE=1 SV=3 |
| Q13123 | 0 | 2 | Protein Red OS=Homo sapiens OX=9606 GN=IK PE=1 SV=3 |
| P13489 | 0 | 2 | Ribonuclease inhibitor OS=Homo sapiens OX=9606 GN=RNH1 PE=1 SV=2 |
| O00629 | 0 | 2 | Importin subunit alpha-3 OS=Homo sapiens OX=9606 GN=KPNA4 PE=1 SV=1 |
| P07305 | 0 | 2 | Histone H1.0 OS=Homo sapiens OX=9606 GN=H1F0 PE=1 SV=3 |
| Q9Y3B2 | 0 | 2 | Exosome complex component CSL4 OS=Homo sapiens OX=9606 GN=EXOSC1 PE=1 SV=1 |
| O95391 | 0 | 2 | Pre-mRNA-splicing factor SLU7 OS=Homo sapiens OX=9606 GN=SLU7 PE=1 SV=2 |
| O43159 | 0 | 2 | Ribosomal RNA-processing protein 8 OS=Homo sapiens OX=9606 GN=RRP8 PE=1 SV=2 |
| Q5VWQ0 | 0 | 2 | Lysine-specific demethylase 9 OS=Homo sapiens OX=9606 GN=RSBN1 PE=1 SV=2 |
| Q9P2J5 | 0 | 2 | Leucine--tRNA ligase cytoplasmic OS=Homo sapiens OX=9606 GN=LARS PE=1 SV=2 |
| Q99623 | 0 | 2 | Prohibitin-2 OS=Homo sapiens OX=9606 GN=PHB2 PE=1 SV=2 |
| P07237 | 0 | 2 | Protein disulfide-isomerase OS=Homo sapiens OX=9606 GN=P4HB PE=1 SV=3 |
| P25788 | 0 | 2 | Proteasome subunit alpha type-3 OS=Homo sapiens OX=9606 GN=PSMA3 PE=1 SV=2 |
| Q8WXI9 | 0 | 2 | Transcriptional repressor p66-beta OS=Homo sapiens OX=9606 GN=GATAD2B PE=1 SV=1 |
| O60264 | 0 | 2 | SWI/SNF-related matrix-associated actin-dependent regulator of chromatin subfamily A member 5 OS=Homo sapiens OX=9606 GN=SMARCA5 PE=1 SV=1 |
| O15479 | 0 | 2 | Melanoma-associated antigen B2 OS=Homo sapiens OX=9606 GN=MAGEB2 PE=1 SV=3 |
| O43663 | 0 | 2 | Protein regulator of cytokinesis 1 OS=Homo sapiens OX=9606 GN=PRC1 PE=1 SV=2 |
| O95295 | 0 | 2 | SNARE-associated protein Snapin OS=Homo sapiens OX=9606 GN=SNAPIN PE=1 SV=1 |
| Q14592 | 0 | 2 | Zinc finger protein 460 OS=Homo sapiens OX=9606 GN=ZNF460 PE=1 SV=2 |
| Q8IY37 | 0 | 2 | Probable ATP-dependent RNA helicase DHX37 OS=Homo sapiens OX=9606 GN=DHX37 PE=1 SV=1 |
| P62879 | 0 | 2 | Guanine nucleotide-binding protein G(I)/G(S)/G(T) subunit beta-2 OS=Homo sapiens OX=9606 GN=GNB2 PE=1 SV=3 |
| P62873 | 0 | 2 | Guanine nucleotide-binding protein G(I)/G(S)/G(T) subunit beta-1 OS=Homo sapiens OX=9606 GN=GNB1 PE=1 SV=3 |
| Q9HAV0 | 0 | 2 | Guanine nucleotide-binding protein subunit beta-4 OS=Homo sapiens OX=9606 GN=GNB4 PE=1 SV=3 |
| P36873 | 0 | 2 | Serine/threonine-protein phosphatase PP1-gamma catalytic subunit OS=Homo sapiens OX=9606 GN=PPP1CC PE=1 SV=1 |
| O43670 | 0 | 2 | BUB3-interacting and GLEBS motif-containing protein ZNF207 OS=Homo sapiens OX=9606 GN=ZNF207 PE=1 SV=1 |
| Q15637 | 0 | 2 | Splicing factor 1 OS=Homo sapiens OX=9606 GN=SF1 PE=1 SV=4 |

| | | | |
|---|---|---|---|
| O15371 | 0 | 2 | Eukaryotic translation initiation factor 3 subunit D OS=Homo sapiens OX=9606 GN=EIF3D PE=1 SV=1 |
| Q01844 | 0 | 2 | RNA-binding protein EWS OS=Homo sapiens OX=9606 GN=EWSR1 PE=1 SV=1 |
| Q14137 | 0 | 2 | Ribosome biogenesis protein BOP1 OS=Homo sapiens OX=9606 GN=BOP1 PE=1 SV=2 |
| A1L020 | 0 | 2 | RNA-binding protein MEX3A OS=Homo sapiens OX=9606 GN=MEX3A PE=1 SV=1 |
| Q9UL40 | 0 | 2 | Zinc finger protein 346 OS=Homo sapiens OX=9606 GN=ZNF346 PE=1 SV=1 |
| P63092 | 0 | 2 | Guanine nucleotide-binding protein G(s) subunit alpha isoforms short OS=Homo sapiens OX=9606 GN=GNAS PE=1 SV=1 |
| Q5JWF2 | 0 | 2 | Guanine nucleotide-binding protein G(s) subunit alpha isoforms XLas OS=Homo sapiens OX=9606 GN=GNAS PE=1 SV=2 |
| P82930 | 0 | 2 | 28S ribosomal protein S34 mitochondrial OS=Homo sapiens OX=9606 GN=MRPS34 PE=1 SV=2 |
| Q9UK59 | 0 | 2 | Lariat debranching enzyme OS=Homo sapiens OX=9606 GN=DBR1 PE=1 SV=2 |
| Q7KZI7 | 0 | 2 | Serine/threonine-protein kinase MARK2 OS=Homo sapiens OX=9606 GN=MARK2 PE=1 SV=2 |
| Q9H9J2 | 0 | 2 | 39S ribosomal protein L44 mitochondrial OS=Homo sapiens OX=9606 GN=MRPL44 PE=1 SV=1 |
| Q9Y5B9 | 0 | 2 | FACT complex subunit SPT16 OS=Homo sapiens OX=9606 GN=SUPT16H PE=1 SV=1 |
| Q9Y2P8 | 0 | 2 | RNA 3'-terminal phosphate cyclase-like protein OS=Homo sapiens OX=9606 GN=RCL1 PE=1 SV=3 |
| Q8WUA2 | 0 | 2 | Peptidyl-prolyl cis-trans isomerase-like 4 OS=Homo sapiens OX=9606 GN=PPIL4 PE=1 SV=1 |
| O43172 | 0 | 2 | U4/U6 small nuclear ribonucleoprotein Prp4 OS=Homo sapiens OX=9606 GN=PRPF4 PE=1 SV=2 |
| Q9H7B2 | 0 | 2 | Ribosome production factor 2 homolog OS=Homo sapiens OX=9606 GN=RPF2 PE=1 SV=2 |
| P13987 | 0 | 2 | CD59 glycoprotein OS=Homo sapiens OX=9606 GN=CD59 PE=1 SV=1 |
| O15235 | 0 | 2 | 28S ribosomal protein S12 mitochondrial OS=Homo sapiens OX=9606 GN=MRPS12 PE=1 SV=1 |
| P43686 | 0 | 2 | 26S proteasome regulatory subunit 6B OS=Homo sapiens OX=9606 GN=PSMC4 PE=1 SV=2 |
| Q8TF39 | 0 | 2 | Zinc finger protein 483 OS=Homo sapiens OX=9606 GN=ZNF483 PE=1 SV=3 |
| Q9Y262 | 0 | 2 | Eukaryotic translation initiation factor 3 subunit L OS=Homo sapiens OX=9606 GN=EIF3L PE=1 SV=1 |
| Q14978 | 0 | 2 | Nucleolar and coiled-body phosphoprotein 1 OS=Homo sapiens OX=9606 GN=NOLC1 PE=1 SV=2 |
| P35251 | 0 | 2 | Replication factor C subunit 1 OS=Homo sapiens OX=9606 GN=RFC1 PE=1 SV=4 |
| Q9BRZ2 | 0 | 2 | E3 ubiquitin-protein ligase TRIM56 OS=Homo sapiens OX=9606 GN=TRIM56 PE=1 SV=3 |
| O75934 | 0 | 2 | Pre-mRNA-splicing factor SPF27 OS=Homo sapiens OX=9606 GN=BCAS2 PE=1 SV=1 |
| Q13595 | 0 | 2 | Transformer-2 protein homolog alpha OS=Homo sapiens OX=9606 GN=TRA2A PE=1 SV=1 |
| Q15185 | 0 | 2 | Prostaglandin E synthase 3 OS=Homo sapiens OX=9606 GN=PTGES3 PE=1 SV=1 |
| P28072 | 0 | 2 | Proteasome subunit beta type-6 OS=Homo sapiens OX=9606 GN=PSMB6 PE=1 SV=4 |
| Q9HCM4 | 0 | 2 | Band 4.1-like protein 5 OS=Homo sapiens OX=9606 GN=EPB41L5 PE=1 SV=3 |
| Q13601 | 0 | 2 | KRR1 small subunit processome component homolog OS=Homo sapiens OX=9606 GN=KRR1 PE=1 SV=4 |
| Q9Y4X5 | 0 | 2 | E3 ubiquitin-protein ligase ARIH1 OS=Homo sapiens OX=9606 GN=ARIH1 PE=1 SV=2 |

| | | | |
|---|---|---|---|
| Q9Y3B7 | 0 | 2 | 39S ribosomal protein L11 mitochondrial OS=Homo sapiens OX=9606 GN=MRPL11 PE=1 SV=1 |
| P40937 | 0 | 2 | Replication factor C subunit 5 OS=Homo sapiens OX=9606 GN=RFC5 PE=1 SV=1 |
| Q15008 | 0 | 2 | 26S proteasome non-ATPase regulatory subunit 6 OS=Homo sapiens OX=9606 GN=PSMD6 PE=1 SV=1 |
| O00567 | 0 | 2 | Nucleolar protein 56 OS=Homo sapiens OX=9606 GN=NOP56 PE=1 SV=4 |
| Q15776 | 0 | 2 | Zinc finger protein with KRAB and SCAN domains 8 OS=Homo sapiens OX=9606 GN=ZKSCAN8 PE=1 SV=2 |
| P53041 | 0 | 2 | Serine/threonine-protein phosphatase 5 OS=Homo sapiens OX=9606 GN=PPP5C PE=1 SV=1 |
| Q8N983 | 0 | 2 | 39S ribosomal protein L43 mitochondrial OS=Homo sapiens OX=9606 GN=MRPL43 PE=1 SV=1 |
| Q9NSI2 | 0 | 2 | Protein FAM207A OS=Homo sapiens OX=9606 GN=FAM207A PE=1 SV=2 |
| O00411 | 0 | 2 | DNA-directed RNA polymerase mitochondrial OS=Homo sapiens OX=9606 GN=POLRMT PE=1 SV=2 |
| Q9Y5Q8 | 0 | 2 | General transcription factor 3C polypeptide 5 OS=Homo sapiens OX=9606 GN=GTF3C5 PE=1 SV=2 |
| O00257 | 0 | 2 | E3 SUMO-protein ligase CBX4 OS=Homo sapiens OX=9606 GN=CBX4 PE=1 SV=3 |
| Q9Y388 | 0 | 2 | RNA-binding motif protein X-linked 2 OS=Homo sapiens OX=9606 GN=RBMX2 PE=1 SV=2 |
| Q92979 | 0 | 2 | Ribosomal RNA small subunit methyltransferase NEP1 OS=Homo sapiens OX=9606 GN=EMG1 PE=1 SV=4 |
| P61326 | 0 | 2 | Protein mago nashi homolog OS=Homo sapiens OX=9606 GN=MAGOH PE=1 SV=1 |
| Q96A72 | 0 | 2 | Protein mago nashi homolog 2 OS=Homo sapiens OX=9606 GN=MAGOHB PE=1 SV=1 |
| Q9H814 | 0 | 2 | Phosphorylated adapter RNA export protein OS=Homo sapiens OX=9606 GN=PHAX PE=1 SV=1 |
| O43818 | 0 | 2 | U3 small nucleolar RNA-interacting protein 2 OS=Homo sapiens OX=9606 GN=RRP9 PE=1 SV=1 |
| Q9H992 | 0 | 2 | E3 ubiquitin-protein ligase MARCH7 OS=Homo sapiens OX=9606 GN=MARCH7 PE=1 SV=1 |
| O75127 | 0 | 2 | Pentatricopeptide repeat-containing protein 1 mitochondrial OS=Homo sapiens OX=9606 GN=PTCD1 PE=1 SV=2 |
| Q9BX10 | 0 | 2 | GTP-binding protein 2 OS=Homo sapiens OX=9606 GN=GTPBP2 PE=1 SV=1 |
| O95400 | 0 | 2 | CD2 antigen cytoplasmic tail-binding protein 2 OS=Homo sapiens OX=9606 GN=CD2BP2 PE=1 SV=1 |
| Q13724 | 0 | 2 | Mannosyl-oligosaccharide glucosidase OS=Homo sapiens OX=9606 GN=MOGS PE=1 SV=5 |
| Q9NWT1 | 0 | 2 | p21-activated protein kinase-interacting protein 1 OS=Homo sapiens OX=9606 GN=PAK1IP1 PE=1 SV=2 |
| P53999 | 0 | 2 | Activated RNA polymerase II transcriptional coactivator p15 OS=Homo sapiens OX=9606 GN=SUB1 PE=1 SV=3 |
| P49590 | 0 | 2 | Probable histidine--tRNA ligase mitochondrial OS=Homo sapiens OX=9606 GN=HARS2 PE=1 SV=1 |
| Q15369 | 0 | 2 | Elongin-C OS=Homo sapiens OX=9606 GN=ELOC PE=1 SV=1 |
| Q02040 | 0 | 2 | A-kinase anchor protein 17A OS=Homo sapiens OX=9606 GN=AKAP17A PE=1 SV=2 |
| Q14004 | 0 | 2 | Cyclin-dependent kinase 13 OS=Homo sapiens OX=9606 GN=CDK13 PE=1 SV=2 |
| Q08945 | 0 | 2 | FACT complex subunit SSRP1 OS=Homo sapiens OX=9606 GN=SSRP1 PE=1 SV=1 |
| Q6PK04 | 0 | 2 | Coiled-coil domain-containing protein 137 OS=Homo sapiens OX=9606 GN=CCDC137 PE=1 SV=1 |
| Q92974 | 0 | 2 | Rho guanine nucleotide exchange factor 2 OS=Homo sapiens OX=9606 GN=ARHGEF2 PE=1 SV=4 |

| | | | |
|---|---|---|---|
| Q16540 | 0 | 2 | 39S ribosomal protein L23 mitochondrial OS=Homo sapiens OX=9606 GN=MRPL23 PE=1 SV=1 |
| Q9P2I0 | 0 | 2 | Cleavage and polyadenylation specificity factor subunit 2 OS=Homo sapiens OX=9606 GN=CPSF2 PE=1 SV=2 |
| Q8TCC3 | 0 | 2 | 39S ribosomal protein L30 mitochondrial OS=Homo sapiens OX=9606 GN=MRPL30 PE=1 SV=1 |
| P19784 | 0 | 2 | Casein kinase II subunit alpha' OS=Homo sapiens OX=9606 GN=CSNK2A2 PE=1 SV=1 |
| Q99460 | 0 | 2 | 26S proteasome non-ATPase regulatory subunit 1 OS=Homo sapiens OX=9606 GN=PSMD1 PE=1 SV=2 |
| P63241 | 0 | 2 | Eukaryotic translation initiation factor 5A-1 OS=Homo sapiens OX=9606 GN=EIF5A PE=1 SV=2 |
| Q6IS14 | 0 | 2 | Eukaryotic translation initiation factor 5A-1-like OS=Homo sapiens OX=9606 GN=EIF5AL1 PE=2 SV=2 |
| Q9GZV4 | 0 | 2 | Eukaryotic translation initiation factor 5A-2 OS=Homo sapiens OX=9606 GN=EIF5A2 PE=1 SV=3 |
| Q9UJZ1 | 0 | 2 | Stomatin-like protein 2 mitochondrial OS=Homo sapiens OX=9606 GN=STOML2 PE=1 SV=1 |
| Q13643 | 0 | 2 | Four and a half LIM domains protein 3 OS=Homo sapiens OX=9606 GN=FHL3 PE=1 SV=4 |
| Q9P2J9 | 0 | 2 | [Pyruvate dehydrogenase [acetyl-transferring]]-phosphatase 2 mitochondrial OS=Homo sapiens OX=9606 GN=PDP2 PE=2 SV=2 |
| Q6T4R5 | 0 | 2 | Nance-Horan syndrome protein OS=Homo sapiens OX=9606 GN=NHS PE=1 SV=2 |
| P09651 | 1 | 97 | Heterogeneous nuclear ribonucleoprotein A1 OS=Homo sapiens OX=9606 GN=HNRNPA1 PE=1 SV=5 |
| P22626 | 2 | 188 | Heterogeneous nuclear ribonucleoproteins A2/B1 OS=Homo sapiens OX=9606 GN=HNRNPA2B1 PE=1 SV=2 |
| P46777 | 1 | 58 | 60S ribosomal protein L5 OS=Homo sapiens OX=9606 GN=RPL5 PE=1 SV=3 |
| P19338 | 3 | 160 | Nucleolin OS=Homo sapiens OX=9606 GN=NCL PE=1 SV=3 |
| P07910 | 1 | 49 | Heterogeneous nuclear ribonucleoproteins C1/C2 OS=Homo sapiens OX=9606 GN=HNRNPC PE=1 SV=4 |
| P50914 | 1 | 42 | 60S ribosomal protein L14 OS=Homo sapiens OX=9606 GN=RPL14 PE=1 SV=4 |
| P16403 | 2 | 70 | Histone H1.2 OS=Homo sapiens OX=9606 GN=HIST1H1C PE=1 SV=2 |
| P61254 | 1 | 32 | 60S ribosomal protein L26 OS=Homo sapiens OX=9606 GN=RPL26 PE=1 SV=1 |
| O14979 | 1 | 32 | Heterogeneous nuclear ribonucleoprotein D-like OS=Homo sapiens OX=9606 GN=HNRNPDL PE=1 SV=3 |
| P31943 | 3 | 89 | Heterogeneous nuclear ribonucleoprotein H OS=Homo sapiens OX=9606 GN=HNRNPH1 PE=1 SV=4 |
| P16989 | 2 | 57 | Y-box-binding protein 3 OS=Homo sapiens OX=9606 GN=YBX3 PE=1 SV=4 |
| Q9UNX3 | 1 | 27 | 60S ribosomal protein L26-like 1 OS=Homo sapiens OX=9606 GN=RPL26L1 PE=1 SV=1 |
| P46781 | 1 | 27 | 40S ribosomal protein S9 OS=Homo sapiens OX=9606 GN=RPS9 PE=1 SV=3 |
| Q9NR30 | 1 | 23 | Nucleolar RNA helicase 2 OS=Homo sapiens OX=9606 GN=DDX21 PE=1 SV=5 |
| O75534 | 1 | 23 | Cold shock domain-containing protein E1 OS=Homo sapiens OX=9606 GN=CSDE1 PE=1 SV=2 |
| P83881 | 1 | 23 | 60S ribosomal protein L36a OS=Homo sapiens OX=9606 GN=RPL36A PE=1 SV=2 |
| P18621 | 3 | 68 | 60S ribosomal protein L17 OS=Homo sapiens OX=9606 GN=RPL17 PE=1 SV=3 |
| P62917 | 3 | 66 | 60S ribosomal protein L8 OS=Homo sapiens OX=9606 GN=RPL8 PE=1 SV=2 |

| | | | |
|---|---|---|---|
| P62249 | 2 | 42 | 40S ribosomal protein S16 OS=Homo sapiens OX=9606 GN=RPS16 PE=1 SV=2 |
| P83731 | 1 | 21 | 60S ribosomal protein L24 OS=Homo sapiens OX=9606 GN=RPL24 PE=1 SV=1 |
| P35268 | 1 | 21 | 60S ribosomal protein L22 OS=Homo sapiens OX=9606 GN=RPL22 PE=1 SV=2 |
| Q13263 | 1 | 20 | Transcription intermediary factor 1-beta OS=Homo sapiens OX=9606 GN=TRIM28 PE=1 SV=5 |
| Q3KQU3 | 1 | 20 | MAP7 domain-containing protein 1 OS=Homo sapiens OX=9606 GN=MAP7D1 PE=1 SV=1 |
| Q969Q0 | 1 | 20 | 60S ribosomal protein L36a-like OS=Homo sapiens OX=9606 GN=RPL36AL PE=1 SV=3 |
| P23396 | 5 | 96 | 40S ribosomal protein S3 OS=Homo sapiens OX=9606 GN=RPS3 PE=1 SV=2 |
| Q7Z2W4 | 1 | 19 | Zinc finger CCCH-type antiviral protein 1 OS=Homo sapiens OX=9606 GN=ZC3HAV1 PE=1 SV=3 |
| O00571 | 3 | 55 | ATP-dependent RNA helicase DDX3X OS=Homo sapiens OX=9606 GN=DDX3X PE=1 SV=3 |
| P51991 | 2 | 35 | Heterogeneous nuclear ribonucleoprotein A3 OS=Homo sapiens OX=9606 GN=HNRNPA3 PE=1 SV=2 |
| P06748 | 4 | 69 | Nucleophosmin OS=Homo sapiens OX=9606 GN=NPM1 PE=1 SV=2 |
| P61978 | 5 | 84 | Heterogeneous nuclear ribonucleoprotein K OS=Homo sapiens OX=9606 GN=HNRNPK PE=1 SV=1 |
| P62241 | 4 | 66 | 40S ribosomal protein S8 OS=Homo sapiens OX=9606 GN=RPS8 PE=1 SV=2 |
| P18124 | 8 | 125 | 60S ribosomal protein L7 OS=Homo sapiens OX=9606 GN=RPL7 PE=1 SV=1 |
| P62750 | 4 | 62 | 60S ribosomal protein L23a OS=Homo sapiens OX=9606 GN=RPL23A PE=1 SV=1 |
| P18077 | 3 | 46 | 60S ribosomal protein L35a OS=Homo sapiens OX=9606 GN=RPL35A PE=1 SV=2 |
| Q9BYX7 | 2 | 30 | Putative beta-actin-like protein 3 OS=Homo sapiens OX=9606 GN=POTEKP PE=5 SV=1 |
| P26599 | 1 | 15 | Polypyrimidine tract-binding protein 1 OS=Homo sapiens OX=9606 GN=PTBP1 PE=1 SV=1 |
| P46778 | 1 | 15 | 60S ribosomal protein L21 OS=Homo sapiens OX=9606 GN=RPL21 PE=1 SV=2 |
| P62280 | 4 | 59 | 40S ribosomal protein S11 OS=Homo sapiens OX=9606 GN=RPS11 PE=1 SV=3 |
| P05388 | 6 | 88 | 60S acidic ribosomal protein P0 OS=Homo sapiens OX=9606 GN=RPLP0 PE=1 SV=1 |
| P26373 | 7 | 102 | 60S ribosomal protein L13 OS=Homo sapiens OX=9606 GN=RPL13 PE=1 SV=4 |
| Q9NZI8 | 7 | 100 | Insulin-like growth factor 2 mRNA-binding protein 1 OS=Homo sapiens OX=9606 GN=IGF2BP1 PE=1 SV=2 |
| Q9BQ39 | 1 | 14 | ATP-dependent RNA helicase DDX50 OS=Homo sapiens OX=9606 GN=DDX50 PE=1 SV=1 |
| P84103 | 1 | 14 | Serine/arginine-rich splicing factor 3 OS=Homo sapiens OX=9606 GN=SRSF3 PE=1 SV=1 |
| O95793 | 1 | 14 | Double-stranded RNA-binding protein Staufen homolog 1 OS=Homo sapiens OX=9606 GN=STAU1 PE=1 SV=2 |
| P39023 | 6 | 81 | 60S ribosomal protein L3 OS=Homo sapiens OX=9606 GN=RPL3 PE=1 SV=2 |
| P62753 | 2 | 27 | 40S ribosomal protein S6 OS=Homo sapiens OX=9606 GN=RPS6 PE=1 SV=1 |
| P08865 | 4 | 53 | 40S ribosomal protein SA OS=Homo sapiens OX=9606 GN=RPSA PE=1 SV=4 |
| P17844 | 5 | 65 | Probable ATP-dependent RNA helicase DDX5 OS=Homo sapiens OX=9606 GN=DDX5 PE=1 SV=1 |
| P46776 | 2 | 26 | 60S ribosomal protein L27a OS=Homo sapiens OX=9606 GN=RPL27A PE=1 SV=2 |

| | | | |
|---|---|---|---|
| Q9Y6M1 | 1 | 13 | Insulin-like growth factor 2 mRNA-binding protein 2 OS=Homo sapiens OX=9606 GN=IGF2BP2 PE=1 SV=2 |
| P61513 | 1 | 13 | 60S ribosomal protein L37a OS=Homo sapiens OX=9606 GN=RPL37A PE=1 SV=2 |
| Q13151 | 1 | 13 | Heterogeneous nuclear ribonucleoprotein A0 OS=Homo sapiens OX=9606 GN=HNRNPA0 PE=1 SV=1 |
| P52597 | 3 | 38 | Heterogeneous nuclear ribonucleoprotein F OS=Homo sapiens OX=9606 GN=HNRNPF PE=1 SV=3 |
| Q9Y230 | 2 | 25 | RuvB-like 2 OS=Homo sapiens OX=9606 GN=RUVBL2 PE=1 SV=3 |
| P62899 | 3 | 37 | 60S ribosomal protein L31 OS=Homo sapiens OX=9606 GN=RPL31 PE=1 SV=1 |
| Q9NUL3 | 1 | 12 | Double-stranded RNA-binding protein Staufen homolog 2 OS=Homo sapiens OX=9606 GN=STAU2 PE=1 SV=2 |
| Q9UKM9 | 1 | 12 | RNA-binding protein Raly OS=Homo sapiens OX=9606 GN=RALY PE=1 SV=1 |
| P05387 | 1 | 12 | 60S acidic ribosomal protein P2 OS=Homo sapiens OX=9606 GN=RPLP2 PE=1 SV=1 |
| P62913 | 2 | 23 | 60S ribosomal protein L11 OS=Homo sapiens OX=9606 GN=RPL11 PE=1 SV=2 |
| P47914 | 2 | 23 | 60S ribosomal protein L29 OS=Homo sapiens OX=9606 GN=RPL29 PE=1 SV=2 |
| O00425 | 2 | 22 | Insulin-like growth factor 2 mRNA-binding protein 3 OS=Homo sapiens OX=9606 GN=IGF2BP3 PE=1 SV=2 |
| P39019 | 2 | 22 | 40S ribosomal protein S19 OS=Homo sapiens OX=9606 GN=RPS19 PE=1 SV=2 |
| P61353 | 1 | 11 | 60S ribosomal protein L27 OS=Homo sapiens OX=9606 GN=RPL27 PE=1 SV=2 |
| P62424 | 8 | 86 | 60S ribosomal protein L7a OS=Homo sapiens OX=9606 GN=RPL7A PE=1 SV=2 |
| P36578 | 8 | 85 | 60S ribosomal protein L4 OS=Homo sapiens OX=9606 GN=RPL4 PE=1 SV=5 |
| P62829 | 3 | 30 | 60S ribosomal protein L23 OS=Homo sapiens OX=9606 GN=RPL23 PE=1 SV=1 |
| P49207 | 3 | 30 | 60S ribosomal protein L34 OS=Homo sapiens OX=9606 GN=RPL34 PE=1 SV=3 |
| P63173 | 2 | 20 | 60S ribosomal protein L38 OS=Homo sapiens OX=9606 GN=RPL38 PE=1 SV=2 |
| P08621 | 1 | 10 | U1 small nuclear ribonucleoprotein 70 kDa OS=Homo sapiens OX=9606 GN=SNRNP70 PE=1 SV=2 |
| P60891 | 1 | 10 | Ribose-phosphate pyrophosphokinase 1 OS=Homo sapiens OX=9606 GN=PRPS1 PE=1 SV=2 |
| P67809 | 9 | 89 | Nuclease-sensitive element-binding protein 1 OS=Homo sapiens OX=9606 GN=YBX1 PE=1 SV=3 |
| P62269 | 7 | 67 | 40S ribosomal protein S18 OS=Homo sapiens OX=9606 GN=RPS18 PE=1 SV=3 |
| Q92841 | 6 | 57 | Probable ATP-dependent RNA helicase DDX17 OS=Homo sapiens OX=9606 GN=DDX17 PE=1 SV=2 |
| P46782 | 2 | 19 | 40S ribosomal protein S5 OS=Homo sapiens OX=9606 GN=RPS5 PE=1 SV=4 |
| P61247 | 9 | 85 | 40S ribosomal protein S3a OS=Homo sapiens OX=9606 GN=RPS3A PE=1 SV=2 |
| Q15717 | 1 | 9 | ELAV-like protein 1 OS=Homo sapiens OX=9606 GN=ELAVL1 PE=1 SV=2 |
| P52272 | 5 | 44 | Heterogeneous nuclear ribonucleoprotein M OS=Homo sapiens OX=9606 GN=HNRNPM PE=1 SV=3 |
| Q8NC51 | 3 | 26 | Plasminogen activator inhibitor 1 RNA-binding protein OS=Homo sapiens OX=9606 GN=SERBP1 PE=1 SV=2 |
| P38159 | 2 | 17 | RNA-binding motif protein X chromosome OS=Homo sapiens OX=9606 GN=RBMX PE=1 SV=3 |
| P62081 | 2 | 17 | 40S ribosomal protein S7 OS=Homo sapiens OX=9606 GN=RPS7 PE=1 SV=1 |

| | | | |
|---|---|---|---|
| P30050 | 11 | 93 | 60S ribosomal protein L12 OS=Homo sapiens OX=9606 GN=RPL12 PE=1 SV=1 |
| O95816 | 3 | 25 | BAG family molecular chaperone regulator 2 OS=Homo sapiens OX=9606 GN=BAG2 PE=1 SV=1 |
| P27635 | 3 | 25 | 60S ribosomal protein L10 OS=Homo sapiens OX=9606 GN=RPL10 PE=1 SV=4 |
| Q96SB4 | 1 | 8 | SRSF protein kinase 1 OS=Homo sapiens OX=9606 GN=SRPK1 PE=1 SV=2 |
| P25786 | 1 | 8 | Proteasome subunit alpha type-1 OS=Homo sapiens OX=9606 GN=PSMA1 PE=1 SV=1 |
| Q07020 | 10 | 78 | 60S ribosomal protein L18 OS=Homo sapiens OX=9606 GN=RPL18 PE=1 SV=2 |
| Q02878 | 25 | 187 | 60S ribosomal protein L6 OS=Homo sapiens OX=9606 GN=RPL6 PE=1 SV=3 |
| E9PAV3 | 3 | 22 | Nascent polypeptide-associated complex subunit alpha muscle-specific form OS=Homo sapiens OX=9606 GN=NACA PE=1 SV=1 |
| Q13765 | 3 | 22 | Nascent polypeptide-associated complex subunit alpha OS=Homo sapiens OX=9606 GN=NACA PE=1 SV=1 |
| P46779 | 8 | 58 | 60S ribosomal protein L28 OS=Homo sapiens OX=9606 GN=RPL28 PE=1 SV=3 |
| P07900 | 14 | 99 | Heat shock protein HSP 90-alpha OS=Homo sapiens OX=9606 GN=HSP90AA1 PE=1 SV=5 |
| Q13200 | 1 | 7 | 26S proteasome non-ATPase regulatory subunit 2 OS=Homo sapiens OX=9606 GN=PSMD2 PE=1 SV=3 |
| O43143 | 1 | 7 | Pre-mRNA-splicing factor ATP-dependent RNA helicase DHX15 OS=Homo sapiens OX=9606 GN=DHX15 PE=1 SV=2 |
| P60900 | 1 | 7 | Proteasome subunit alpha type-6 OS=Homo sapiens OX=9606 GN=PSMA6 PE=1 SV=1 |
| P09661 | 1 | 7 | U2 small nuclear ribonucleoprotein A' OS=Homo sapiens OX=9606 GN=SNRPA1 PE=1 SV=2 |
| Q07021 | 1 | 7 | Complement component 1 Q subcomponent-binding protein mitochondrial OS=Homo sapiens OX=9606 GN=C1QBP PE=1 SV=1 |
| A0A075B6S2 | 11 | 74 | Immunoglobulin kappa variable 2D-29 OS=Homo sapiens OX=9606 GN=IGKV2D-29 PE=3 SV=1 |
| A0A0A0MRZ7 | 11 | 74 | Immunoglobulin kappa variable 2D-26 OS=Homo sapiens OX=9606 GN=IGKV2D-26 PE=3 SV=1 |
| A2NJV5 | 11 | 74 | Immunoglobulin kappa variable 2-29 OS=Homo sapiens OX=9606 GN=IGKV2-29 PE=3 SV=2 |
| P46783 | 4 | 26 | 40S ribosomal protein S10 OS=Homo sapiens OX=9606 GN=RPS10 PE=1 SV=1 |
| P61313 | 7 | 43 | 60S ribosomal protein L15 OS=Homo sapiens OX=9606 GN=RPL15 PE=1 SV=2 |
| P14625 | 3 | 18 | Endoplasmin OS=Homo sapiens OX=9606 GN=HSP90B1 PE=1 SV=1 |
| Q86UE4 | 1 | 6 | Protein LYRIC OS=Homo sapiens OX=9606 GN=MTDH PE=1 SV=2 |
| O43242 | 1 | 6 | 26S proteasome non-ATPase regulatory subunit 3 OS=Homo sapiens OX=9606 GN=PSMD3 PE=1 SV=2 |
| Q9Y281 | 1 | 6 | Cofilin-2 OS=Homo sapiens OX=9606 GN=CFL2 PE=1 SV=1 |
| P23528 | 1 | 6 | Cofilin-1 OS=Homo sapiens OX=9606 GN=CFL1 PE=1 SV=3 |
| P40429 | 12 | 68 | 60S ribosomal protein L13a OS=Homo sapiens OX=9606 GN=RPL13A PE=1 SV=2 |
| Q9Y265 | 6 | 34 | RuvB-like 1 OS=Homo sapiens OX=9606 GN=RUVBL1 PE=1 SV=1 |
| Q02543 | 11 | 62 | 60S ribosomal protein L18a OS=Homo sapiens OX=9606 GN=RPL18A PE=1 SV=2 |
